# Altered stereostructures of the DNA-binding domains in the mutant mating proteins of *Ophiocordyceps sinensis* and the *Cordyceps sinensis* insect‒fungal complex

**DOI:** 10.1101/2025.06.26.661214

**Authors:** Xiu-Zhang Li, Yu-Ling Li, Jia-Shi Zhu

## Abstract

The MATα_HMGbox and HMG-box_ROX1-like domains of the MAT1-1-1 and MAT1-2-1 proteins, respectively, play essential roles in DNA binding and the downstream regulation of gene transcription that controls sexual reproduction in *Ophiocordyceps sinensis*. Alternative splicing, differential occurrence and transcription of the *MAT1-1-1* and *MAT1-2-1* genes have been revealed in *Hirsutella sinensis* (Genotype #1 of the 17 *O. sinensis* genotypes), suggesting the occurrence of self-sterility under heterothallic or hybrid outcrossing. This study demonstrates that the MATα_HMGbox domains of MAT1-1-1 proteins in wild-type *Cordyceps sinensis* isolates cluster into 5 clades with or without branches in the Bayesian clustering tree and belong to diverse 3D structural morphs under 19 AlphaFold codes. The HMG-box_ROX1-like domains of MAT1-2-1 proteins cluster into 2 clades with branches in the Bayesian clustering tree and belong to diverse 3D structural morphs under 25 AlphaFold codes. Correlation analysis reveals that 1−3 amino acid substitutions at various sites in the DNA-binding domains of the mutant mating proteins result in alterations in the hydrophobicity and the secondary and tertiary structures of the DNA-binding domains of the proteins. Fungal origin analysis reveals heterospecific fungal sources or genome-independent, genotypic sources of the mating proteins with stereostructure variations in the wild-type *C. sinensis* isolates that represent the changed functionalities of the proteins in regulating and ensuring the accuracy and genetic diversity of the heterothallic and hybrid sexual reproductive processes of *O. sinensis* during the lifecycle of the *C. sinensis* insect‒fungi complex.

## I. Introduction

Natural *Cordyceps sinensis* is one of most expensive therapeutic agents in traditional Chinese medicine and has a rich history of clinical use for several centuries for health maintenance, disease amelioration, post-illness and postoperative recovery, and antiaging therapy [Zhu *et al*. 1998a, 1998b, 2011]. This prestige natural therapeutic agent consists of the *Ophiocordyceps sinensis* fruiting body and the remains of a *Hepialidae* moth larva containing an intact, thick larval body wall with numerous bristles, an intact larval intestine, head tissues, and fragments of other larval tissues [Ren *et al*. 2013; Zhang *et al*. 2014; Lu *et al*. 2016; Zhu & Li 2017; Li *et al*. 2022, 2023a, 2023d, 2023e]. Studies of natural *C. sinensis* have demonstrated its multicellular heterokaryotic structures and genetic heterogeneity; the latter includes 17 genomically independent genotypes of *O. sinensis* fungi, >90 other fungal species spanning at least 37 fungal genera, and larval genes [Li 1988; Dai *et al*. 1989; Kinjo & Zang 2001; Jiang & Yao 2003; Stensrud *et al*. 2005, 2007; Leung *et al*. 2006; Zhu *et al*. 2007, 2010; Yang *et al*. 2008, 2021; Zhang *et al*. 2010, 2018; Li *et al*. 2013, 2016b, 2020b, 2022, 2023e; Ren *et al*. 2013; Meng *et al*. 2015; Xia *et al*. 2015; Guo *et al*. 2017; Zhu & Li 2017; Zhong *et al*. 2018; Kang *et al*. 2024]. Thus, the Chinese Pharmacopoeia defines natural *C. sinensis* as an insect‒fungal complex classified as a LEVEL-II endangered natural species [China Ministry of Agriculture and Rural Affairs 2021]. Among the numerous heterogeneous fungal species [Jiang & Yao 2003; Zhang *et al*. 2010; Xia *et al*. 2015], *Hirsutella sinensis* has been postulated to be the sole anamorph of *O. sinensis* by Wei *et al*. [2006]. However, ten years later in an artificial cultivation study conducted in an industrial product-oriented setting, Wei *et al*. [2016] reported a species contradiction between anamorphic inoculates of GC-biased Genotype #1 *H. sinensis* on *Hepialidae* moth larvae and the sole teleomorph (the genomically independent AT-biased Genotype #4 of *O. sinensis*) in the fruiting body of cultivated *C. sinensis*. Notably, the Latin name *Cordyceps sinensis* has been indiscriminately used since the 1840s to refer to both the teleomorph/holomorph of the fungus *C. sinensis* and the wild insect‒fungal complex; the fungus was renamed *O. sinensis* in 2007 with use of the *H. sinensis* strain EFCC 7287 as the nomenclature reference [Sung *et al*. 2007; Zhang *et al*. 2012; Ren *et al*. 2013; Zhu & Li 2017; Wang *et al*. 2018; Li *et al*. 2022]. Although Zhang *et al*. [2013] proposed improper implementation of the “One Fungus=One Name” nomenclature rule set by the International Mycological Association [Hawksworth *et al*. 2011], disregarded the presence of multiple genomically independent genotypes of *O. sinensis* fungi and inappropriately replaced the anamorphic name *H. sinensis* with the teleomorphic name *O. sinensis*, in this paper, we continue to use the anamorphic name *H. sinensis* for GC-biased Genotype #1 of the 17 *O. sinensis* genotypes that may share the same evolutionary ancestor [Stensrud *et al*. 2007] and refer to the genomically independent Genotypes #2‒17 fungi as *O. sinensis* following the taxonomic descriptions that are used in several public depository databases, such as the GenBank and AlphaFold databases, before the systematic positions of these evolutionarily related genotypes are individually determined and differentiated, regardless of whether they are GC- or AT-biased genetically. In this study, we continue the customary use of the name *C. sinensis* to refer to the wild or cultivated insect‒fungal complex because the renaming of *C. sinensis* to *O. sinensis* in 2007 using the *H. sinensis* strain EFCC 7287 as the nomenclature reference did not involve the indiscriminately used Latin name for the natural insect‒fungal complex [Sung *et al*. 2007; Yao & Zhu 2016; Zhu & Li 2017; Li *et al*. 2022]. However, this practice will likely be revised in the future by the differential use of proprietary and exclusive Latin names for the multiple genome‒independent *O. sinensis* genotypic fungi and the insect‒fungal complex.

The sexual reproductive behavior of ascomycetes is strictly regulated by transcription factors encoded at the mating-type (*MAT*) locus. These transcription factors constitute the core mechanism that determines mating compatibility, regulates mating type recognition, and controls the development of fruiting bodies, reproductive structure ascocarps and ascospores [Turgeon & Yoder 2006; Debuchy *et al*. 2006; Jones & Bennett 2011; Zheng & Wang 2013; Wilson *et al*. 2015; Sun *et al*. 2019; Ramšak *et al*. 2020, 2021; Ramšak & Kück 2022]. Like other fungi in Ascomycota, the reproductive behavior of *O. sinensis* relies on the synergistic interaction of 2 mating proteins, the *MAT1-1* and *MAT1-2* idiomorphs. These proteins contain 2 types of critical domains that specifically regulate gene expression related to sexual reproduction: (1) the mating-type alpha high mobility group box (MATα_HMGbox) domain in the MAT1-1-1 protein and (2) the high mobility group box ROX1-like (HMG-box_ROX1-like) domain in the MAT1-2-1 protein [Metin *et al*. 2010; Bushley *et al*. 2013; Hu *et al*. 2013; Zheng & Wang 2013; Li *et al*. 2025]. However, alternative splicing, differential occurrence, and differential transcription of mating-type and pheromone receptor genes and heteromorphic stereostructures of the entire MAT1-1-1 and MAT1-2-1 proteins in *H. sinensis* strains and wild-type *C. sinensis* isolates have been demonstrated, invalidating the self-fertilization hypothesis under homothallism and pseudohomothallism for *H. sinensis*, which was postulated as the sole anamorph of *O. sinensis* and instead suggesting that self-sterility under heterothallism or hybridization requires sexual partners to accomplish sexual reproduction within the lifecycle of natural *C. sinensis* insect−fungi complex [Wei *et al*. 2006; Zhou *et al*. 2013; Zhang & Zhang 2015; Li *et al*. 2019, 2023c, 2024b, 2025].

Mutations of the 2 critical DNA-binding domains of mating proteins that result in a few amino acid substitutions may cause conformational changes in the core stereostructures of the domains, thereby affecting the synergistic interaction of the proteins, probably through regulation of the expression of complementary downstream genes of the 2 idiomorphic mating proteins. Continuing our previous study [Li *et al*. 2025], which demonstrated the variations in the 3D structures of complete mating proteins, this study focuses on amino acid substitutions within the MATα_HMGbox domain and the HMG-box_ROX1-like domain of the full-length MAT1-1-1 and MAT1-2-1 proteins, respectively, and correlates the variable primary structures of these domains with changes in the hydrophobic properties and the secondary and tertiary structures of the DNA-binding domains in wild-type *C. sinensis* isolates. Correlations between changes in hydrophobicity and the primary and secondary structures of the DNA-binding domains of mating proteins encoded by the genome, transcriptome and metatranscriptome assemblies of *H. sinensis* and C*. sinensis* insect‒fungal complexes were also analyzed.

## II. Materials and methods

### II-1 The MAT1-1-1 and MAT1-2-1 protein sequences of the wild-type *C. sinensis* isolates

The AlphaFold database lists the AlphaFold-predicted 3D structures for 138 MAT1-1-1 proteins and 74 MAT1-2-1 proteins [Li *et al*. 2025]. Of these proteins, 118 MAT1-1-1 proteins and 69 MAT1-2-1 proteins are full-length proteins, as shown in Tables S1−S2. Three full-length MAT1-1-1 proteins (ALH24945, AGW27560, and EQK97643) and 2 full-length MAT1-2-1 proteins (AEH27625 and EQL04085) are derived from *H. sinensis* strains. All other full-length mating proteins are from wild-type *C. sinensis* isolates that were collected from various production areas on the Qinghai‒Tibet Plateau [Zhang *et al*. 2009, 2011, 2012, 2013, 2014; Hu *et al*. 2013; Zhang & Zhang 2015; Tunyasuvunakool *et al*. 2021; Li *et al*. 2025]. The internal transcribed spacer (ITS) genotypic information is available in the GenBank database for the mutant full-length MAT1-1-1 and MAT1-2-1 proteins that are derived from 34 wild-type *C. sinensis* isolates and aligned with the reference ITS sequences of the 17 genotypes of *O. sinensis* (*cf*. Table S3).

The remaining 20 MAT1-1-1 proteins (AGW27517−AGW27536) and 5 MAT1-2-1 proteins (AGW27543, AGW27548, AGW27552, AGW27554, and AGW27555) are truncated and will be analyzed elsewhere [Bushley *et al*. 2013; Li *et al*. 2013].

### II-2 Genome, transcriptome, and metatranscriptome assemblies of *H. sinensis* strains and *C. sinensis* insect−fungal complexes

The GenBank database lists 5 genome assemblies, LKHE00000000, NGJJ00000000, ANOV00000000, JAAVMX000000000, and LWBQ00000000, of the *H. sinensis* strains 1229, CC1406-20395, Co18, IOZ07, and ZJB12195, respectively, and a transcriptome assembly, GCQL00000000, of the *H. sinensis* strain L0106 [Hu *et al*. 2013; Liu *et al*. 2015, 2020; Li *et al*. 2016a; Jin *et al*. 2020; Shu *et al*. 2020].

The metatranscriptome assembly GAGW00000000 for the *C. sinensis* insect‒fungal samples (unknown maturation stages) collected from Kangding County, Sichuan Province, China [Xiang *et al*. 2014] is available in the GenBank database.

Another metatranscriptome assembly was derived from mature *C. sinensis* samples collected from Deqin, Yunnan Province, China, and was uploaded to a repository database, www.plantkingdomgdb.com/Ophiocordyceps_sinensis/data/cds/Ophiocordyceps_sinensis_CDS.fas [Xia *et al*. 2017]. Although this database is currently inaccessible, it was accessed from 18 May 2017 to 18 January 2018, and a cDNA file was downloaded.

The genome, transcriptome, and metatranscriptome assemblies were used to analyze the MATα_HMGbox and HMG-box_ROX1-like domains of the MAT1-1-1 and MAT1-2-1 proteins, respectively.

### II-3 Alignment of the DNA-binding domain sequences of the mating proteins

The amino acid sequences of the MATα_HMGbox domains of the MAT1-1-1 proteins and the HMG-box_ROX1-like domains of the MAT1-2-1 proteins of the wild-type *C. sinensis* isolates and *C. sinensis* insect‒fungal complexes were aligned using the GenBank Blastp program (https://blast.ncbi.nlm.nih.gov/ (Bethesda, MD, USA), accessed from 18 October 2024 to 10 July 2025).

### II-4 Bayesian clustering analysis of the DNA-binding domains of the mating proteins

Multiple sequence alignment of the DNA-binding domains of the mating proteins of the wild-type *C. sinensis* isolates and *C. sinensis* insect‒fungal complexes was performed via the auto mode of MAFFT (v7.427; Osaka, Japan). The intron regions were removed using trimAl (v1.4. rev15; Barcelona, Spain). Under the Bayesian information criterion, ProtTest v3.4.2 (University of Vigo and University of A Coruña, Spain) was used to find the optimal amino acid substitution model. Bayesian clustering trees of the DNA binding domain sequences of the MAT1-1-1 and MAT1-2-1 proteins were then inferred using MrBayes v3.2.7 software (Markov chain Monte Carlo [MCMC] algorithm; Uppsala, Sweden) at a sampling frequency of 100 iterations after discarding the initial 25% of the samples from a total of 1 million iterations. This yielded final majority rule consensus trees for the MATα_HMGbox domains of the full-length MAT1-1-1 proteins and the HMG-box_ROX1-like domains of the full-length MAT1-2-1 proteins [Huelsenbeck & Ronquist 2001; Ronquist *et al*. 2012; Li *et al*. 2022, 2023c, 2024b, 2025]. Clustering analysis was conducted at Nanjing Genepioneer Biotechnologies Co. (Nanjing, China).

### II-5 Amino acid properties and scale analysis

The amino acid sequences of the DNA-binding domains of the mating proteins were scaled on the basis of the general chemical characteristics of their side chains (Table S4) as reported at https://web.expasy.org/protscale/ (Basel, Switzerland), accessed from 18 October 2024 to 20 May 2025 [Kyte & Doolittle 1982; Deleage & Roux 1987; Gasteiger *et al*. 2005; Peters & Elofsson 2014; Simm *et al*. 2016; Li *et al*. 2024b, 2025]. The MATα_HMGbox domain of MAT1-1 proteins (amino acids 51→225 of the reference sequence AGW27560 derived from the *H. sinensis* strain CS68-2-1229 [Bushley *et al*. 2013]) and the HMG-box_ROX1-like domain of MAT1-2-1 proteins (amino acids 127→197 of the reference sequence AEH27625 derived from the *H. sinensis* strain CS2 [Zhang *et al*. 2009]), with additional 9 amino acid residues extending upstream and downstream of the domains, were plotted sequentially with a window size of 9 amino acid residues using the linear weight variation model of the ExPASy ProtScale algorithms [Kyte & Doolittle 1982; Deleage & Roux 1987; Gasteiger *et al*. 2005; Li *et al*. 2024b, 2025] to generate ExPASy ProtScale plots to display and compare the topology and waveform changes in hydrophobicity and the 2D structures for α-helices, β-sheets, β-turns, and coils of the DNA-binding domains of the mating proteins.

### II-6 AlphaFold-based prediction of the 3D structures of the mating proteins

For heteromorphic stereostructure analysis in this study, the 3D structures of the MAT1-1-1 and MAT1-2-1 proteins of the wild-type *C. sinensis* isolates were computationally predicted from their amino acid sequences using the artificial intelligence (AI)-based machine learning technology AlphaFold (https://alphafold.com/, Cambridgeshire, UK), downloaded from the AlphaFold database (accessed from 18 October 2024 to 10 May 2025) [Jumper *et al*. 2021; David *et al*. 2022; Monzon *et al*. 2022; Rettie *et al*. 2023; Xu *et al*. 2023; Abramson *et al*. 2024; Varadi *et al*. 2024; Wroblewski & Kmiecik 2024; Li *et al*. 2025].

The AlphaFold database provides per-residue model confidence, predicts scores between 0 and 100 on the local distance difference test (pLDDT) and provides a per-residue score that is assigned to each individual residue [Jumper *et al*. 2021; David *et al*. 2022; Monzon *et al*. 2022; Xu *et al*. 2023; Abramson *et al*. 2024; Varadi *et al*. 2024]. The model confidence bands are used to color-code the residues in the 3D structures: residues with very high confidence (pLDDT>90) are shown in dark blue, those with high confidence (90>pLDDT>70) appear in light blue, residues with low confidence (70>pLDDT>50) are shown in yellow, and residues with very low confidence (pLDDT<50) are shown in orange [Mariani *et al*. 2013; Wroblewski & Kmiecik 2024; Li *et al*. 2025]. The AlphaFold database provides an average pLDDT score for each of the AI-predicted 3D structural models.

### II-7 Correlation of the primary, secondary, and tertiary structures of the DNA-binding domains of the MAT1-1-1 and MAT1-2-1 proteins

The AlphaFold-predicted 3D structures of the MATα_HMGbox domains of the MAT1-1-1 proteins and the HMG-box_ROX1-like domains of the MAT1-2-1 proteins were amplified at the sites of amino acid substitutions. The locally magnified 3D structures at the mutation sites were correlated with the amino acid substitutions to obtain the primary structures and the changes in topology and waveform using ExPASy ProtScale plotting showing the hydropathy, α-helices, β-sheets, β-turns, and coils for altered hydrophobicity and the 2D structures of the DNA binding domains of the MAT1-1-1 and MAT1-2-1 proteins.

## III. Results

### III-1 Primary structures of the MATα**_**HMGbox domains of the full-length MAT1-1-1 proteins

Of the 138 MAT1-1-1 proteins in the AlphaFold database, 118 (85.5%) are full length. Each of these proteins contains 372 amino acid residues. These proteins contribute to 15 heteromorphic 3D structure morphs belonging to 5 Bayesian clusters (Table S1) [Li *et al*. 2025]. Twenty-nine (24.6%) of the 118 full-length MAT1-1-1 proteins derived from wild-type *C. sinensis* isolates contain variable amino acid substitutions.

The MAT1-1-1 proteins contain a MATα_HMGbox domain that locates at amino acids 51→225 (shown in blue and underlined in Figure S1), as represented by the reference MAT1-1-1 protein AGW27560 derived from the *H. sinensis* strain CS68-2-1229 [Bushley *et al*. 2013]. Other MAT1-1-1 proteins derived from wild-type *C. sinensis* isolates or encoded by the genome assemblies of the *H. sinensis* strains and the metatranscriptome assemblies of *C. sinensis* insect‒fungal complexes contain various amino acid substitutions, as shown in red in Figure S1. The MATα_HMGbox domains of 10 (34.5%) of the 29 mutant proteins are 100% identical to that of the query protein AGW27560. Each of the remaining 19 proteins (65.5%) contains 1 or 2 substitutions in the amino acid residues at various sites within the MATα_HMGbox domains.

Figure S1 also shows the MATα_HMGbox domains of the MAT1-1-1 proteins encoded by the genome assemblies ANOV01017390 (794←1129 & 1280←1396), LKHE01001116 (4183←4620 & 4667←4759), and JAAVMX010000001 (6,699,061→6,699,153 & 6,699,203→6,699.637) of the *H. sinensis* strains Co18, 1229, and IOZ07, respectively [Hu *et al*. 2013; Li *et al*. 2016a; Shu *et al*. 2020]. (Note: the arrows “→” and “←” indicate sequences in the sense and antisense strands of the genomes, respectively; “&” refers to the removed intron portion.) These genome-encoded MAT1-1-1 proteins are truncated at their C-termini; the truncation segments are located far downstream of the MATα_HMGbox domains.

**Figure 1.**
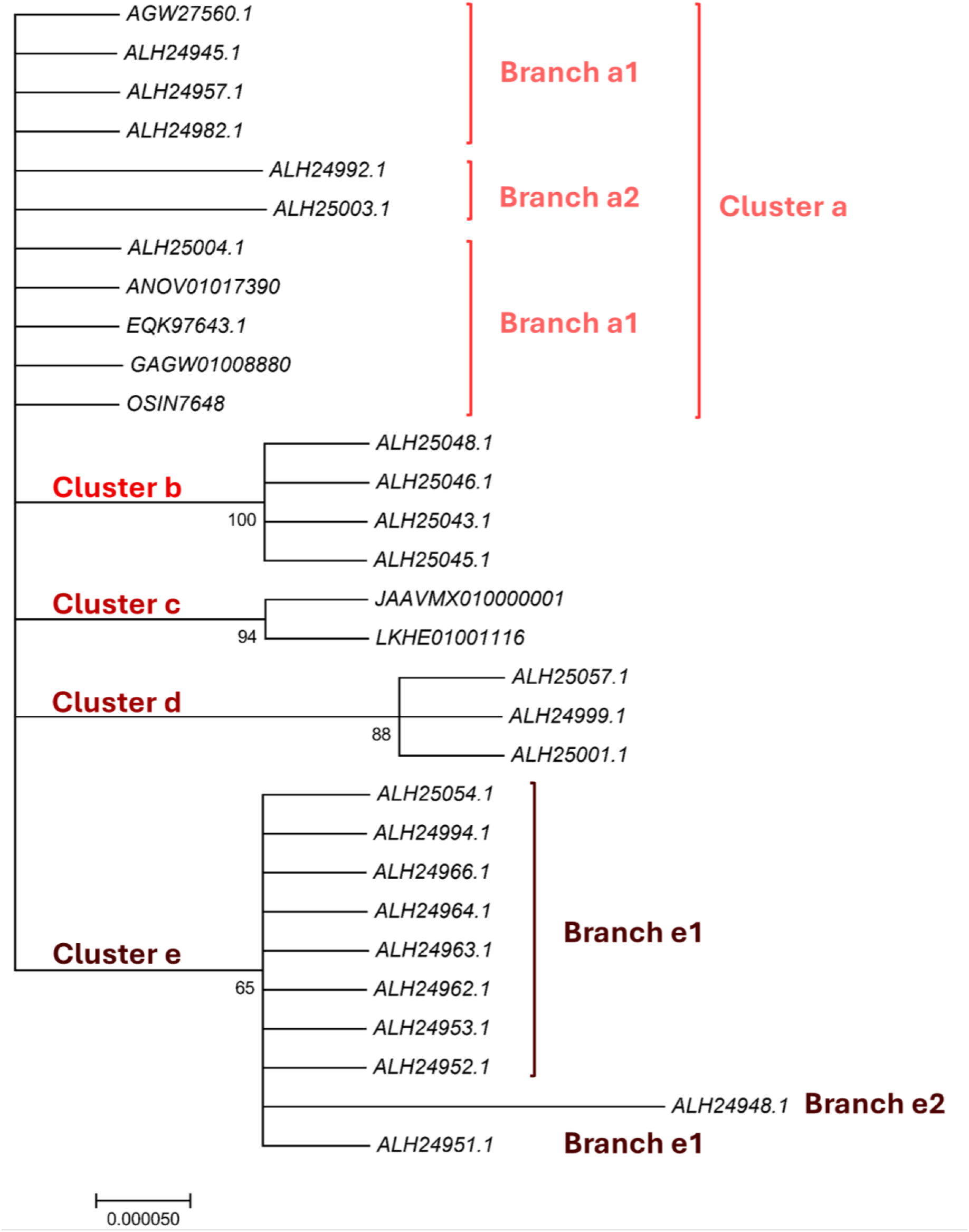
A Bayesian majority rule consensus clustering tree for the MATα_HMGbox domain sequences of the 25 MAT1-1-1 proteins of the wild-type *C. sinensis* isolates and the corresponding domain sequences encoded by the genome and metatranscriptome assemblies of the *H. sinensis* strains and *C. sinensis* insect‒fungal complexes was obtained by MrBayes v3.2.7 software.

The genome assemblies LWBQ00000000 and NGJJ00000000 and the transcriptome assembly GCQL00000000 of the *H. sinensis* strains ZJB12195, CC1406-20395, and L0106, respectively, do not contain the genes or transcripts that encode the MAT1-1-1 proteins [Jin *et al*. 2020; Liu *et al*. 2015, 2020].

The metatranscriptome assemblies GAGW01008880 (300←1127) and OSIN7648 (1→1065) of natural *C. sinensis* insect‒fungal complexes encode the MAT1-1-1 proteins (*cf*. Figure S1) [Xiang *et al*. 2014; Xia *et al*. 2017]. The MAT1-1-1 protein OSIN7648 is truncated in midsequence by 18 amino acid residues (SMQREYQAPRFFYDYSVS) between residues 863 and 864; the truncation segment is located downstream of the MATα_HMGbox domain (151→675). The N-terminus of the MATα_HMGbox domain (741←1127) of the MAT1-1-1 protein encoded by GAGW01008880 is truncated by 46 amino acid residues (AAASRATRQTKEASCDRAKRPLNAFMAFRSYYLKLFPDVQQKTASG).

### III-2 Primary structures of the HMG-box_ROX1-like domains of the full-length MAT1-2-1 proteins

Of the 74 MAT1-2-1 proteins listed in the AlphaFold database, 69 (93.2%) are full-length proteins, each of which contains 249 amino acid residues; these proteins contribute to 17 heteromorphic AlphaFold tertiary structure morphs belonging to 5 Bayesian clusters (Table S2) [Li *et al*. 2025]. Of the 69 full-length proteins, 39 (56.5%) are 100% identical to the representative MAT1-2-1 protein AEH27625, which was derived from the *H. sinensis* strain CS2 [Zhang *et al*. 2009]. The remaining 30 full-length proteins (43.5%) contain amino acid substitutions at various sites, as shown in red in Figure S2. The HMG-box_ROX1-like domains of the MAT1-2-1 proteins are located at amino acid residues 127→197, as shown in blue and underlined in Figure S2; whereas the 9 external amino acid residues upstream and downstream of the domain are shown blue but not underlined. The HMG-box_ROX1-like domain sequences of 3 of the 30 variable AMT1-2-1 proteins are 100% identical to the domain sequence of the query protein AEH27625, whereas the domains of the remaining 27 proteins contain 1−3 amino acid substitutions, as shown in red in Figure S2.

Figure S2 also shows the MAT1-2-1 proteins encoded by the genome assemblies ANOV01000063, LKHE01001605, LWBQ01000021, and NGJJ01000619 of the *H. sinensis* strains Co18, 1229, ZJB12195, and CC1406-20395, respectively [Hu *et al*. 2013; Li *et al*. 2016a; Jin *et al*. 2020; Liu *et al*. 2020]. An S-to-A substitution occurred in each of the HMG-box_ROX1-like domains of the genome-encoded MAT1-2-1 proteins ANOV01000063 (9759→9851 & 9907→10,026), LKHE01001605 (14,016←14,135 & 14,191←14,283), LWBQ01000021 (239,029←239,148 & 239,204←239,269), and NGJJ01000619 (23,186←23,305 & 23,361←23,453). An additional Y-to-H substitution occurred in the HMG-box_ROX1-like domains of LKHE01001605, LWBQ01000021 and NGJJ01000619 but not in the HMG-box_ROX1-like domain of ANOV01000063, resulting in 97.2−98.6% similarity to the sequence of the HMG-box_ROX1-like domain of the query protein AEH27625 [Zhang *et al*. 2009].

**Figure 2.**
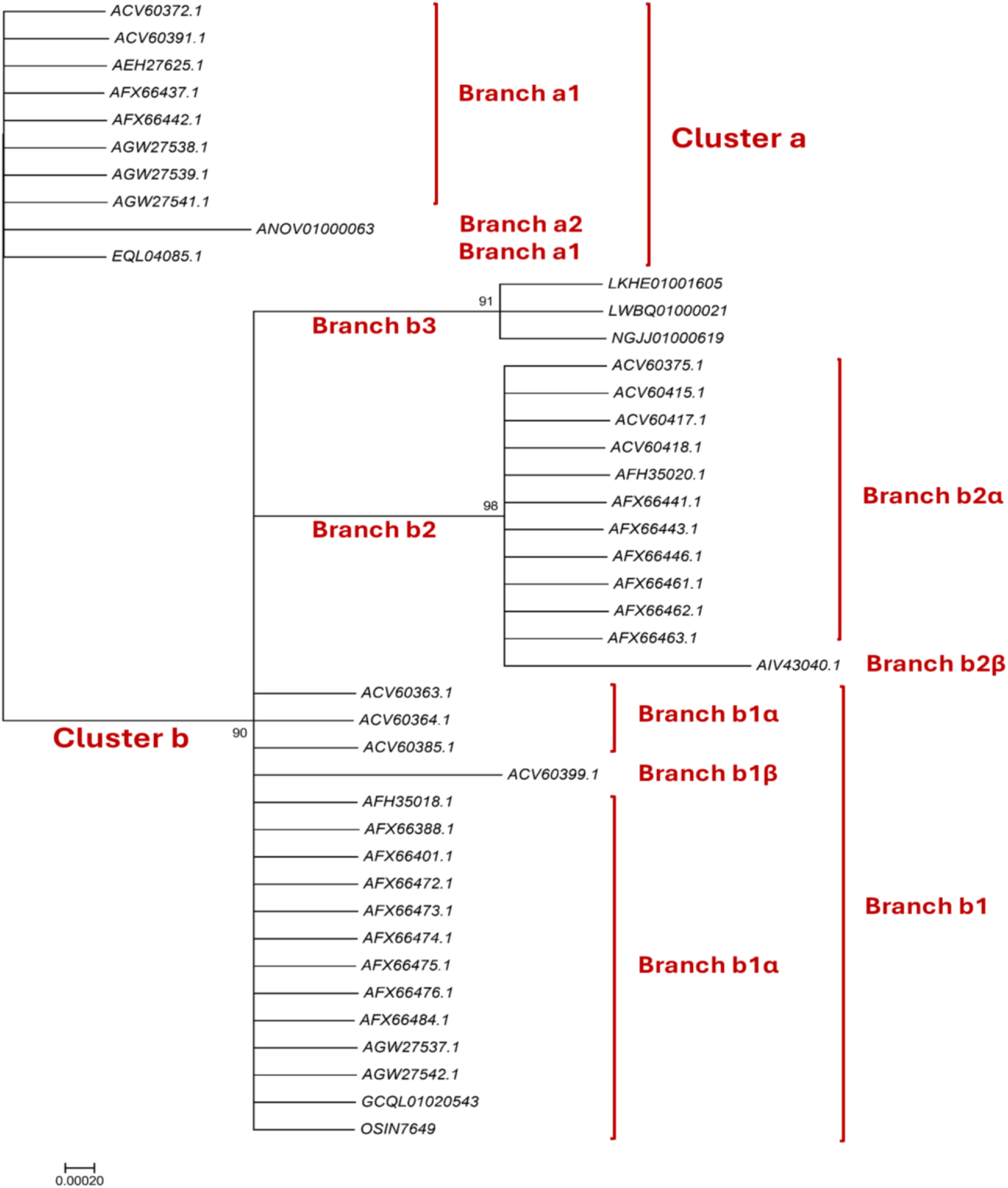
Bayesian majority rule consensus clustering tree inferred via MrBayes v3.2.7 software for the HMG-box_ROX1-like domain sequences of 35 full-length MAT1-2-1 proteins of the wild-type *C. sinensis* isolates and the corresponding domain segments of the translated genome, transcriptome, and metatranscriptome assemblies of the *H. sinensis* strains and *C. sinensis* insect‒fungal complexes.

The genome assembly JAAVMX000000000 of the *H. sinensis* strain IOZ07 and the metatranscriptome assembly GAGW00000000 of the *C. sinensis* insect–fungi complex do not contain genes or transcripts encoding MAT1-2-1 proteins [Xiang *et al*. 2014; Shu *et al*. 2020].

Figure S2 also shows the protein sequences encoded by the MAT1-2-1 transcripts of the transcriptome assembly GCQL01020543 (397←1143) of the *H. sinensis* strain L0106 and the metatranscriptome assembly OSIN7649 (1→747) of the mature *C. sinensis* insect‒fungal complex [Liu *et al*. 2015; Xia *et al*. 2017]. The HMG-box_ROX1-like domains (553←765 of GCQL01020543 and 379→591 of OSIN7649) of these proteins contain Y-to-H substitutions, resulting in 98.6% similarity to the sequence of that domain in the query protein AEH27625 [Zhang *et al*. 2013].

### III-3 Bayesian analysis of the MATα_HMGbox domains of the full-length MAT1-1-1 proteins

The sequences of the MATα_HMGbox domains of 19 full-length MAT1-1-1 proteins that display various amino acid substitutions were subjected to Bayesian clustering analysis and compared with the sequences of 6 authentic MAT1-1-1 proteins as the references (Figure 1).

The MATα_HMGbox domains of the full-length MAT1-1-1 proteins were clustered into 5 Bayesian clusters (Figure 1), among which Clusters a and d were branched. The domains of the 19 proteins with various amino acid substitutions were clustered into Clusters/Branches a2, b, d, and e1−e2 in the Bayesian clustering tree. Cluster d has a longer clustering distance than some of the other clusters do, and Branch e2 has the longest clustering distance. The MATα_HMGbox domains encoded by the genome assemblies JAAVMX010000001 (6,699,061→6,699,153 & 6,699,203→6,699.637) and LKHE01001116 (4183←4620 & 4667←4759) of the *H. sinensis* strains 1229 and IOZ07 contain Y-to-M substitutions [Li *et al*. 2016; Shu *et al*. 2020]. The domains encoded by the genome assembly ANOV01017390 (794←1129 & 1280←1396) of *H. sinensis* strain Co18 and the metatranscriptome assembly OSIN7648 (151→675) of the *C. sinensis* insect‒fungal complex contain no changed amino acid residues [Hu *et al*. 2013; Xia *et al*. 2017]. The domain encoded by the metatranscriptome assembly GAGW01008880 (714←1127) of the *C. sinensis* insect‒fungal complex is truncated at the N-terminus (Figure S1) [Xiang *et al*. 2014]. The MATα_HMGbox domains encoded by the genome assembly ANOV01017390 and metatranscriptome assemblies OSIN7648 and GAGW01008880 cluster into Branch a1 in the Bayesian clustering tree, together with the 6 authentic MAT1-1-1 proteins (Figure 1).

Tables S5−S6 summarize the Bayesian clustering results and show the amino acid substitutions and deletions in the MATα_HMGbox domains of the MAT1-1-1 proteins derived from wild-type *C. sinensis* isolates and those encoded by the genome and metatranscriptome assemblies of *H. sinensis* strains and natural *C. sinensis* insect‒fungal complexes, respectively.

### III-4 Bayesian analysis of the HMG-box_ROX1-like domains of the full-length MAT1-2-1 protein sequences

Thirty full-length MAT1-2-1 proteins that are present in wild-type *C. sinensis* isolates or encoded by the genome and transcriptome assemblies of *H. sinensis* strains and by the metatranscriptome assembly of the *C. sinensis* insect‒fungal complex and that have various amino acid substitutions in their HMG-box_ROX1-like domains were subjected to Bayesian clustering analysis and compared with the HMG-box_ROX1-like domains of 6 authentic MAT1-2-1 proteins as the references. Figure 2 shows 2 branched clusters (a and b). Branches b2−b3 have longer clustering distances than do some of the other branches, and Branch b2β has the longest clustering distance.

The HMG-box_ROX1-like domain encoded by the genome assembly ANOV01000063 (9759→9851 & 9907→10,026) of *H. sinensis* strain Co18 contains an S-to-A substitution and is clustered into Branch a2 with a slightly longer clustering distance (Figures 2 and S2) [Hu *et al*. 2013]. The domains encoded by other genome assemblies [LKHE01001605 (14,016←14,135 & 14,191←14,135 & 14,191←14,283), LWBQ01000021 (239,029←239,148 & 239,204←239,269), and NGJJ01000619 (23,186←23,305 & 23,361←23,453) of the *H. sinensis* strains 1229, ZJB12195, and CC1406-20395, respectively] [Li *et al*. 2016; Jin *et al*. 2020; Liu *et al*. 2020] contain Y-to-H and S-to-A substitutions and are clustered into Branch b3 in the Bayesian clustering tree (Figures 2 and S2).

The HMG-box_ROX1-like domain of the MAT1-2-1 protein encoded by the transcriptome assembly GCQL01020543 (553←765) of the *H. sinensis* strain L0106 contains a Y-to-H substitution [Liu *et al*. 2015]. The domain of the MAT1-2-1 protein encoded by the metatranscriptome assembly OSIN7649 (379→591) of the *C. sinensis* insect‒fungi complex is 100% identical to the reference MAT1-2-1 protein AEH27625 [Zhang *et al*. 2009; Xia *et al*. 2017]. The domains of the MAT1-2-1 proteins encoded by the transcriptome and metatranscriptome assemblies were clustered into Branch b1α in the Bayesian clustering tree (Figures 2 and S2).

Tables S7−S8 summarize the Bayesian clustering results that reflect the amino acid substitutions and deletions in the HMG-box_ROX1-like domains of MAT1-2-1 proteins derived from wild-type *C. sinensis* isolates or encoded by the genome, transcriptome and metatranscriptome assemblies of *H. sinensis* strains and natural *C. sinensis* insect‒fungal complexes, respectively.

### III-5 Heteromorphic stereostructures of the MATα_HMGbox domains of full-length MAT1-1-1 proteins derived from wild-type *C. sinensis* isolates

Section III-5 presents analyses of the correlations between the changes in the structure and hydrophobicity of the MATα_HMGbox domains of full-length MAT1-1-1 proteins (under 7 AlphaFold 3D structure codes; *cf*. Table S5) derived from wild-type *C. sinensis* isolates, compared to the reference MAT1-1-1 proteins represented by AGW27560 (under the AlphaFold code U3N942) derived from the *H. sinensis* strain CS68-2-1229 (*cf*. Table S1) [Bushley *et al*. 2013]. The MATα_HMGbox domain sequences (amino acid residues 51→225) were extended by 9 amino acid residues both upstream and downstream of the domains because of the setting of the sequential ExPASy ProtScale plotting for hydropathy, α-helices, β-sheets, β-turns, and coils at a window size of 9 amino acid residues (*cf*. Section II-5 of Materials and methods).

Figure 3 compares the hydrophobicities and the structures of the MATα_HMGbox domain of the mutant MAT1-1-1 protein ALH24992 (under the AlphaFold code A0A0N9R5B3) derived from the wild-type *C. sinensis* isolate SC09_65 (*cf*. Tables S1 and S5) [Zhang & Zhang 2015] with those of the MATα_HMGbox domain of the reference protein AGW27560 (under the AlphaFold code U3N942) derived from the *H. sinensis* strain CS68-2-1229 [Bushley *et al*. 2013]. Panel A shows an A-to-T substitution (aliphatic alanine to threonine with polar side chains; hydropathy indices changed from 1.8 to −0.7). This substitution causes reduced hydrophobicity (the larger the hydropathy value is the stronger the hydrophobicity, whereas negative values indicate hydrophilicity [Kyte & Doolittle 1982]), as illustrated by the changes in topology and waveform in the hydropathy plot shown in Panel B. These changes result in an altered secondary structure of the protein in the area surrounding the mutation site in the MATα_HMGbox domain of the mutant MAT1-1-1 protein ALH24992, as illustrated by the changes in topology and waveform in the ExPASy ProtScale plots for α-helices, β-sheets, β-turns, and coils shown in Panels C−F, respectively.

**Figure 3.**
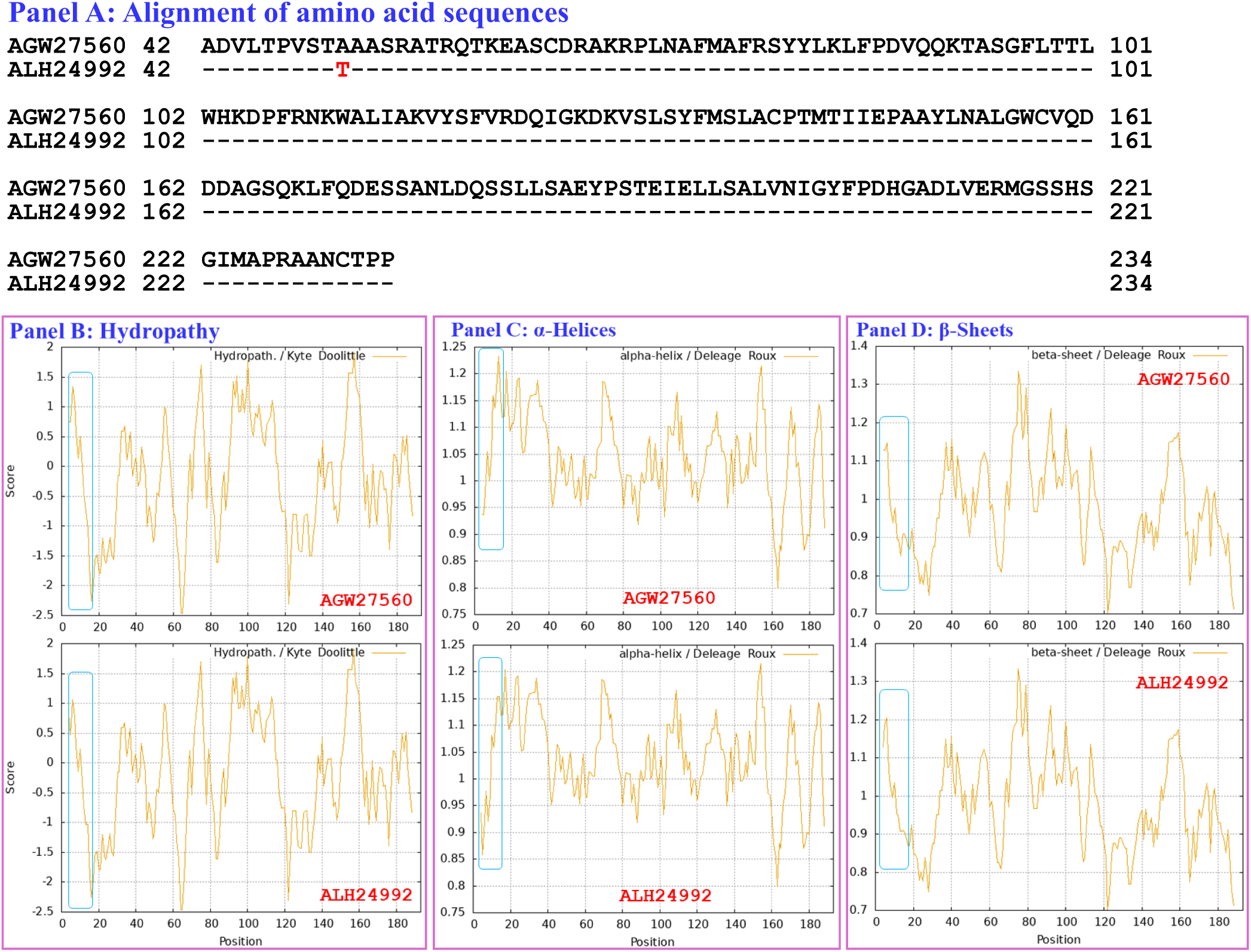
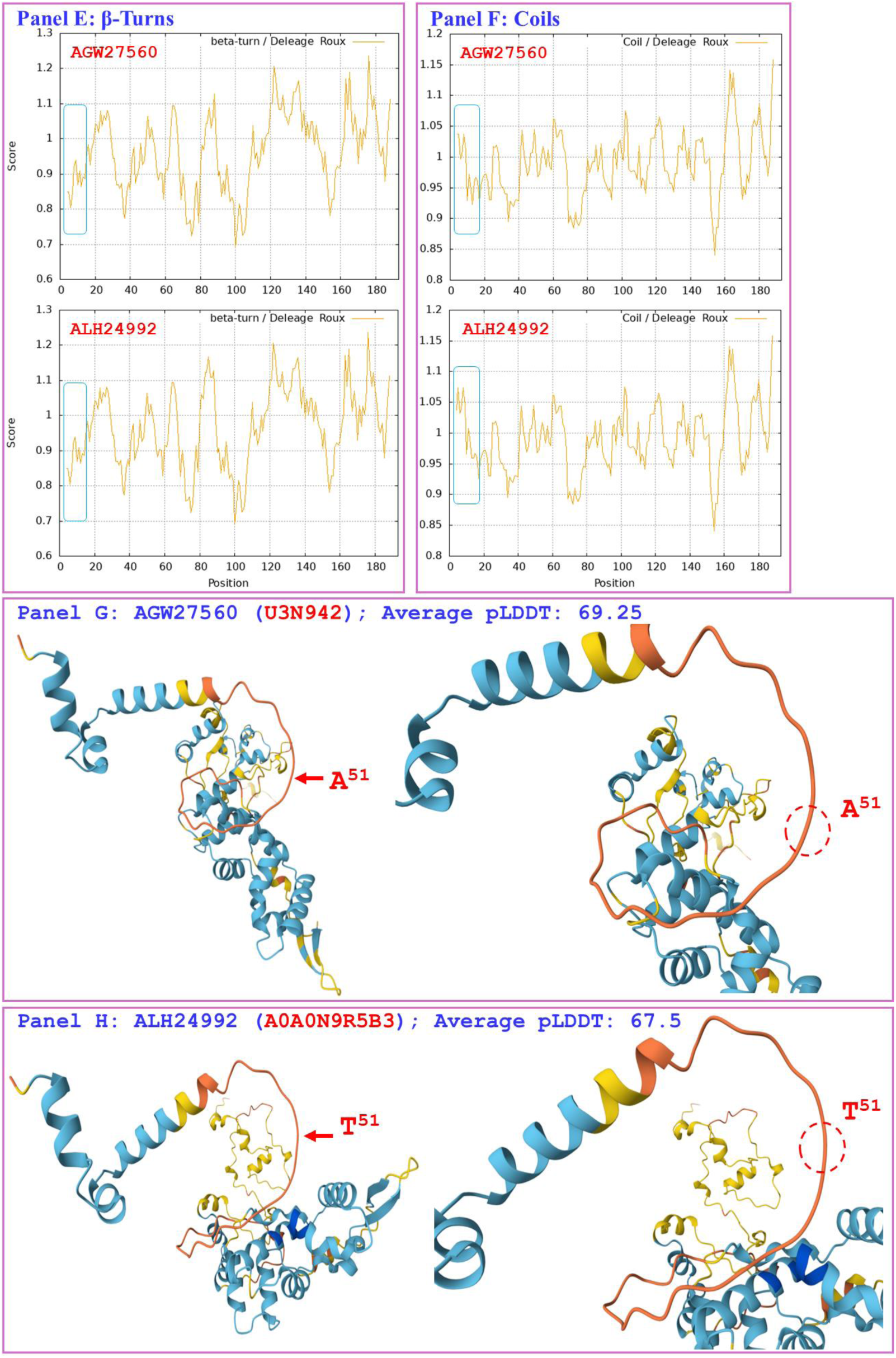
Correlation of the changes in the hydrophobicity and the primary, secondary, and tertiary structures of the MATα_HMGbox domains of MAT1-1-1 proteins: the reference protein AGW27560 (under the AlphaFold code U3N942) derived from the *H. sinensis* strain CS68-2-1229 and the mutant protein ALH24992 (under the AlphaFold code A0A0N9R5B3) derived from the wild-type *C. sinensis* isolate SC09_65. Panel A shows an alignment of the amino acid sequences of the MATα_HMGbox domains of the MAT1-1-1 proteins; the amino acid substitution is shown in red, whereas the hyphens indicate identical amino acid residues. The ExPASy ProtScale plots show the changes in hydrophobicity (Panel B) and in the 2D structure (Panels C−F for α-helices, β-sheets, β-turns, and coils, respectively) of the protein; the open rectangles in blue outline the changes in topology and waveform of the ExPASy plots. Panels G−H show the 3D structures of the full-length proteins on the left and the locally amplified structures surrounding the mutation site on the right. The model confidence for the AlphaFold-predicted 3D structures is indicated: 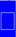 very high (pLDDT>90); 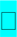 high (90>pLDDT>70); 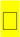 low (70>pLDDT>50); 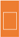 very low (pLDDT<50).

As shown in Panels G−H of Figure 3, which illustrate the 3D structures of the proteins, the substitution is located upstream of the core structure of 3 α-helices and may not directly participate in stabilizing the hydrophobic core structure of the DNA-binding domain [Baxevanis *et al*. 1995; Thapar 2015]. However, replacement of alanine with a threonine residue with reduced hydrophobicity caused altered stereostructure of the MATα_HMGbox domain of the MAT1-1-1 protein ALH24992 under the AlphaFold code A0A0N9R5B3, perhaps through long-range spatial interactions. The mutant MATα_HMGbox domain clustered into Branch a2 in the Bayesian clustering tree (*cf*. Figure 1, Table S5).

Figure 4 compares the hydrophobicities and the structure of the MATα_HMGbox domain of the MAT1-1-1 protein ALH24948 (under the AlphaFold code A0A0N9QUF3) derived from the wild-type *C. sinensis* isolate GS09_143 (*cf*. Table S1) [Zhang & Zhang 2015] with those of the MATα_HMGbox domain of the reference protein AGW27560 (under the AlphaFold code U3N942) derived from the *H. sinensis* strain CS68-2-1229 [Bushley *et al*. 2013]. Panel A shows substitutions of two adjacent residues: Q-to-S (glutamine to serine, both of which have polar side chains; hydropathy indices changed from −3.5 to −0.8 [Kyte & Doolittle 1982]) and I-to-F (aliphatic isoleucine to aromatic phenylalanine; hydropathy indices changed from 4.5 to 2.8), causing altered hydrophobicity as illustrated by the changes in topology and waveform in the hydropathy plot in Panel B. These changes alter the secondary structure surrounding the mutation sites in the MATα_HMGbox domain, as illustrated by the changes in topology and waveform in the ExPASy ProtScale plots for α-helices, β-sheets, β-turns, and coils shown in Panels C−F. As shown in Panels G−H for the diagrams of 3D structure, the adjacent amino acid substitutions are located at the end of one of the 3 α-helices that form the hydrophobic core stereostructure [Baxevanis *et al*. 1995; Thapar 2015], and they cause the altered tertiary structure of the MATα_HMGbox domain of the mutant protein ALH24948 under the AlphaFold code A0A0N9QUF3. The mutant MATα_HMGbox domain clustered into Branch e2 in the Bayesian clustering tree (*cf*. Figure 1, Table S5).

**Figure 4.**
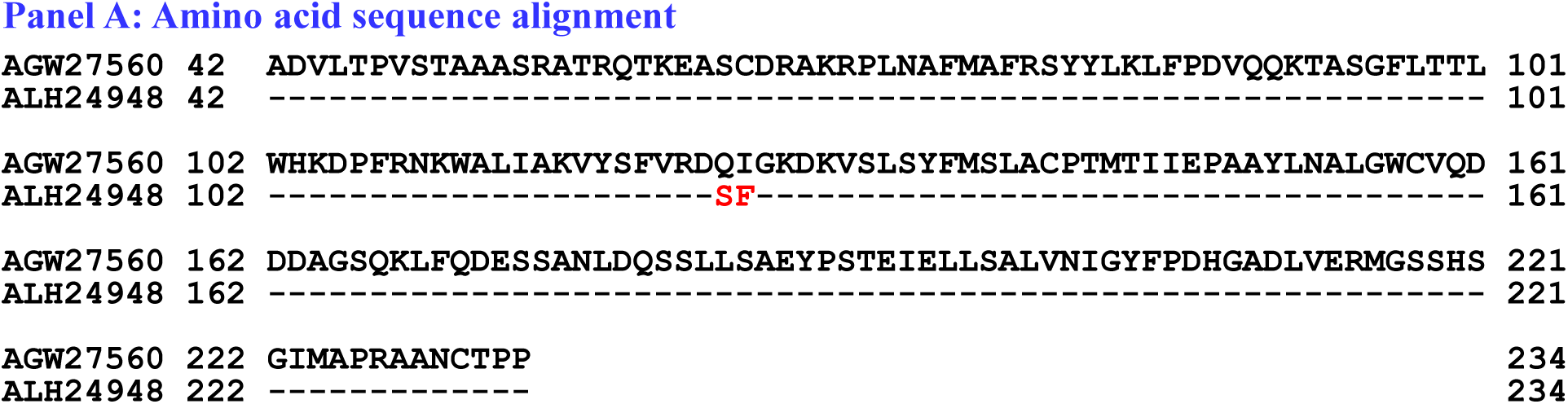
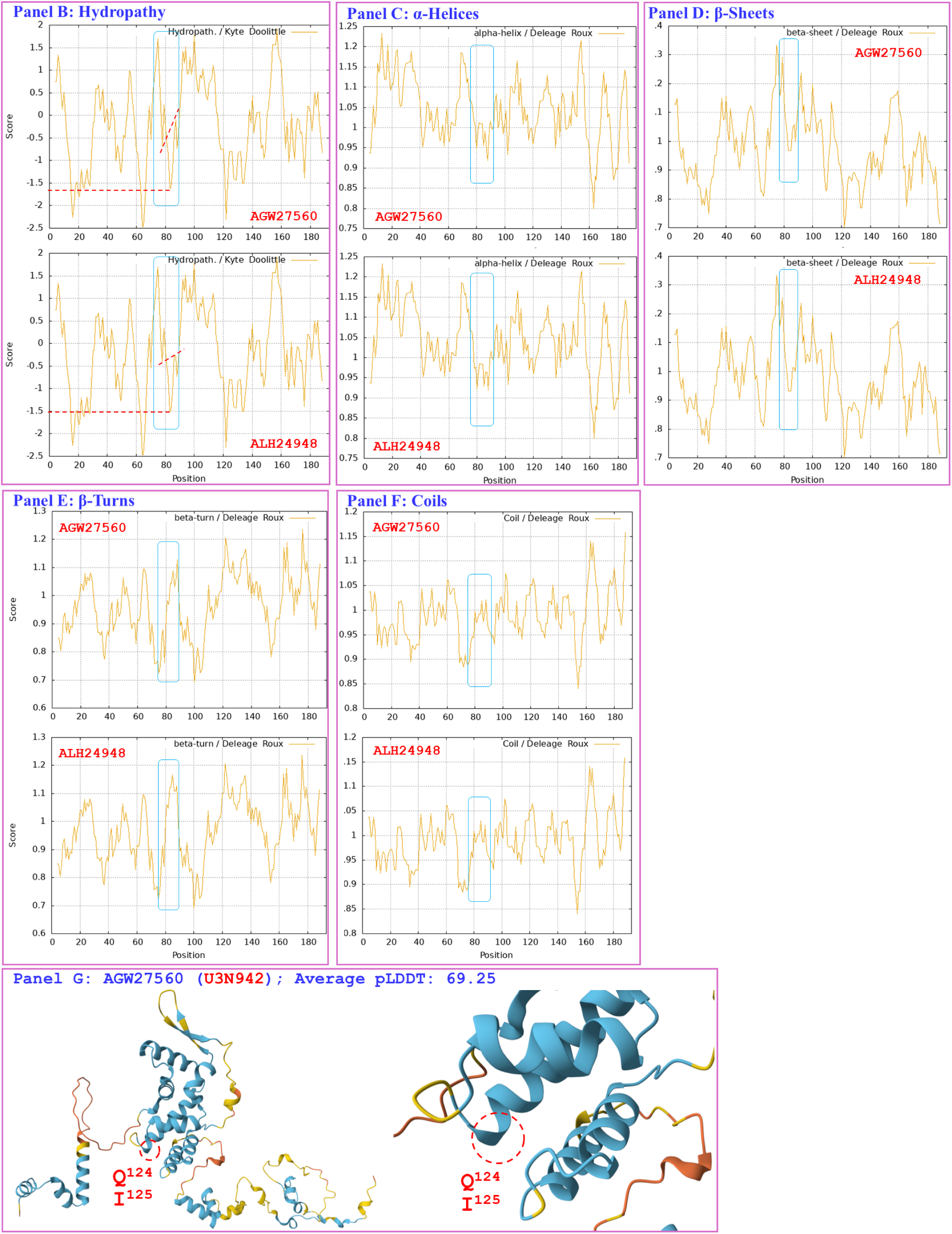
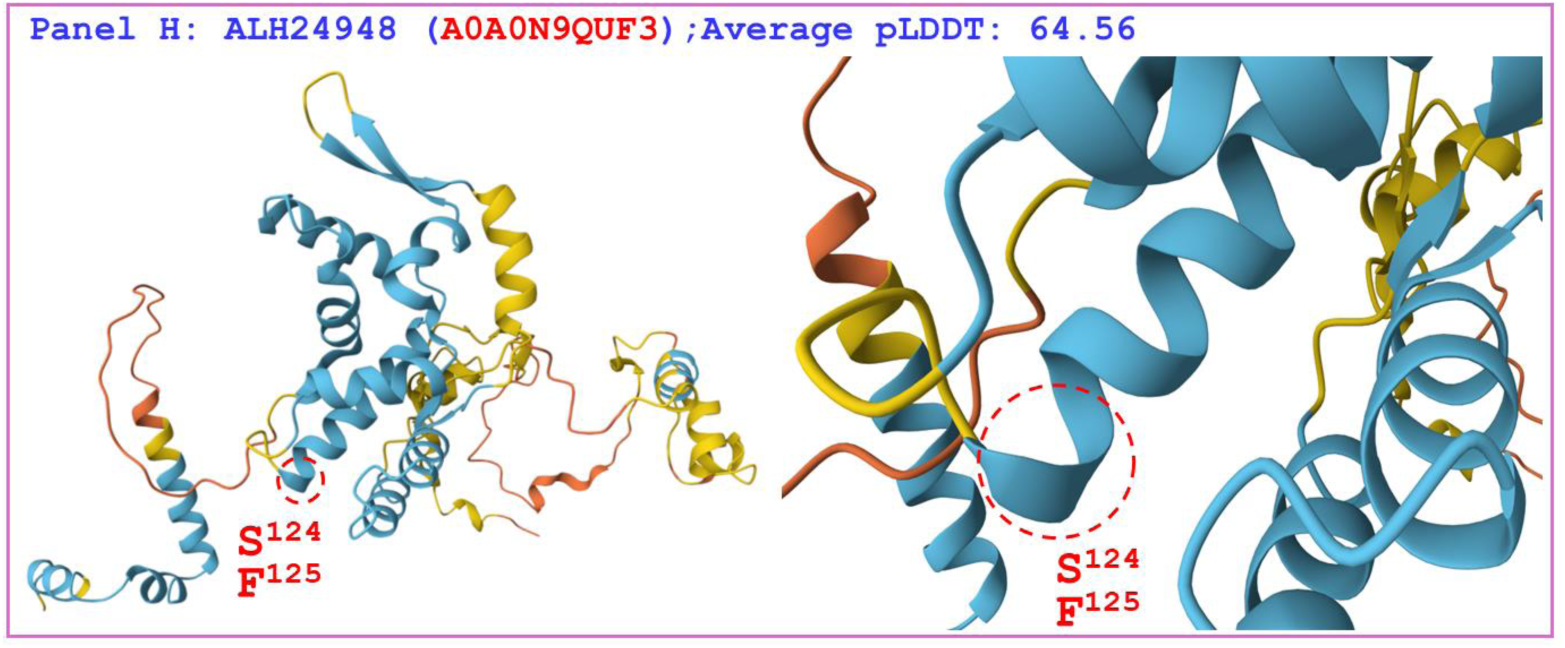
Correlations among the changes in the primary, secondary, and tertiary structures of the MATα_HMGbox domains of MAT1-1-1 proteins: the reference protein AGW27560 (under the AlphaFold code U3N942) derived from the *H. sinensis* strain CS68-2-1229 and the mutant MAT1-1-1 protein ALH24948 (under the AlphaFold code A0A0N9QUF3) derived from the wild-type *C. sinensis* isolate GS09_143. Panel A shows an alignment of the MATα_HMGbox domain sequences of the MAT1-1-1 proteins; amino acid substitutions are shown in red, whereas the hyphens indicate identical amino acid residues. The ExPASy ProtScale plots show the changes in hydrophobicity (Panel B) and in the 2D structure (Panels C−F for α-helices, β-sheets, β-turns, and coils, respectively) of the protein; the open rectangles in blue outline the changes in topology and waveform shown in the ExPASy plots. Panels G−H show the 3D structures of the full-length proteins on the left and the locally amplified structures at the sites of substitutions on the right. The model confidence for the AlphaFold-predicted 3D structures is indicated as follows: 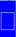 very high (pLDDT>90); 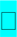 high (90>pLDDT>70); 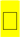 low (70>pLDDT>50); 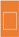 very low (pLDDT<50).

Figure 5 compares the hydrophobicities and the structures of the MATα_HMGbox domains of the MAT1-1-1 proteins ALH25043, ALH25045, ALH25046, and ALH25048 (under the AlphaFold code A0A0N9QMS9) derived from the wild-type *C. sinensis* isolates YN09_22, YN09_51, YN09_6, and YN09_64, respectively, (*cf*. Table S1) [Zhang & Zhang 2015] with those of the MATα_HMGbox domain of the reference protein AGW27560 (under the AlphaFold code U3N942) derived from the *H. sinensis* strain CS68-2-1229 [Bushley *et al*. 2013]. Panel A shows substitutions of R-to-I (basic arginine to aliphatic isoleucine; hydropathy indices changed from −4.5 to 4.5 [Kyte & Doolittle 1982]) and P-to-T (proline to threonine, both of which possess polar neutral side chains; hydropathy indices changed from −1.6 to −0.7). These substitutions result in increased hydrophobicity, as illustrated by the changes in topology and waveform that are apparent in the hydropathy plot in Panel B. They also result in altered secondary structure surrounding the mutation sites in the MATα_HMGbox domains of the MAT1-1-1 proteins ALH25043, ALH25045, ALH25046, and ALH25048, as illustrated by the changes in topology and waveform shown in the ExPASy ProtScale plots for α-helices, β-sheets, β-turns, and coils in Panels C−F. As shown in the diagrams of 3D structure in Panels G−H, the sites of the R-to-I and P-to-T substitutions are inside and outside, respectively, the hydrophobic core formed by the 3 α-helices, and the substantial increases in hydrophobicity alter the tertiary structures of the MATα_HMGbox domains of MAT1-1-1 proteins under the AlphaFold code A0A0N9QMS9. The mutant MATα_HMGbox domain clustered into Cluster 2 in the Bayesian clustering tree (*cf*. Figure 1, Table S5).

**Figure 5.**
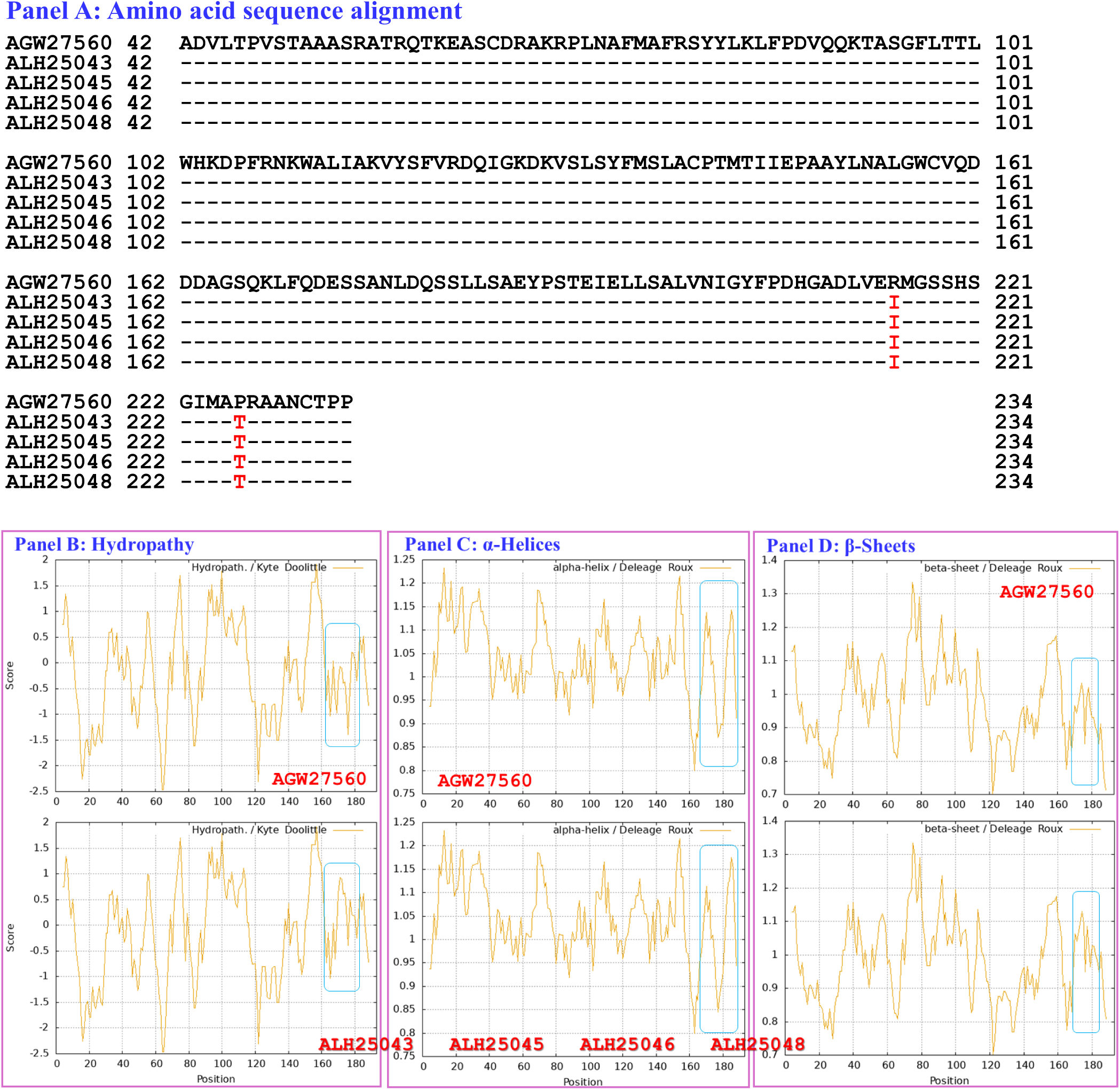
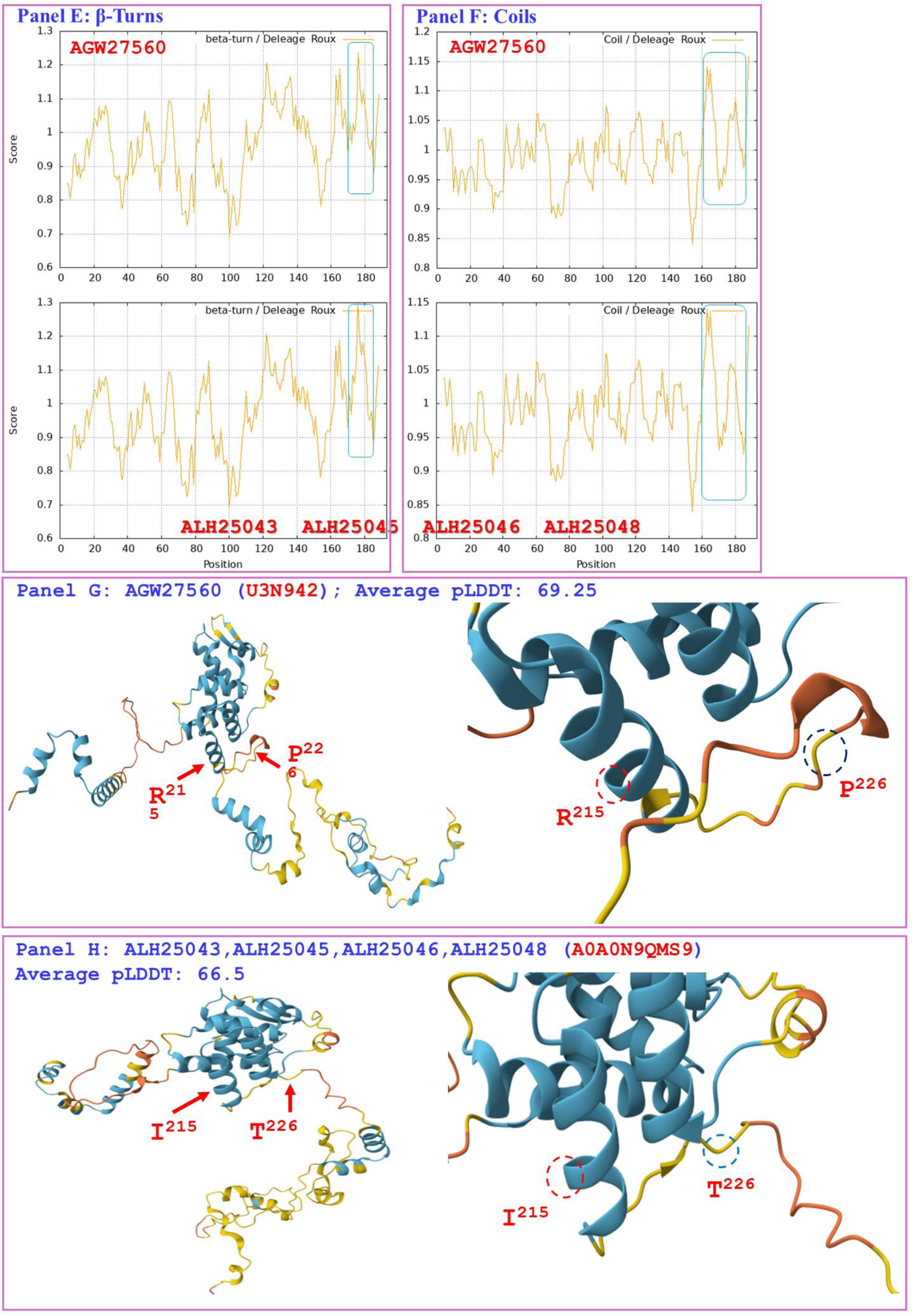
Correlation of changes in the primary, secondary, and tertiary structures of the MATα_HMGbox domains of the MAT1-1-1 proteins: the reference protein AGW27560 (under the AlphaFold code U3N942) is derived from the *H. sinensis* strain CS68-2-1229 and the mutant MAT1-1-1 proteins ALH25043, ALH25045, ALH25046, and ALH25048 (under the AlphaFold code A0A0N9QMS9) are derived from the wild-type *C. sinensis* isolates YN09_22, YN09_51, YN09_6, and YN09_64, respectively. Panel A shows an alignment of the amino acid sequences of the MATα_HMGbox domains of the MAT1-1-1 proteins; amino acid substitutions are shown in red, whereas the hyphens indicate identical amino acid residues. The ExPASy ProtScale plots show the changes in hydrophobicity (Panel B) and in the 2D structure (Panels C−F for α-helices, β-sheets, β-turns, and coils, respectively) of the proteins; the open rectangles in blue outline the changes in topology and waveform of the plots. Panels G−H show the 3D structures of the full-length proteins on the left and the locally amplified structures at the sites of substitutions on the right. The model confidence for the AlphaFold-predicted 3D structures is indicated as follows: 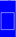 very high (pLDDT>90); 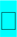 high (90>pLDDT>70); 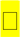 low (70>pLDDT>50); 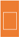 very low (pLDDT<50).

Figure 6 compares the hydrophobicities and structures of the MATα_HMGbox domains of the MAT1-1-1 proteins ALH25054, ALH24951, ALH24952, ALH24953, ALH24962, ALH24963, ALH24964, ALH24966, and ALH24994 (under the AlphaFold code A0A0N7G845) derived from the wild-type *C. sinensis* isolates GS09_311, GS09_229, GS09_281, GS10_1, QH09_164, QH09_173, QH09_201, QH09_210, and SC09_87, respectively (*cf*. Table S1) [Zhang & Zhang 2015] with those of the reference protein AGW27560 (under the AlphaFold code U3N942) derived from the *H. sinensis* strain CS68-2-1229 [Bushley *et al*. 2013]. Panel A shows I-to-L substitutions (aliphatic isoleucine to aliphatic leucine, hydropathy indices changed from 4.5 to 3.8 [Kyte & Doolittle 1982]); these substitutions resulted in reduced hydrophobicity, as illustrated by the changes in topology and waveform shown in Panel B for the hydropathy plot.

**Figure 6.**
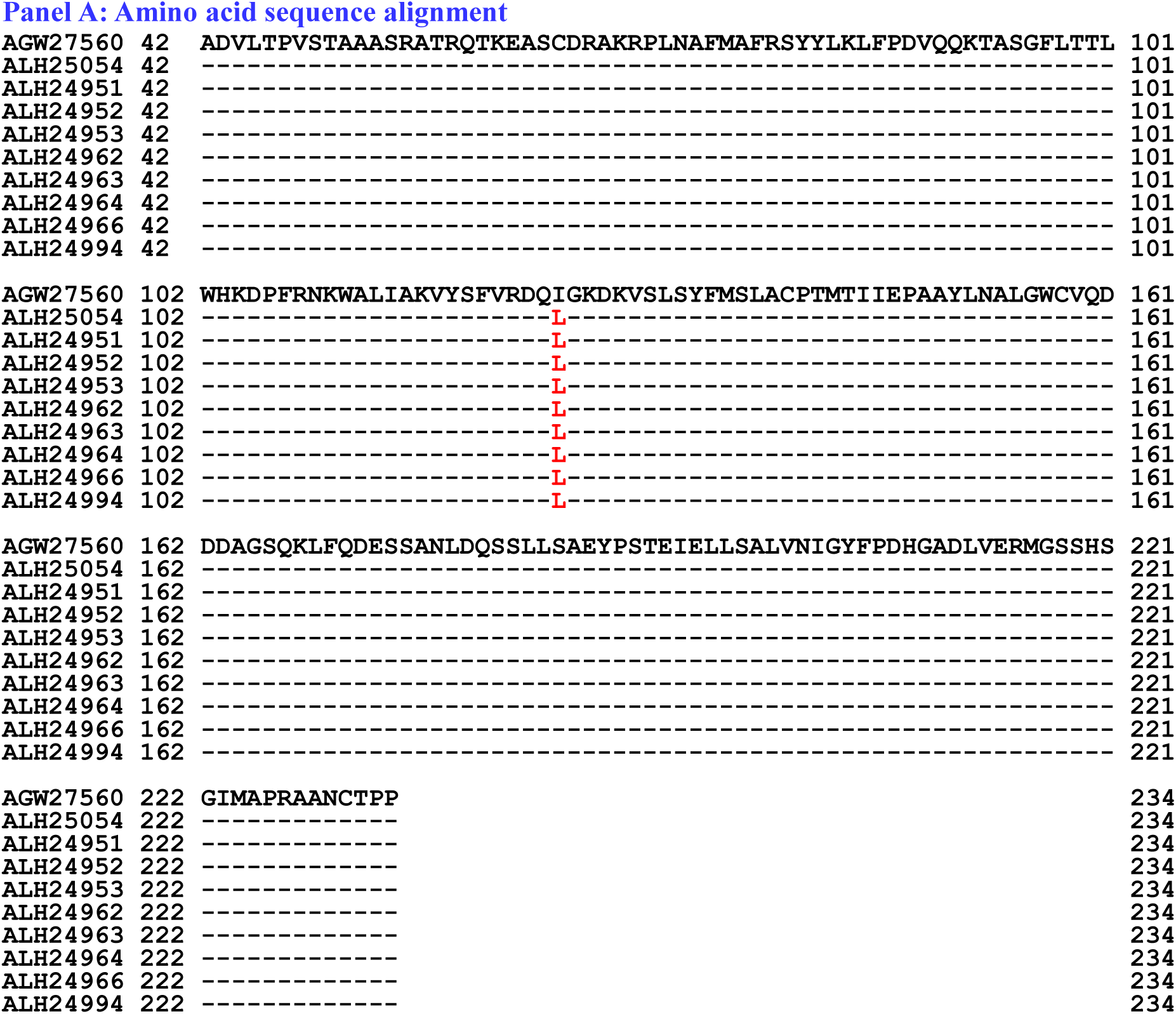
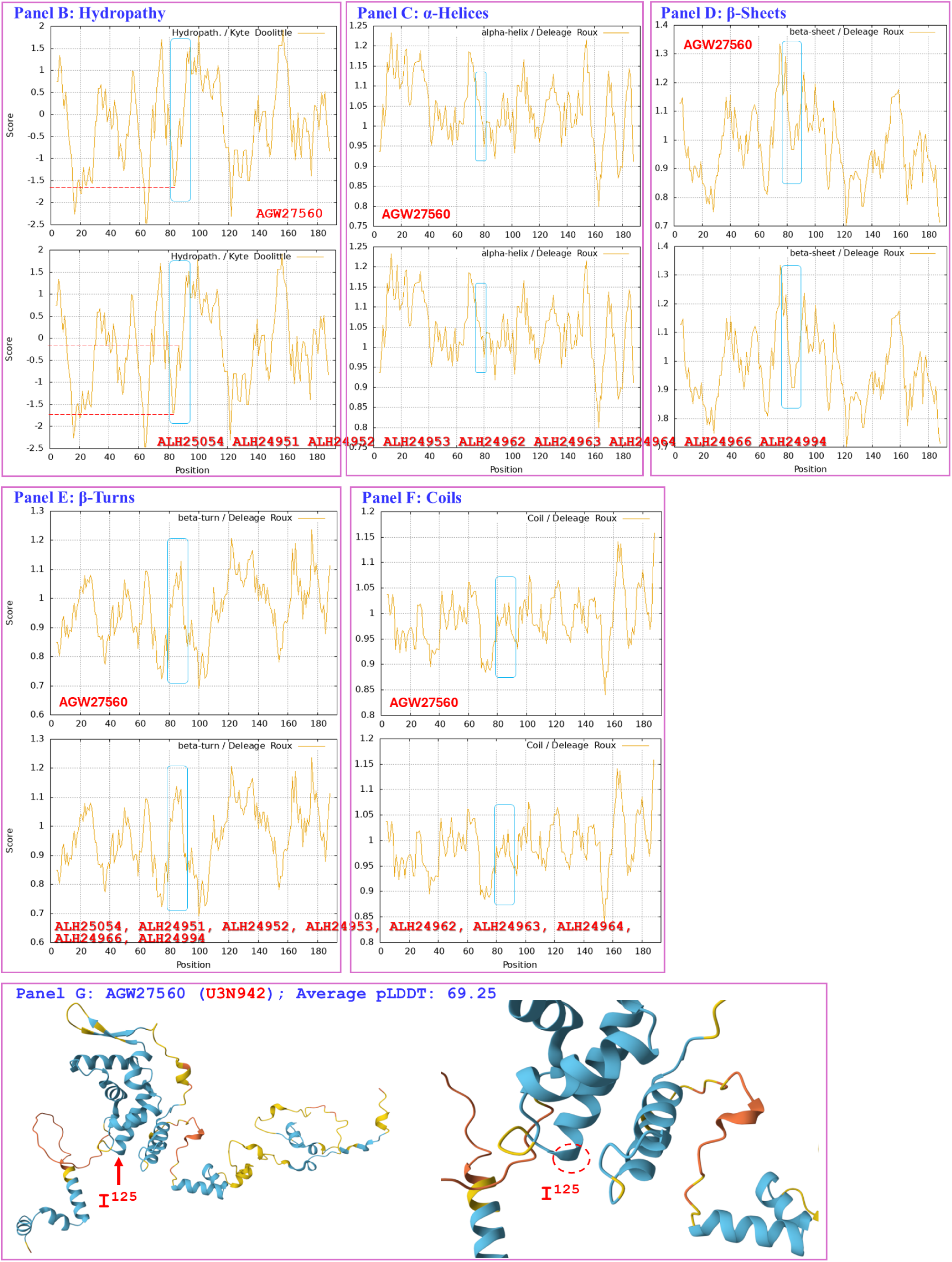
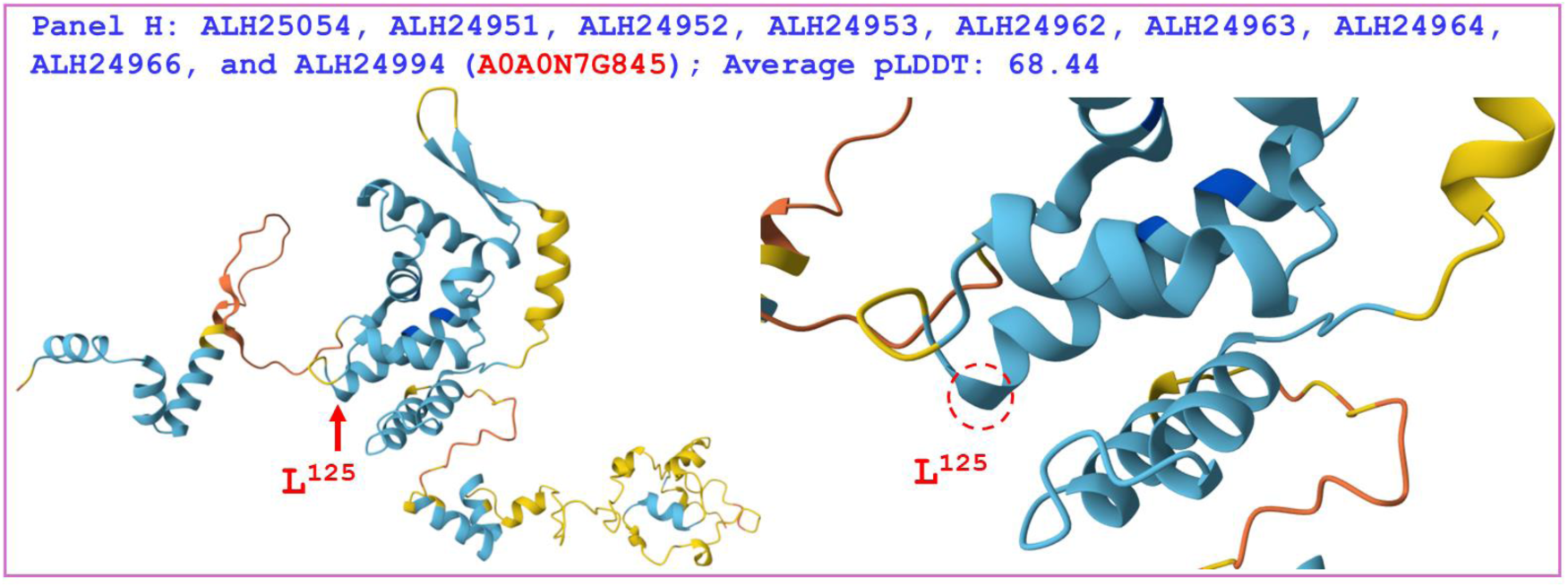
Correlation of changes in the primary, secondary, and tertiary structures of the MATα_HMGbox domains in the MAT1-1-1 proteins. The reference protein AGW27560 (under the AlphaFold code U3N942) is derived from the *H. sinensis* strain CS68-2-1229, and the mutant MAT1-1-1 proteins ALH25054, ALH24951, ALH24952, ALH24953, ALH24962, ALH24963, ALH24964, ALH24966, and ALH24994 (under the AlphaFold code A0A0N7G845) are derived from the wild-type *C. sinensis* isolates GS09_311, GS09_229, GS09_281, GS10_1, QH09_164, QH09_173, QH09_201, QH09_210, and SC09_87, respectively. Panel A shows an alignment of the amino acid sequences of the MATα_HMGbox domains of the MAT1-1-1 proteins; amino acid substitutions are shown in red, whereas the hyphens indicate identical amino acid residues. The ExPASy ProtScale plots show the changes in hydrophobicity (Panel B) and in the 2D structure (Panels C−F for α-helices, β-sheets, β-turns, and coils, respectively) of the proteins; the open rectangles in blue outline the changes in topology and waveform shown in the plots. Panels G−H show the 3D structures of the full-length proteins on the left and the locally amplified structures at the site of substitution on the right. The model confidence for the AlphaFold-predicted 3D structures is indicated as follows: 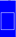 very high (pLDDT>90); 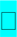 high (90>pLDDT>70); 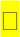 low (70>pLDDT>50); 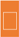 very low (pLDDT<50).

As shown in Panels C−F of Figure 6, the secondary structure of the proteins near the mutation sites in the MATα_HMGbox domains of the MAT1-1-1 proteins ALH25054, ALH24951, ALH24952, ALH24953, ALH24962, ALH24963, ALH24964, ALH24966, and ALH24994 (*cf*. Table S1) was altered, as illustrated by the changes in topology and waveform in the ExPASy ProtScale plots for α-helices, β-sheets, β-turns, and coils, respectively. As shown in the representations of the 3D structures presented in Panels G−H, the substituted amino acid residues are located within the core structures of the α-helices, and they destabilize the hydrophobic core of the protein [Baxevanis *et al*. 1995; Thapar 2015]; this alters the tertiary structures of the MATα_HMGbox domains of the MAT1-1-1 proteins under the AlphaFold code A0A0N7G845, which clustered into Branch e1 in the Bayesian clustering tree (*cf*. Figure 1, Table S5).

Figure 7 shows the hydrophobicities and structures of the MATα_HMGbox domains of the MAT1-1-1 proteins ALH24999 and ALH25057 (under the AlphaFold code A0A0N9QMT4) and the MAT1-1-1 protein ALH25001 (under the AlphaFold code A0A0N9R4Q4) derived from the wild-type *C. sinensis* isolates XZ07_H2, XZ12_16, and XZ05_2, respectively, (*cf*. Table S1) [Zhang & Zhang 2015], compared with the reference protein AGW27560 (under the AlphaFold code U3N942) derived from the *H. sinensis* strain CS68-2-1229 [Bushley *et al*. 2013]. Panel A shows substitutions of A-to-V (aliphatic alanine to aliphatic valine; hydropathy indices changed from 1.8 to 4.2 [Kyte & Doolittle 1982]) and A-to-T (aliphatic isoleucine to threonine, an amino acid that contains polar neutral side chains; hydropathy indices changed from 1.8 to −0.7). These substitutions altered the hydrophobicity of the protein, as illustrated by the changes in topology and waveform shown in the hydropathy plot in Panel B. These changes altered the secondary structure surrounding the mutation sites in the MATα_HMGbox domain of the MAT1-1-1 proteins ALH24999, ALH25057, and ALH25001, as illustrated by the changes in topology and waveform in the ExPASy ProtScale plots for α-helices, β-sheets, β-turns, and coils shown in Panels C−F, respectively. As shown in the 3D structures represented in Panels G−H, the mutation sites are located upstream of the hydrophobic core structure of 3 α-helices in the MATα_HMGbox domains, and mutations at these sites alter the tertiary structures of the MATα_HMGbox domains of MAT1-1-1 proteins under the AlphaFold codes A0A0N9QMT4 and A0A0N9R4Q4, which clustered into Cluster d in the Bayesian clustering tree (*cf*. Figure 1, Table S5).

**Figure 7.**
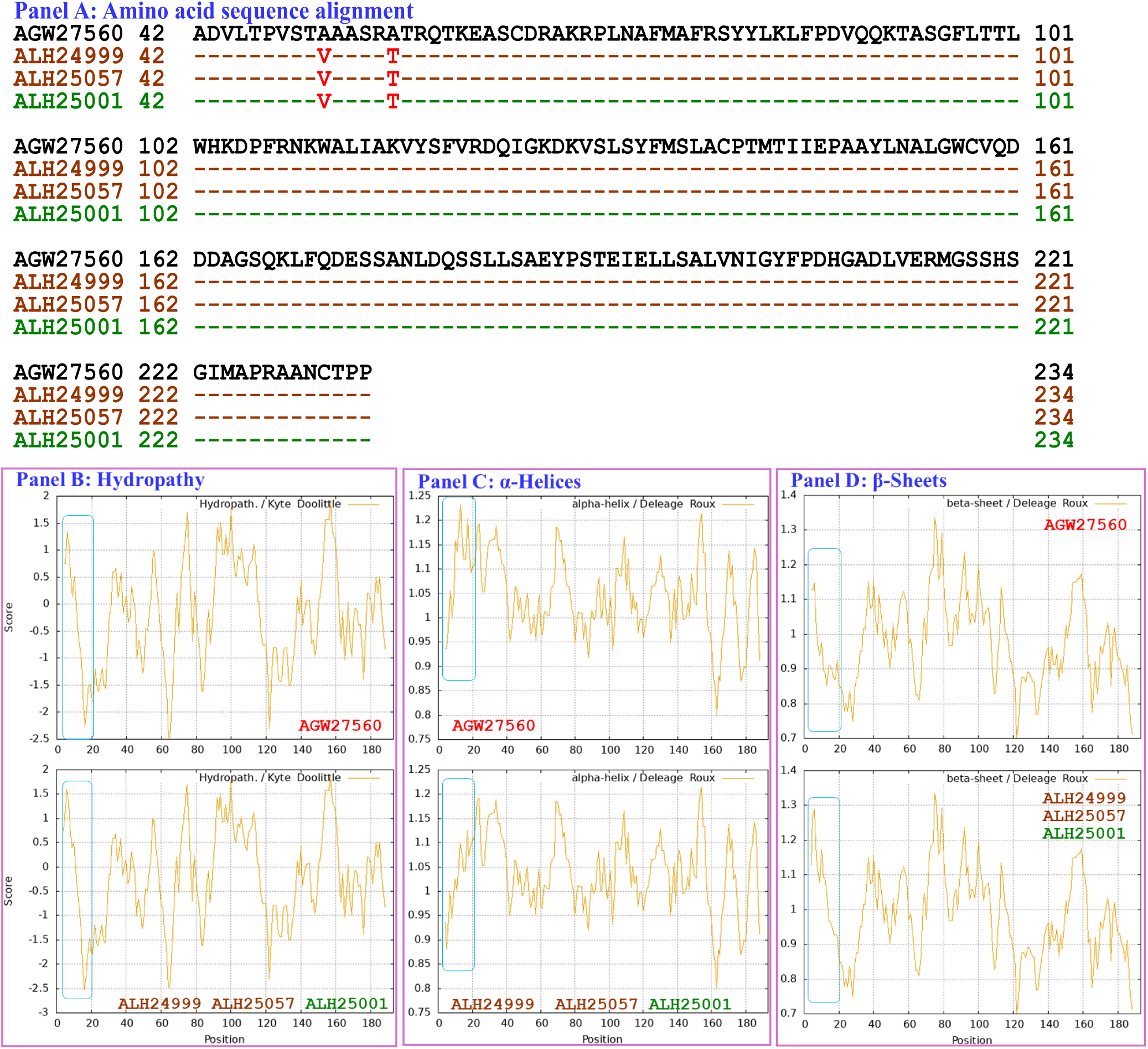
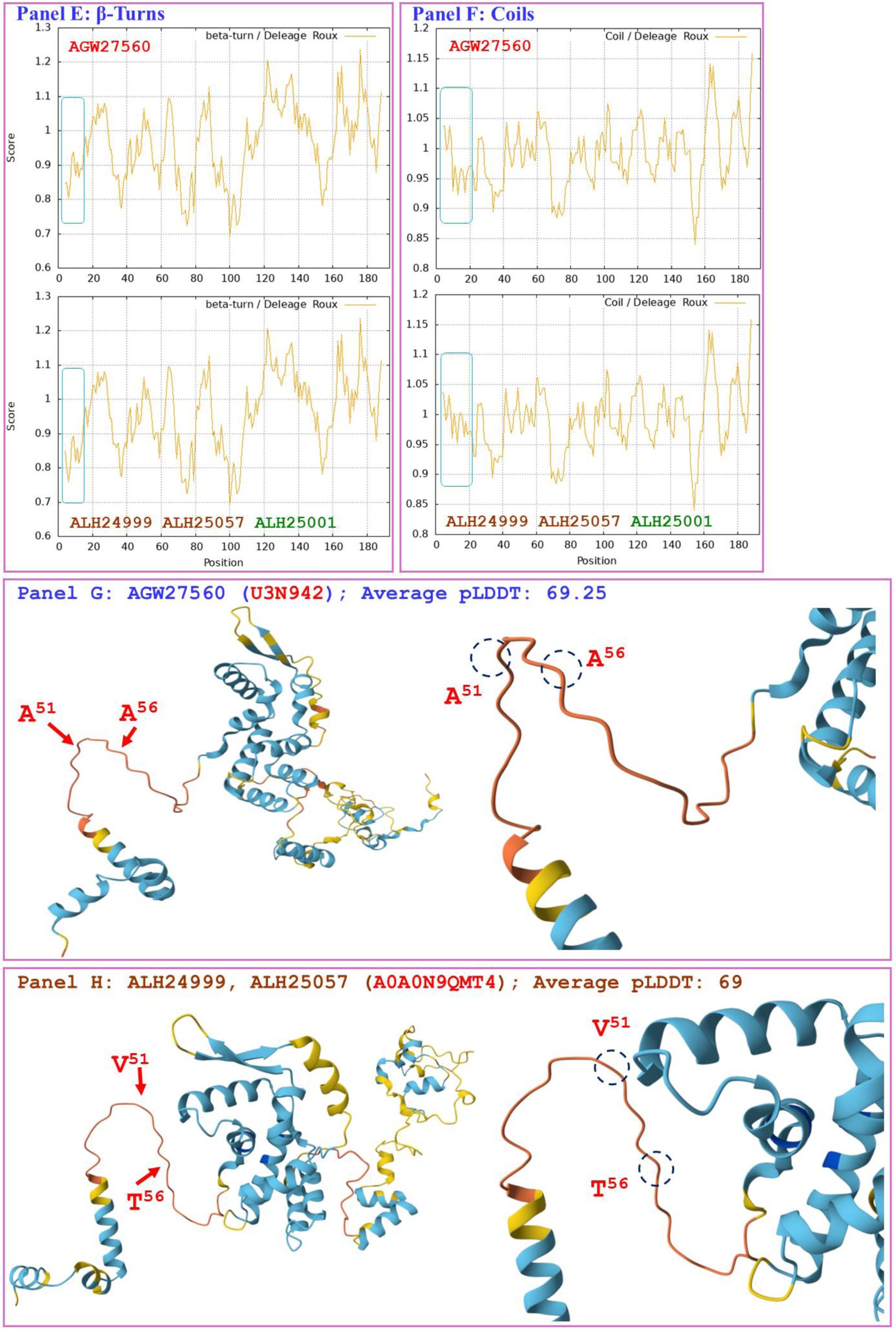
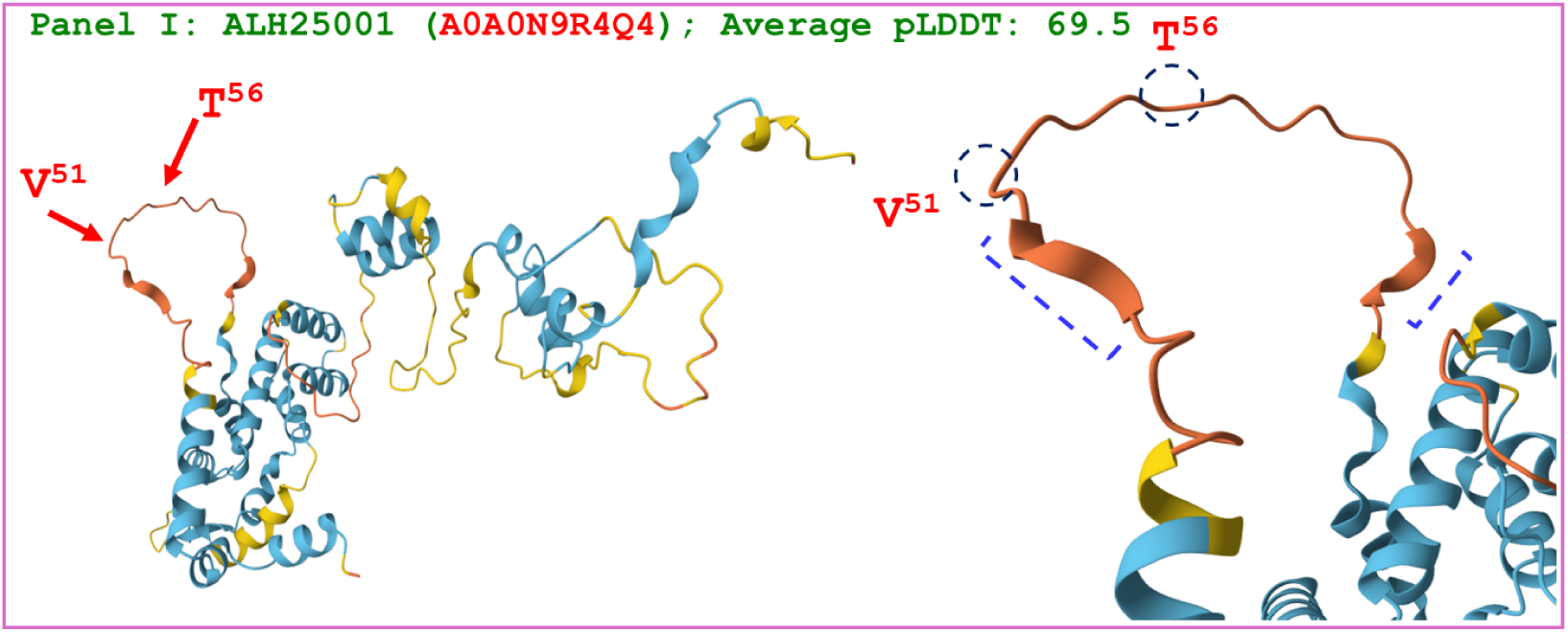
Correlation of the changes in the primary, secondary, and tertiary structure of the MATα_HMGbox domains in the MAT1-1-1 proteins: the reference protein AGW27560 (under the AlphaFold code U3N942) is derived from the *H. sinensis* strain CS68-2-1229, and the mutant MAT1-1-1 proteins ALH24999 and ALH25057 (under the AlphaFold code A0A0N9QMT4) and the protein ALH25001 (under the AlphaFold code A0A0N9R4Q4) are derived from the wild-type *C. sinensis* isolates XZ07_H2, XZ12_16, and XZ05_2, respectively. Panel A shows an alignment of the amino acid sequences of the MATα_HMGbox domains of the MAT1-1-1 proteins; amino acid substitutions are shown in red; the sequences displayed in brown and green represent the MAT1-1-1 proteins under the AlphaFold codes A0A0N9QMT4 and A0A0N9R4Q4, respectively, and the hyphens indicate identical amino acid residues. The ExPASy ProtScale plots show the changes in hydrophobicity (Panel B) and in the 2D structure (Panels C−F, which display the α-helices, β-sheets, β-turns, and coils, respectively) of the proteins. The open rectangles in blue outline the changes in topology and waveform shown in the plots. Panels G−H show the 3D structures of the full-length proteins on the left and the locally amplified structures at the substitution sites on the right. The model confidence for the AlphaFold-predicted 3D structures is indicated as follows: 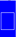 very high (pLDDT>90); 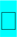 high (90>pLDDT>70); 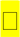 low (70>pLDDT>50); 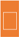 very low (pLDDT<50).

Although Panels A of Figure 7 shows the same substitutions of A-to-V and A-to-T in the MATα_HMGbox domains of the mutant proteins ALH24999, ALH25057, and ALH25001 (under the AlphaFold codes A0A0N9QMT4 and A0A0N9R4Q4) [Zhang & Zhang 2015], Panels H and I show different tertiary structures for the MATα_HMGbox domains of the mutant MAT1-1-1 proteins under the 2 AlphaFold codes. Specifically, the protein ALH25001 contains 2 extra short β-sheet structures upstream and downstream of the mutation sites in the MATα_HMGbox domain shown in Panel I; these are marked with dashed half brackets in blue. In contrast, the distinct tertiary structures for the mutant proteins ALH24999 and ALH25057 without the β-sheet structures are shown in Panel H.

Figure 8 compares the hydrophobicities and structures of the MATα_HMGbox domain of the MAT1-1-1 protein ALH25003 (under the AlphaFold code A0A0N7G850) derived from the wild-type *C. sinensis* isolate XZ05_6 (*cf*. Table S1) [Zhang & Zhang 2015] with those of the reference protein AGW27560 (under the AlphaFold code U3N942) derived from the *H. sinensis* strain CS68-2-1229 [Bushley *et al*. 2013]. Panel A shows an S-to-G substitution (serine, which has polar neutral side chains, is replaced by the unique amino acid glycine; hydropathy indices changed from −0.8 to −0.4 [Kyte & Doolittle 1982]). This substitution resulted in a slight increase in hydrophobicity, as illustrated by the changes in topology and waveform shown in the hydropathy plot in Panel B. These changes altered the secondary structure of the mutant MAT1-1-1 protein ALH25003 in the region surrounding the mutation site in the MATα_HMGbox domain, as illustrated by the changes in topology and waveform in the ExPASy ProtScale plots for α-helices, β-sheets, β-turns, and coils shown in Panels C−F. As shown in the representations of the protein’s 3D structure in Panels G−H, the amino acid residue (serine), which is located at the end of one of the 3 α-helices that form the hydrophobic core structure of the MATα_HMGbox domain [Baxevanis *et al*. 1995; Thapar 2015], was substituted by glycine, which became the first residue after the α-helix in the domain. This single-residue substitution plus additional E-to-K and Y-to-H substitutions downstream of the MATα_HMGbox domain (*cf*. Figure S1) significantly altered the tertiary structure of the protein ALH25003 under the unique AlphaFold code A0A0N7G850. The mutant MATα_HMGbox domain clustered into Branch a2 in the Bayesian clustering tree (*cf*. Figure 1, Table S5).

**Figure 8.**
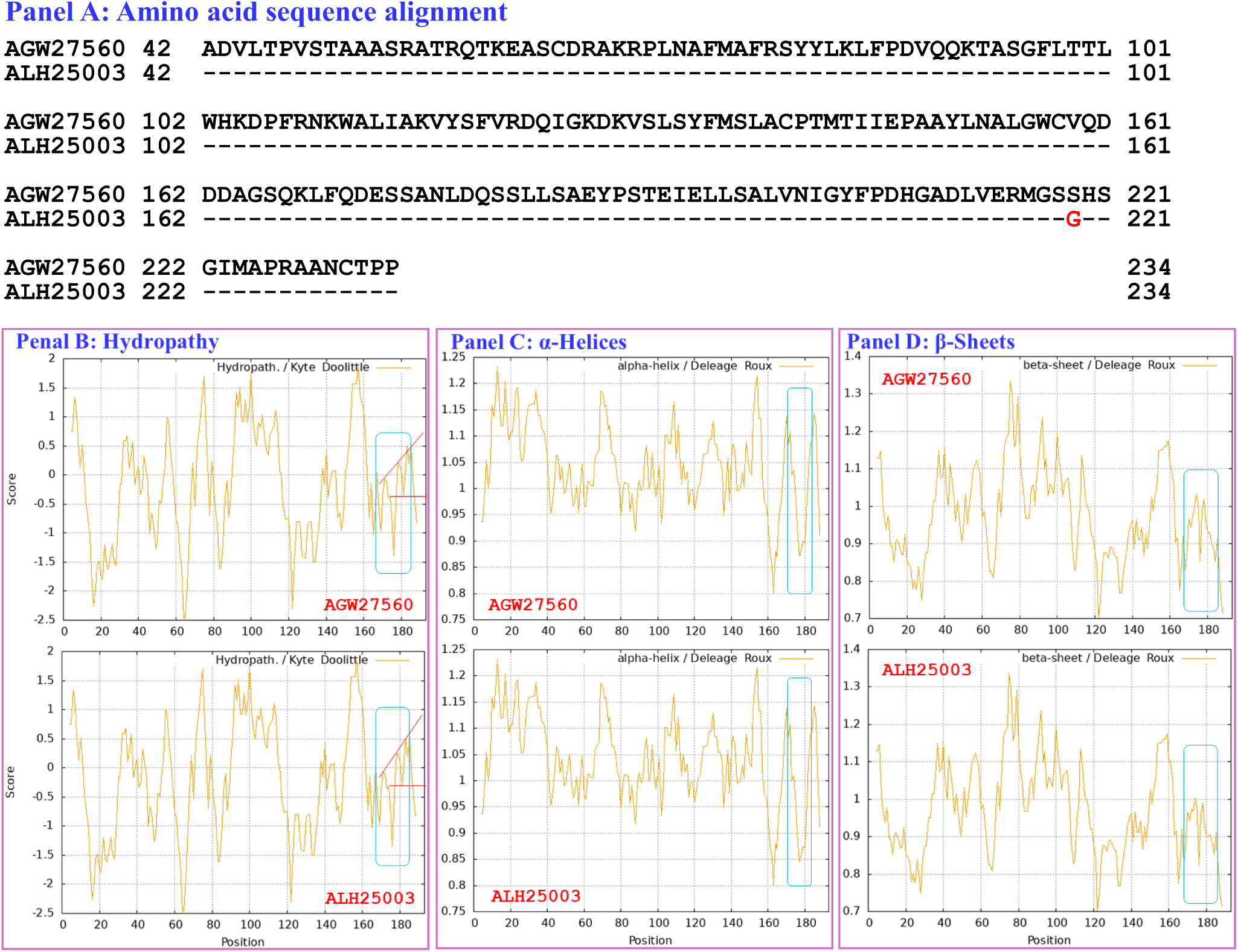
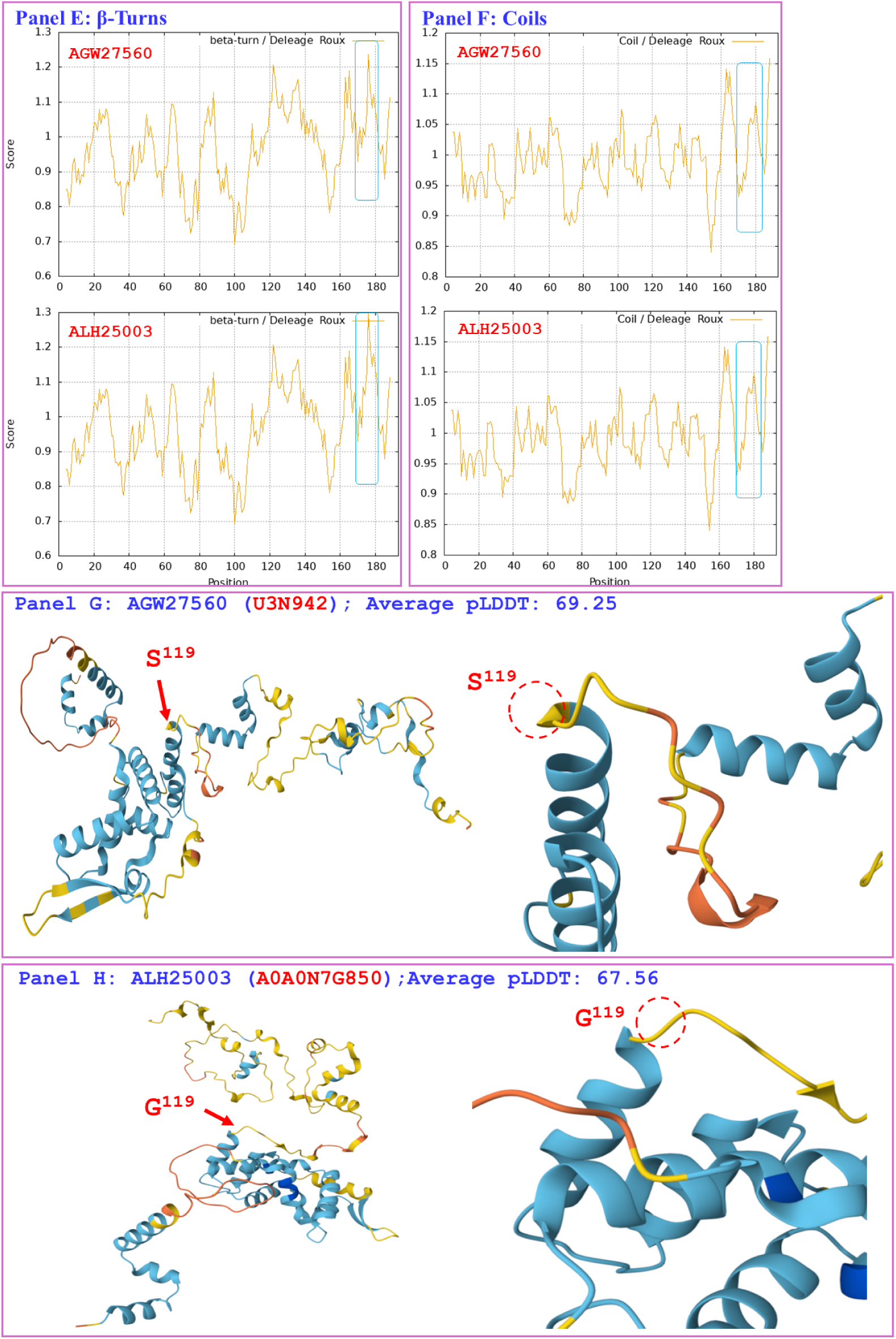
Correlation of the changes in the primary, secondary, and tertiary structures of the MATα_HMGbox domains of the MAT1-1-1 proteins: the reference protein AGW27560 (under the AlphaFold code U3N942) is derived from the *H. sinensis* strain CS68-2-1229, and the mutant MAT1-1-1 protein ALH25003 (under the AlphaFold code A0A0N7G850) is derived from the wild-type *C. sinensis* isolate XZ05_6. Panel A shows an alignment of the amino acid sequences of the MATα_HMGbox domains of the MAT1-1-1 proteins; the amino acid substitution is shown in red, whereas the hyphens indicate identical amino acid residues. The ExPASy ProtScale plots show the changes in the hydrophobicity (Panel B) and in the 2D structure of the protein (Panels C−F for the α-helices, β-sheets, β-turns, and coils, respectively) of the protein, and the open rectangles in blue outline the changes in topology and waveform shown in the plots. Panels G−H show the 3D structures of the full-length proteins on the left and the locally amplified structures at the sites of substitutions on the right. The model confidence for the AlphaFold-predicted 3D structures is indicated as follows: 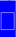 very high (pLDDT>90); 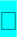 high (90>pLDDT>70); 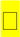 low (70>pLDDT>50); 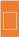 very low (pLDDT<50).

Aggregating the data presented in Section III-5, Panels A of Figures 3−8 demonstrate the 1−2 amino acid substitutions in the MATα_HMGbox domains of MAT1-1-1 proteins under various AlphaFold 3D structure codes derived from wild-type *C. sinensis* isolates compared with the reference MAT1-1-1 protein AGW27560 under the AlphaFold code U3N942 [Bushley *et al*. 2013]. The substitutions within the MATα_HMGbox domains caused changes in the topologies and waveforms shown in the ExPASy ProtScale plots in Panels B−F of Figures 3−8. These changes indicate the changes in the hydrophobicity and secondary structure (α-helices, β-sheets, β-turns, and coils) of the mutant full-length proteins. The changes in hydrophobicity and in the primary and secondary structures of the MATα_HMGbox domains altered the AlphaFold 3D structures of the domains of the proteins under various AlphaFold codes (*cf*. Panels H of Figures 3−8 and Panel I of Figure 7). The mutant domain sequences containing 1 or 2 amino acid substitutions were clustered into different clades in the Bayesian clustering tree (*cf*. Figure 1, Table S5), and the altered 3D structures of the proteins ultimately changed the DNA binding affinities, and the functionalities of gene transcriptional regulation related to the sexual reproduction of *O. sinensis*.

### III-6 Diverse primary and secondary structures of the MATα_HMGbox domains of MAT1-1-1 proteins encoded by the genome and metatranscriptome assemblies of *H. sinensis* and *C. sinensis* insect‒fungal complexes

Figure 9 compares the hydrophobicities and secondary structures (α-helices, β-sheets, β-turns, and coils) of the MATα_HMGbox domains of the MAT1-1-1 proteins encoded by the genome assemblies JAAVMX010000001 (6,699,061→6,699,153 & 6,699,206→6,699,637) and LKHE01001116 (4183←4620 & 4667←4759) of the *H. sinensis* strains IOZ07 and 1229 [Li *et al*. 2016a; Shu *et al*. 2020] with those of the reference MAT1-1-1 protein AGW27560 derived from the *H. sinensis* strain CS68-2-1229 [Bushley *et al*. 2013]. Panel A shows a Y-to-M substitution (aromatic tyrosine to methionine, an amino acid that has polar neutral side chains; hydropathy indices changed from −1.3 to 1.9 [Kyte & Doolittle 1982]). This substitution caused increased hydrophobicity in the MATα_HMGbox domains of the MAT1-1-1 proteins encoded by the genome assemblies JAAVMX010000001 and LKHE01001116 [Li *et al*. 2016a; Shu *et al*. 2020], as illustrated by the changes in topology and waveform in the hydropathy plot shown in Panel B. These changes resulted in altered secondary structures surrounding the mutation sites in the MATα_HMGbox domains of the MAT1-1-1 proteins, as illustrated in the ExPASy ProtScale plots for α-helices, β-sheets, β-turns, and coils in Panels C−F, respectively. The MATα_HMGbox domains of the MAT1-1-1 proteins were clustered into Branch c in the Bayesian clustering tree (*cf*. Figure 1, Table S6).

**Figure 9.**
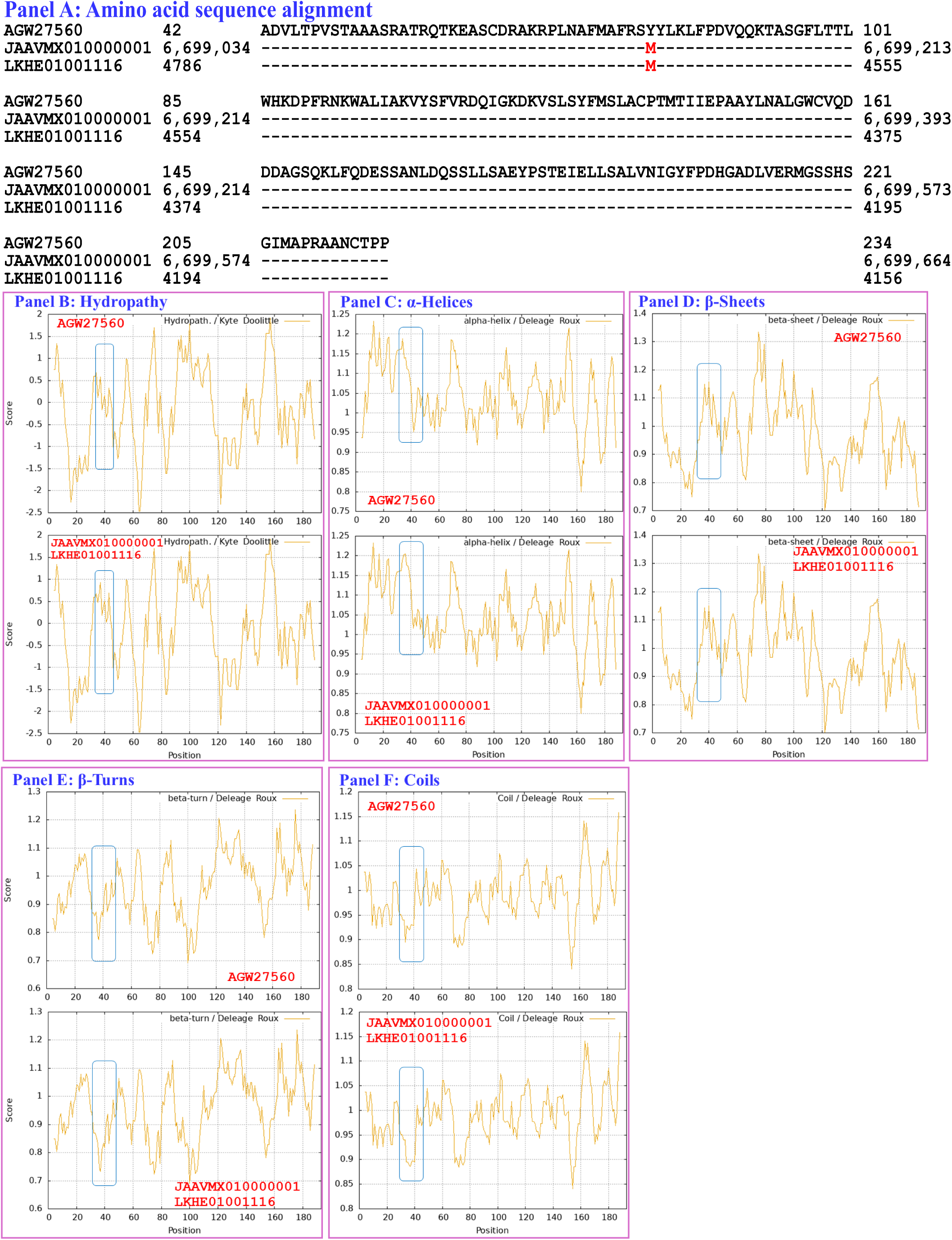
Correlation of the changes in the primary and secondary structures of the MATα_HMGbox domains of the MAT1-1-1 proteins: the reference protein AGW27560 is derived from the *H. sinensis* strain CS68-2-1229, and the truncated MAT1-1-1 proteins are encoded by the genome assemblies JAAVMX010000001 and LKHE01001116 of the *H. sinensis* strains IOZ07 and 1229, respectively. Panel A shows an alignment of the amino acid sequences in the MATα_HMGbox domains of the MAT1-1-1 proteins; amino acid substitutions are shown in red, whereas the hyphens indicate identical amino acid residues. The ExPASy ProtScale plots show the changes in hydrophobicity (Panel B) and in the 2D structures (Panels C−F show the α-helices, β-sheets, β-turns, and coils, respectively) of the proteins; the open rectangles in blue outline the changes in topology and waveform shown in the plots.

Compared with the reference MAT1-1-1-1 protein AGW27560 (*cf*. Figure S1, Table S6), no changes were detected in the amino acid sequences, hydrophobicities, or secondary structures (α-helices, β-sheets, β-turns, and coils) of the MATα_HMGbox domains of the MAT1-1-1 proteins encoded by the genome assembly ANOV01017390 (794←1228 & 1280←1369) of the *H. sinensis* strain Co18 or the metatranscriptome assembly OSIN7648 (151→675) of the *C. sinensis* insect‒fungal complex collected from Deqin, Yunnan, China [Kyte & Doolittle 1982; Bushley *et al*. 2013; Hu *et al*. 2013; Liu *et al*. 2015]. The MATα_HMGbox domains of the genome- and metatranscriptome-encoded MAT1-1-1 proteins clustered into Branch a1 in the Bayesian clustering tree (*cf*. Figure 1, Table S6).

The MATα_HMGbox domain of the MAT1-1-1 protein encoded by the metatranscriptome assembly GAGW01008880 (714←1127) of the *C. sinensis* insect‒fungi complex is truncated at its N-terminus, as shown in Figure S1 and in Panel A of Figure S3 [Xiang *et al*. 2014]. The remaining portion of the MATα_HMGbox domain of this protein is 100% identical to the query protein AGW27560 (under the AlphaFold code U3N942) derived from the *H. sinensis* strain CS68-2-1229 [Bushley *et al*. 2013], and it showed no apparent changes in hydrophobicity or secondary structure, as illustrated by the identical topology and waveform that can be observed in the ExPASy ProtScale plots for hydropathy, α-helices, β-sheets, β-turns, and coils shown in Panels B−F, respectively, of Figure S3. Although the MATα_HMGbox domain of the metatranscriptome-encoded protein clustered into Branch a1 in the Bayesian clustering tree together with the 6 reference authentic MAT1-1-1 proteins (*cf*. Figure 1, Table S6), the truncated MATα_HMGbox domain, which lacks 50 and 48 amino acid residues encoded by exons I and II of the *MAT1-1-1* gene, respectively, may not be able to form the core stereostructure containing 3 critical α-helices that supports high-affinity DNA binding and full functionality in the regulation of gene transcription related to the sexual reproduction of *O. sinensis* [Baxevanis *et al*. 1995; Thapar 2015; Li *et al*. 2025].

The genome assemblies LWBQ00000000 and NGJJ00000000 and the transcriptome assembly GCQL00000000 of the *H. sinensis* strains ZJB12195, CC1406-20395, and L0106, respectively, do not contain genes or transcripts that encode MAT1-1-1 proteins [Liu *et al*. 2015, 2020; Jin *et al*. 2020].

### III-7 Heteromorphic stereostructures of the HMG-box_ROX1-like domains of the MAT1-2-1 proteins derived from wild-type *C. sinensis* isolates

Section III-7 presents analyses of the correlation between the changes in the hydrophobicity and structure of the HMG-box_ROX1-like domains of full-length MAT1-2-1 proteins (under 12 AlphaFold codes, *cf*. Table S7) derived from wild-type *C. sinensis* isolates, compared to the reference MAT1-2-1 proteins represented by AEH27625 (under the AlphaFold code D7F2E9) derived from the *H. sinensis* strain CS2 (*cf*. Table S2) [Zhang *et al*. 2009]. The HMG-box_ROX1-like domain sequences (amino acid residues 127→197) were extended both upstream and downstream by 9 external amino acid residues because the sequential ExPASy ProtScale plotting for hydropathy, α-helices, β-sheets, β-turns, and coils was set up at a window size of 9 amino acids (*cf*. Section II-5 of Materials and Methods).

Figure 10 compares the hydrophobicities and structures of the HMG-box_ROX1-like domain of the MAT1-2-1 protein AIV43040 (under the AlphaFold code A0A0A0RCF5) derived from the *C. sinensis* isolate XZ12_16 (*cf*. Table S2) [Zhang *et al*. 2014] with those of the reference protein AEH27625 (under the AlphaFold code D7F2E9) derived from the *H. sinensis* strain CS2 [Zhang *et al*. 2009]. Panel A shows 3 amino acid substitutions: Y-to-H (aromatic tyrosine to basic histidine; hydropathy indices changed from - 1.3 to −3.2 [Kyte & Doolittle 1982]), M-to-I (methionine, which possesses polar neutral side chains, to aliphatic isoleucine; hydropathy indices changed from 1.9 to 4.5) and Q-to-R (glutamine, which has polar neutral side chains, to basic arginine; hydropathy indices changed from −3.5 to −4.5). Individually, these substitutions caused decreased or increased hydrophobicity, as illustrated by the changes in topology and waveform in the hydropathy plot shown in Panel B. These changes resulted in altered secondary structures surrounding the mutation sites in the HMG-box_ROX1-like domain of the MAT1-2-1 protein AIV43040, as illustrated by the changes in topology and waveform in the ExPASy ProtScale plots for α-helices, β-sheets, β-turns, and coils shown in Panels C−F, respectively. As shown in the representations of 3D structures in Panels G−H, the replaced residues are located in each of the 3 core α-helices, and they alter the stability of the hydrophobic core of the HMG-box_ROX1-like domain [Baxevanis *et al*. 1995; Thapar 2015] as well as the stereostructure of the domain of the MAT1-2-1 protein AIV43040 under the AlphaFold code A0A0A0RCF5. The mutant HMG-box_ROX1-like domain clustered into Branch b2β, which displays the longest clustering distance in the Bayesian clustering tree (*cf*. Figure 2, Table S7).

**Figure 10.**
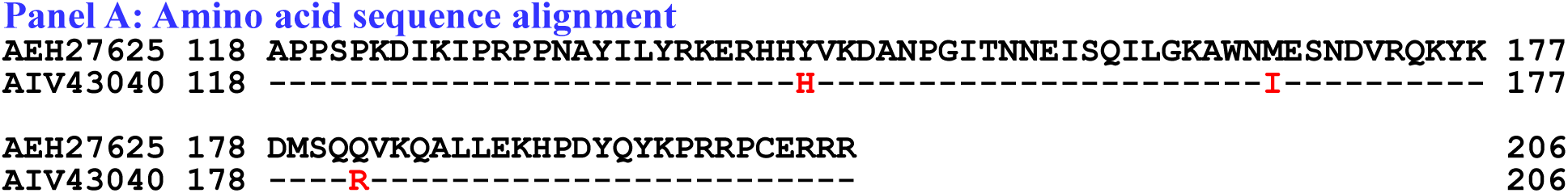
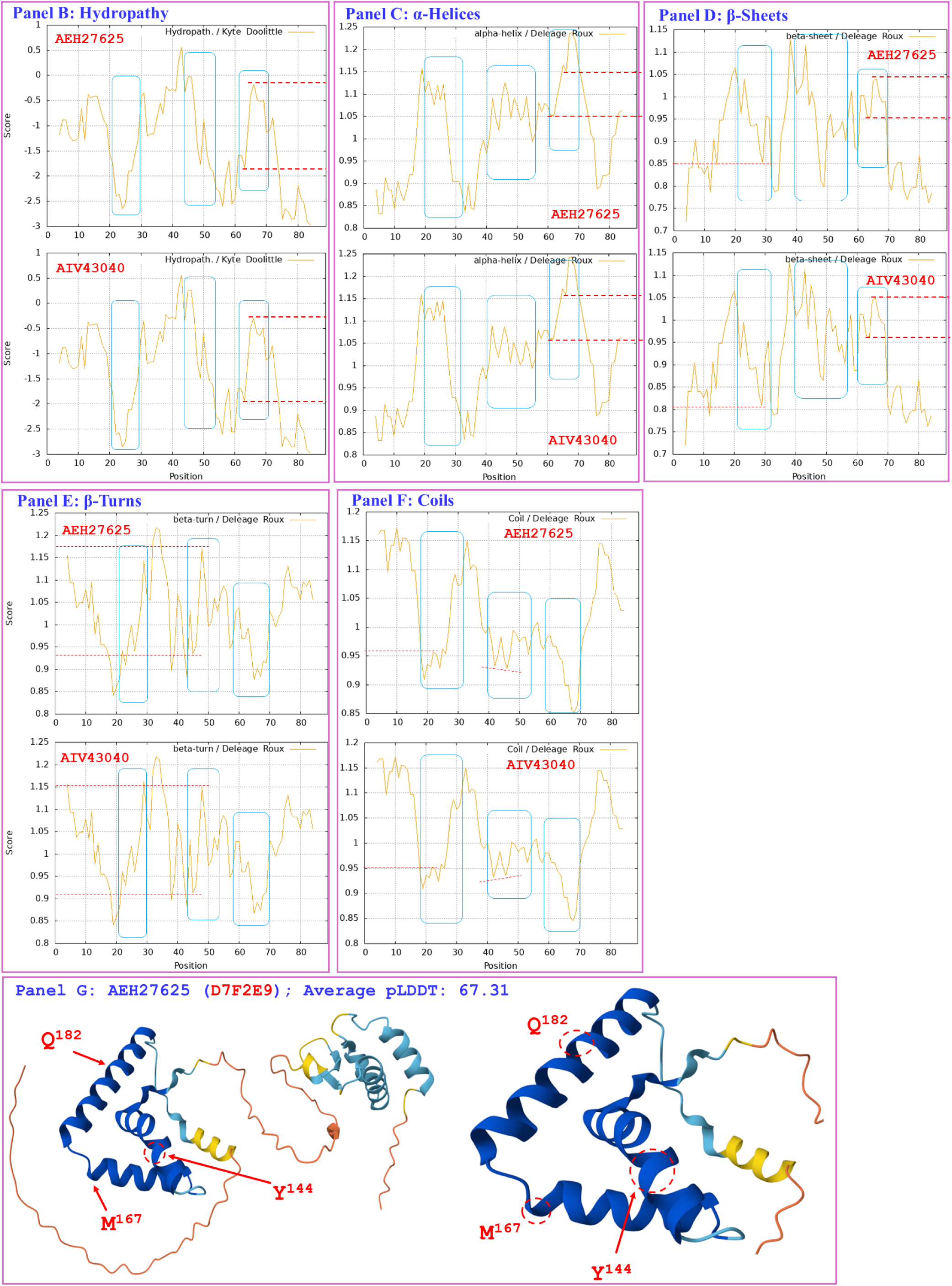
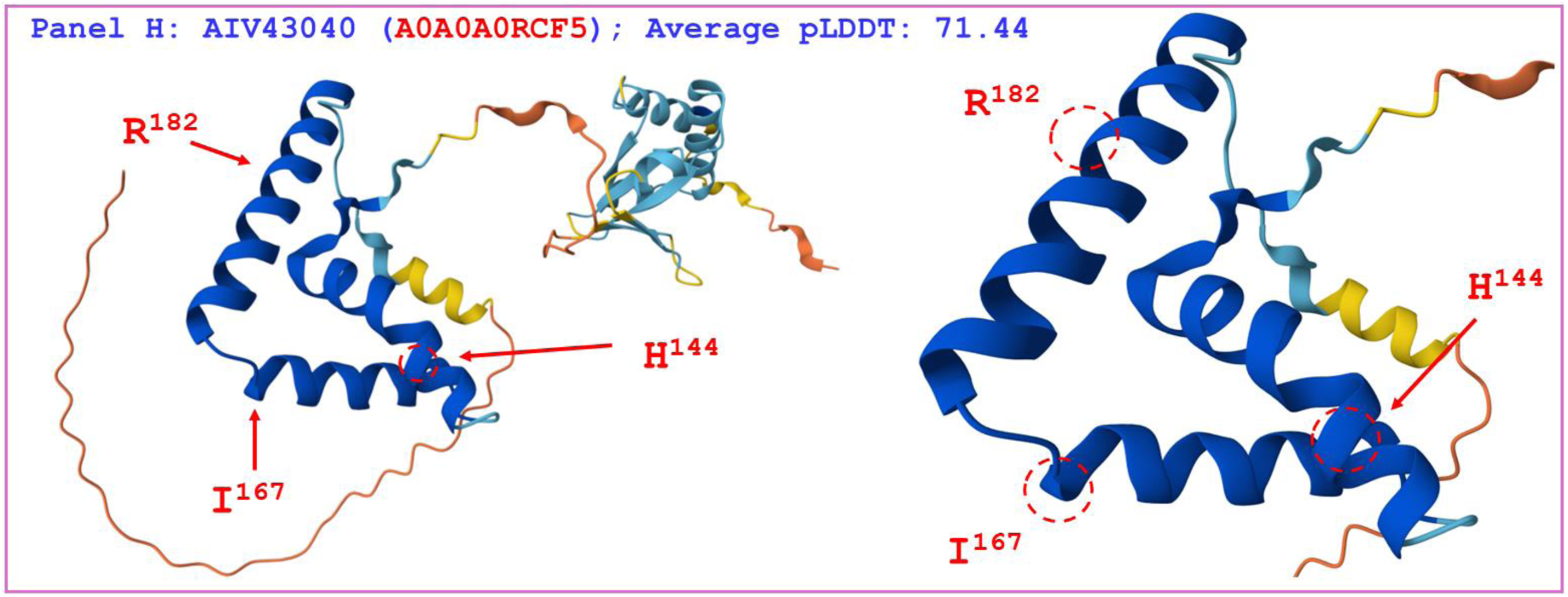
Correlation of the changes in the primary, secondary, and tertiary structures of the HMG-box_ROX1-like domains of MAT1-2-1 proteins. The reference protein AEH27625 (under the AlphaFold code D7F2E9) is derived from the *H. sinensis* strain CS2, and the mutant MAT1-2-1 protein AIV43040 (under the AlphaFold code A0A0A0RCF5) is derived from the wild-type *C. sinensis* isolate XZ12_16. Panel A shows an alignment of the amino acid sequences of the HMG-box_ROX1-like domains of the MAT1-2-1 proteins; amino acid substitutions are shown in red, whereas the hyphens indicate identical amino acid residues. The ExPASy ProtScale plots show the changes in hydrophobicity (Panel B) and in the 2D structure (Panels C−F show the α-helices, β-sheets, β-turns, and coils, respectively) of the protein; the open rectangles in blue outline the changes in topology and waveform seen in the plots. Panels G−H show representations of the 3D structures of the full-length proteins on the left; the locally amplified structures at the sites of substitutions are shown on the right. The model confidence for the AlphaFold-predicted 3D structures is indicated as follows: 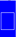 very high (pLDDT>90); 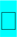 high (90>pLDDT>70); 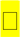 low (70>pLDDT>50); 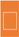 very low (pLDDT<50).

Figure 11 compares the hydrophobicities and structures of the HMG-box_ROX1-like domains of the MAT1-2-1 proteins under the AlphaFold code D7F2J7 (ACV60417, ACV60418, AFH35020, and AFX66443 shown in brown in Figure 11, derived from the wild-type *C. sinensis* isolates XZ-LZ07-H1, XZ-LZ07-H2, XZ06-124, and XZ05_8, respectively), and the hydrophobicities and structures of the proteins under the AlphaFold code D7F2F5 (ACV60375, ACV60415, AFX66441, AFX66446, and AFX66461 shown in green in Figure 11, derived from the wild-type *C. sinensis* isolates XZ-SN-44, XZ-LZ05-6, XZ05_2, XZ06_260, and XZ09_80, respectively) (*cf*. Table S2) [Zhang *et al*. 2009, 2012, 2014] with those of the reference protein AEH27625 (under the AlphaFold code D7F2E9) derived from the *H. sinensis* strain CS2 [Zhang *et al*. 2009]. Panel A shows substitutions of Y-to-H (aromatic tyrosine to basic histidine; hydropathy indices changed from −1.3 to −3.2 [Kyte & Doolittle 1982]) and M-to-I (methionine, which contains polar neutral side chains, to aliphatic isoleucine; hydropathy indices changed from 1.9 to 4.5). These substitutions caused changes in hydrophobicity, as illustrated by the changes in topology and waveform shown in the hydropathy plot in Panel B. They also resulted in altered secondary structures surrounding the mutation sites in the HMG-box_ROX1-like domains of the mutant MAT1-2-1 proteins ACV60417, ACV60418, AFH35020, and AFX66443 (under the AlphaFold code D7F2J7) and in those regions in the proteins ACV60375, ACV60415, AFX66441, AFX66446, and AFX66461 (under the AlphaFold code D7F2F5), as illustrated by the changes in topology and waveform shown in Panels C−F, which display the ExPASy ProtScale plots for α-helices, β-sheets, β-turns, and coils. As shown in the 3D structures presented in Panels G−I of Figure 11, the substituted residues are located within two of the 3 core α-helices, and they differentially alter the stability of the core structure of the HMG-box_ROX1-like domain [Baxevanis *et al*. 1995; Thapar 2015] and the tertiary structure of this domain in the MAT1-2-1 proteins under the AlphaFold 3D structural codes D7F2J7 and D7F2F5. The sequences of HMG-box_ROX1-like domain of the mutant proteins ACV60417, ACV60418, AFH35020, AFX66443, ACV60375, ACV60415, AFX66441, AFX66446, and AFX66461 clustered into Branch b2α in the Bayesian clustering tree (*cf*. Figure 2, Table S7).

**Figure 11.**
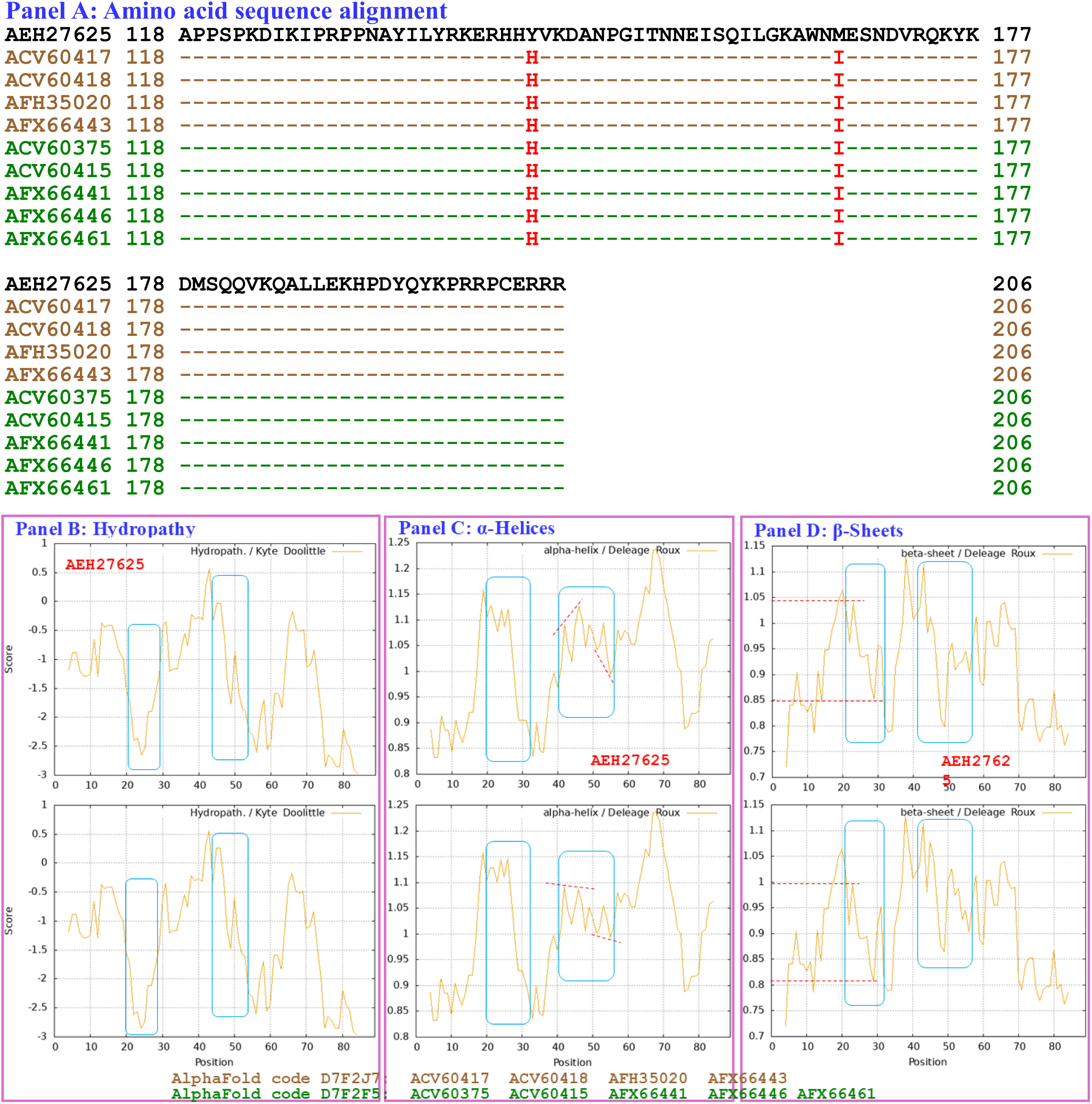
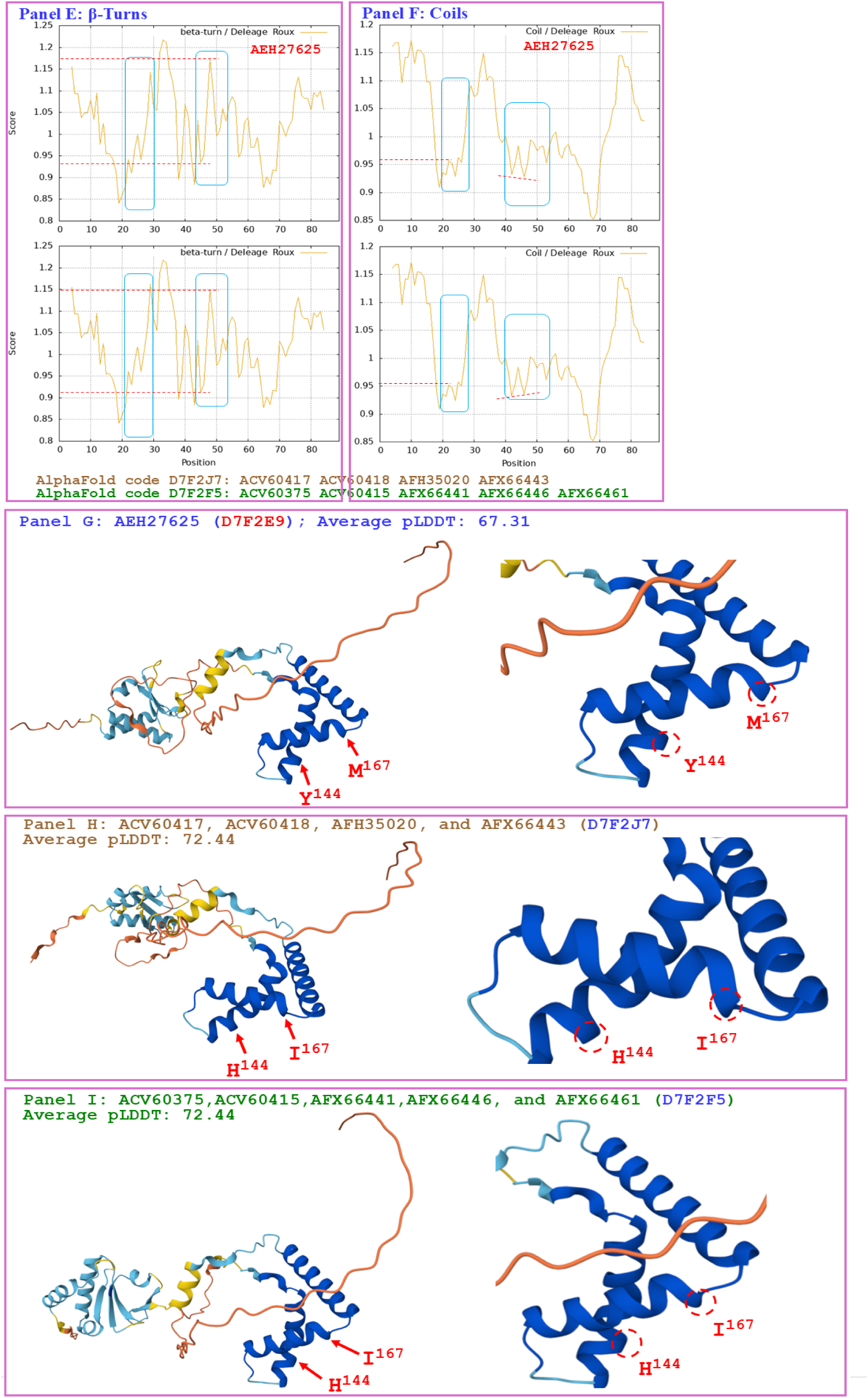
Correlation of the changes in the primary, secondary, and tertiary structures of the HMG-box_ROX1-like domains of MAT1-2-1 proteins. The reference protein AEH27625 (under the AlphaFold code D7F2E9) is derived from the *H. sinensis* strain CS2; the mutant MAT1-2-1 proteins under the AlphaFold code D7F2J7 (ACV60417, ACV60418, AFH35020, and AFX66443 shown in brown are derived from the wild-type *C. sinensis* isolate XZ-LZ07-H1, XZ-LZ07-H2, XZ06-124, and XZ05_8), and the mutant proteins under the AlphaFold code D7F2F5 (ACV60375, ACV60415, AFX66441, AFX66446, and AFX66461 shown in green are derived from the wild-type *C. sinensis* isolates XZ-SN-44, XZ-LZ05-6, XZ05_2, XZ06_260, and XZ09_80). Panel A shows an alignment of the amino acid sequences of the HMG-box_ROX1-like domains of the MAT1-2-1 proteins; amino acid substitutions are shown in red; the sequences displayed in brown and green represent the MAT1-2-1 proteins under the AlphaFold codes D7F2J7 and D7F2F5, respectively, and the hyphens indicate identical amino acid residues. The ExPASy ProtScale plots show the changes in hydrophobicity (Panel B) and in the 2D structures (Panels C−F for α-helices, β-sheets, β-turns, and coils, respectively) of the HMG-box_ROX1-like domains of the proteins; the open rectangles in blue outline the changes in topology and waveform seen in the plots. Panels G−H show representations of the 3D structures of the full-length proteins on the left and of the locally amplified structures at the sites of substitutions on the right. The model confidence for the AlphaFold-predicted 3D structures is indicated as follows: 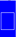 very high (pLDDT>90); 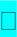 high (90>pLDDT>70); 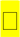 low (70>pLDDT>50); 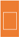 very low (pLDDT<50).

Figure 12 compares the hydrophobicities and structures of the HMG-box_ROX1-like domains of the MAT1-2-1 proteins under the AlphaFold codes **D7F2E3** (ACV60363, ACV60364, AFX66388, AFH35018, and AGW27542), **D7F2G5** (ACV60385), **V9LW71** (AFX66401), **V9LVS8** (AFX66472, AFX66473, and AFX66474), **V9LVU8** (AFX66475), **V9LWC9** (AFX66476), **V9LWG5** (AFX66484), and **U3N6V5** (AGW27537) derived from the wild-type *C. sinensis* isolates YN09_64, YN09_6, YN09_22, YN09_51, XZ-NQ-154, XZ-NQ-155, GS09_111, QH09-93, CS560-961, QH-YS-199, QH09_11, ID10_1, and CS6-251, respectively (*cf*. Table S2) [Zhang *et al*. 2009, 2012, 2014; Bushley *et al*. 2013], with the hydrophobicity and structure of the reference protein AEH27625 (under the AlphaFold code D7F2E9) derived from the *H. sinensis* strain CS2 [Zhang *et al*. 2009]. Panel A shows Y-to-H substitutions (aromatic tyrosine to basic histidine; hydropathy indices changed from −1.3 to −3.2 [Kyte & Doolittle 1982]) in the HMG-box_ROX1-like domains of all the mutant MAT1-2-1 proteins mentioned above; these substitutions caused reduced hydrophobicity, as illustrated by the changes in topology and waveform shown in the hydropathy plots in Panel B. The substitutions resulted in alteration of the secondary structure surrounding the mutation sites in the HMG-box_ROX1-like domains of the MAT1-2-1 proteins, as illustrated by the changes in topology and waveform in the ExPASy ProtScale plots for α-helices, β-sheets, β-turns, and coils that can be seen in Panels C−F, respectively.

**Figure 12.**
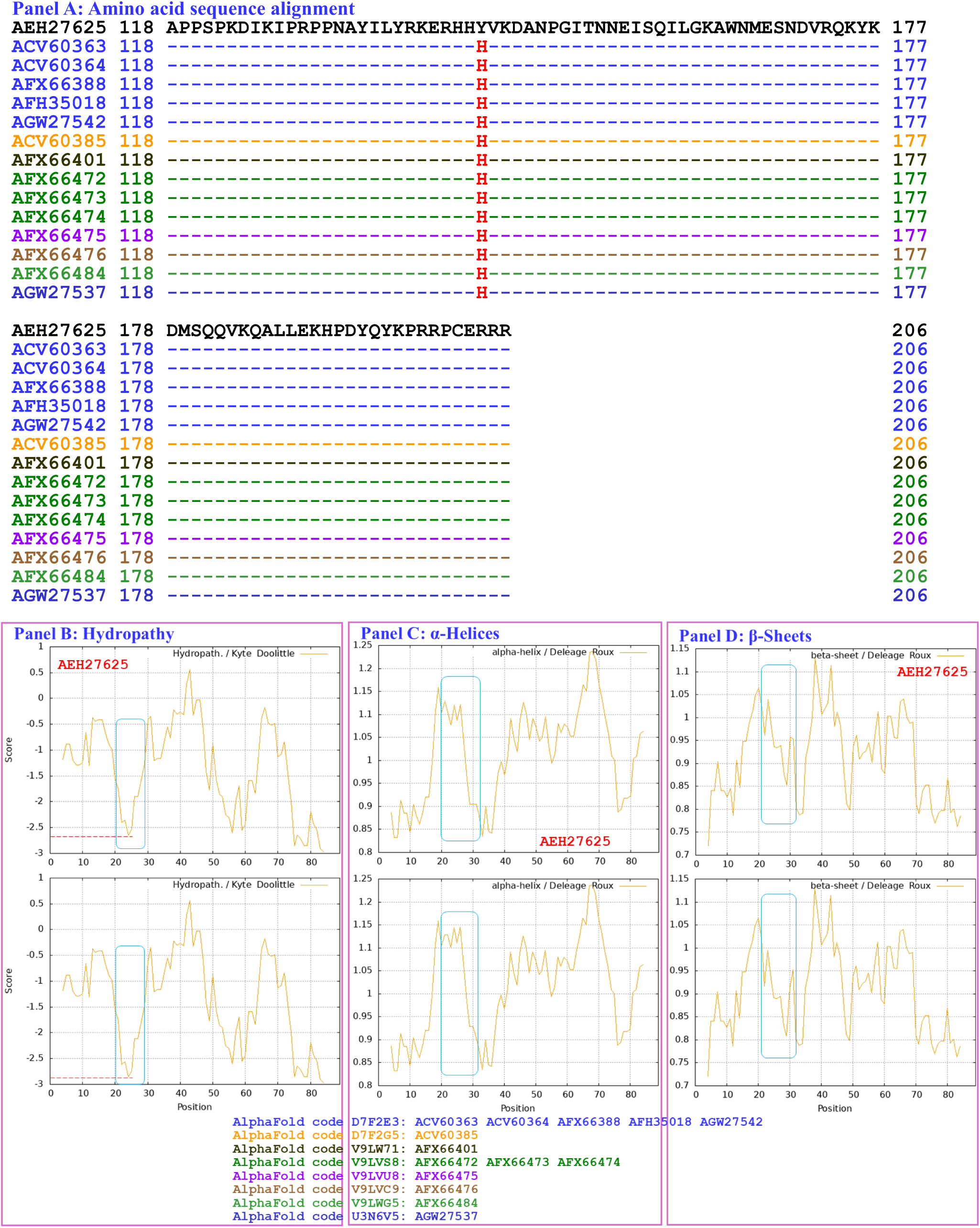
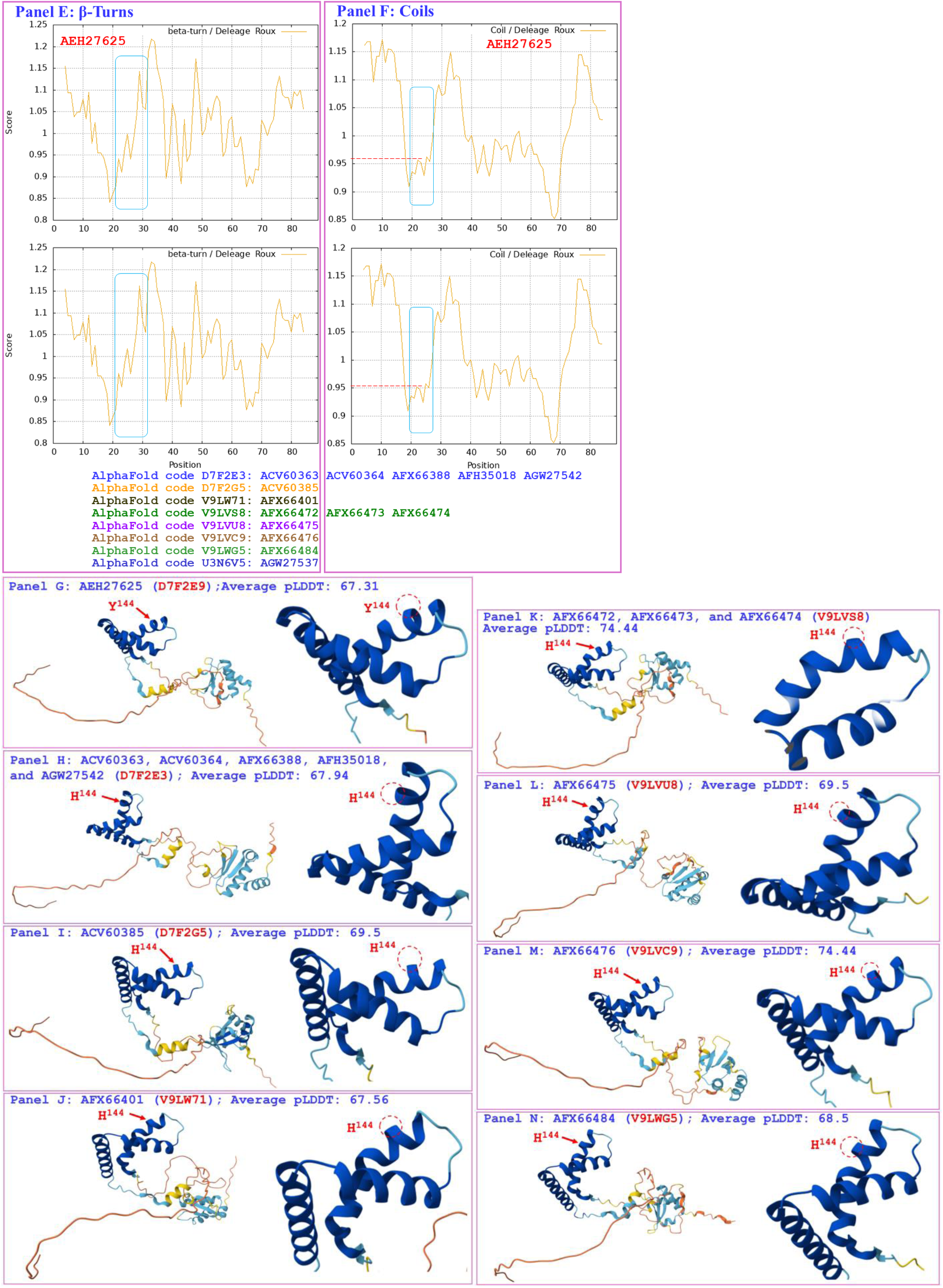
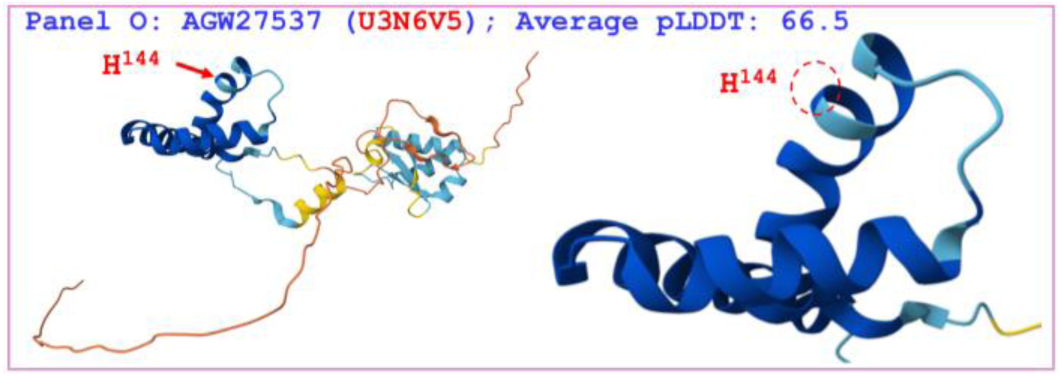
Correlation of the changes in the hydrophobicities and the primary, secondary, and tertiary structures of the HMG-box_ROX1-like domains of MAT1-2-1 proteins: the reference protein AEH27625 (under the AlphaFold code D7F2E9) is derived from the *H. sinensis* strain CS2, and the mutant MAT1-2-1 proteins under the AlphaFold codes **D7F2E3** (ACV60363, ACV60364, AFX66388, AFH35018, and AGW27542), **D7F2G5** (ACV60385), **V9LW71** (AFX66401), **V9LVS8** (AFX66472, AFX66473, and AFX66474), **V9LVU8** (AFX66475), **V9LWC9** (AFX66476), **V9LWG5** (AFX66484), and **U3N6V5** (AGW27537) are derived from the wild-type *C. sinensis* isolates YN09_64, YN09_6, YN09_22, YN09_51, XZ-NQ-154, XZ-NQ-155, GS09_111, QH09-93, CS560-961, QH-YS-199, QH09_11, ID10_1, and CS6-251, respectively. Panel A shows an alignment of the amino acid sequences of the HMG-box_ROX1-like domains of the MAT1-2-1 proteins; amino acid substitutions are shown in red, and the sequences depicted in various colors represent the MAT1-2-1 proteins under the AlphaFold codes V9LWC9, V9LVS8, D7F2E3, V9LVU8, D7F2G5, V9LW71, V9LWG5, and U3N6V5; the hyphens indicate identical amino acid residues. The ExPASy ProtScale plots show the changes in hydrophobicity (Panel B) and in the 2D structures (Panels C−F for α-helices, β-sheets, β-turns, and coils, respectively) of the proteins. The open rectangles in blue outline the changes in topology and waveform shown in the plots. Panels G−O show representations of the 3D structures of the full-length proteins on the left and the locally amplified structures at the mutation site on the right. The model confidence for the AlphaFold-predicted 3D structures is indicated as follows: 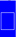 very high (pLDDT>90); 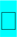 high (90>pLDDT>70); 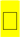 low (70>pLDDT>50);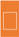 very low (pLDDT<50).

As shown in the representations of the tertiary structures presented in Panels G−O of Figure 12, the substituted residues with reduced hydrophobicity are located within the core α-helices in the HMG-box_ROX1-like domains, and they destabilize the hydrophobic core and alter the hydrophobic stereostructures of the domains [Baxevanis *et al*. 1995; Thapar 2015]. Although Panel A of Figure 12 shows the same Y-to-H substitution that caused the changes in hydrophobicity and second-order structure (shown in the ExPASy ProtScale plots in Panels B−F) in the HMG-box_ROX1-like domains of the MAT1-2-1 proteins ACV60363−ACV60364, ACV60385, AFH35018, AFX66388, AFX66472−AFX66476, AFX66401, AFX66484, AGW27537, and AGW27542, additional amino acid substitutions outside the HMG-box_ROX1-like domains (*cf*. Figure S2) might play synergistic roles in altering the tertiary structures of the proteins through long-range spatial interactions with the mutations within the domains (Panels H−O of Figure 12) of the MAT1-2-1 proteins under the 8 AlphaFold codes summarized in Table S7. The mutant HMG-box_ROX1-like domains clustered into Branch b1α in the Bayesian clustering tree (*cf*. Figure 2, Table S7).

Figure 13 compares the hydrophobicity and structure of the HMG-box_ROX1-like domain of the MAT1-2-1 protein ACV60399 (under the AlphaFold code D7F2H9) derived from the wild-type *C. sinensis* isolate SC-3 (*cf*. Table S2) [Zhang *et al*. 2009] with those of the HMG-box_ROX1-like domain of the reference protein AEH27625 (under the AlphaFold code D7F2E9) derived from the *H. sinensis* strain CS2 [Zhang *et al*. 2009]. Panel A shows substitutions of Y-to-H (aromatic tyrosine to basic histidine; hydropathy indices changed from −1.3 to −3.2 [Kyte & Doolittle 1982]) and Q-to-R (glutamine, which possesses polar neutral side chains, to basic arginine; hydropathy indices changed from −3.5 to −4.5); both of these substitutions caused reduced hydrophobicity, as illustrated by the changes in topology and waveform shown in the hydropathy plot in Panel B. These changes resulted in alteration of the secondary structure surrounding the mutation sites in the HMG-box_ROX1-like domain of the mutant MAT1-2-1 protein ACV60399, as illustrated by the changes in topology and waveform seen in the ExPASy ProtScale plots for α-helices, β-sheets, β-turns, and coils presented in Panels C−F. As shown in the representations of 3D structures in Panels G−H, the 2 replaced residues with reduced hydrophobicity are located inside and outside the hydrophobic core structure of the 3 α-helices, and they destabilize and alter the core stereostructure of the domain [Baxevanis *et al*. 1995; Thapar 2015]. The HMG-box_ROX1-like domain of the mutant protein ACV60399 under the AlphaFold code D7F2H9 clustered into Branch b1β in the Bayesian clustering tree (*cf*. Figure 2, Table S7).

**Figure 13.**
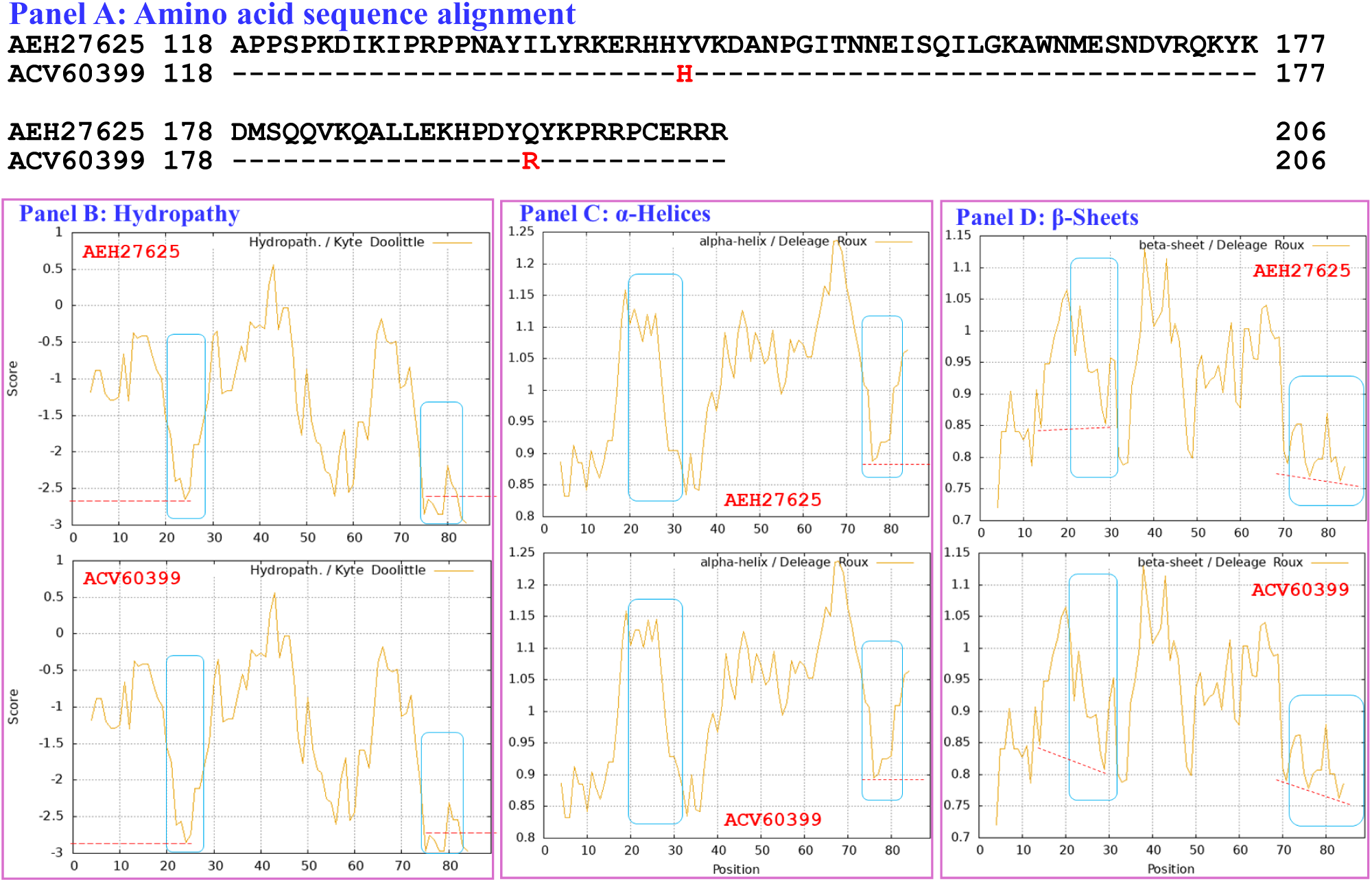
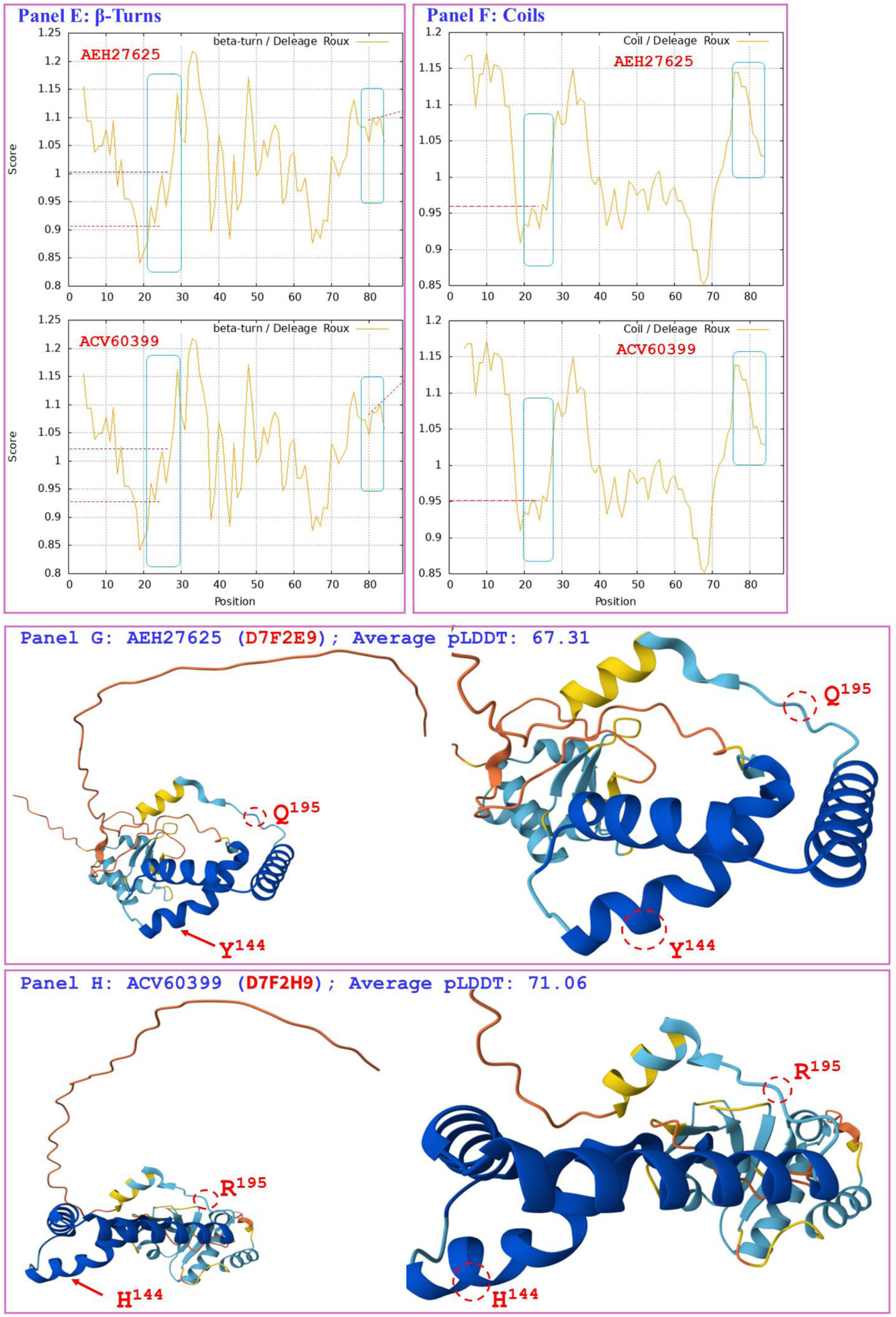
Correlation of the changes in the primary, secondary, and tertiary structures of the HMG-box_ROX1-like domains of MAT1-2-1 proteins: the reference protein AEH27625 (under the AlphaFold code D7F2E9) is derived from the *H. sinensis* strain CS2, and the mutant MAT1-2-1 protein ACV60399 (under the AlphaFold code D7F2H9) is derived from the wild-type *C. sinensis* isolate SC-3. Panel A shows an alignment of the amino acid sequences of the HMG-box_ROX1-like domains of the MAT1-2-1 proteins; amino acid substitutions are shown in red, whereas the hyphens indicate identical amino acid residues. The ExPASy ProtScale plots show the changes in hydrophobicity (Panel B) and in the 2D structure (Panels C−F for α-helices, β-sheets, β-turns, and coils) of the protein; the open rectangles in blue outline the changes in topology and waveform of the plots. Panels G−H show the 3D structures of the full-length proteins on the left and the locally amplified structures at the sites of substitutions on the right. The model confidence for the AlphaFold-predicted 3D structures is indicated as follows: 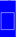 very high (pLDDT>90); 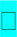 high (90>pLDDT>70); 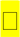 low (70>pLDDT>50); 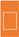 very low (pLDDT<50).

Aggregating the data presented in Section III-7, Panels A in Figures 10−13 demonstrate the 1−3 amino acid substitutions detected in the HMG-box_ROX1-like domains that caused changes in the topologies and waveforms shown in the ExPASy ProtScale plots presented in Panels B−F of the figures. These changes indicate the presence of altered hydrophobicity and secondary structures (α-helices, β-sheets, β-turns, and coils) of the HMG-box_ROX1-like domains of the mutant MAT1-2-1 proteins. These changes resulted in alterations in the stereostructures of the domains of the mutant proteins under different AlphaFold codes, as illustrated in Panel H of Figures 10−13, Panel I of Figure 11, Panels I−O of Figures 12 and summarized in Table S7. The mutant HMG-box_ROX1-like domains clustered into various clades in the Bayesian clustering tree (*cf*. Figure 2; Table S7) and ultimately change the DNA-binding specificities and affinities and the functionalities of gene transcriptional regulation, factors that play key roles in the sexual reproduction of *O. sinensis*.

### III-8 Diverse primary and secondary structures of the HMG-box_ROX1-like domains of MAT1-2-1 proteins encoded by the genome and transcriptome assemblies of *H. sinensis* and the metatranscriptome assembly of the *C. sinensis* insect‒fungal complex

Figure 14 shows a comparison of the hydrophobicity and structure of the HMG-box_ROX1-like domain of the MAT1-2-1 protein encoded by the genome assembly ANOV01000063 (9759→9851 & 9907→10,026) derived from *H. sinensis* strain Co18 [Hu *et al*. 2013] with the hydrophobicity and structure of the reference MAT1-2-1 protein AEH27625 derived from *H. sinensis* strain CS2 (under the AlphaFold code D7F2E9) [Zhang *et al*. 2009]. Panel A shows an S-to-A substitution (serine with polar neutral side chains to aliphatic alanine; hydropathy indices changed from −0.8 to 1.8 [Kyte & Doolittle 1982]). This substitution caused increased hydrophobicity surrounding the mutation site in the domain of the MAT1-2-1 protein encoded by the genome assembly ANOV01000063, as illustrated by the changes in topology and waveform changes seen in the hydropathy plot in Panel B.

**Figure 14.**
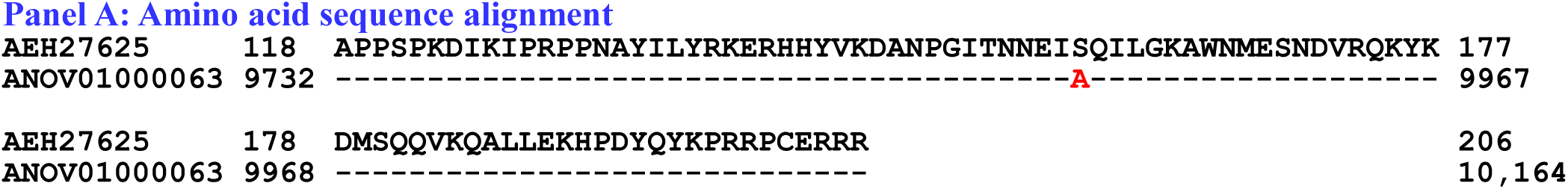
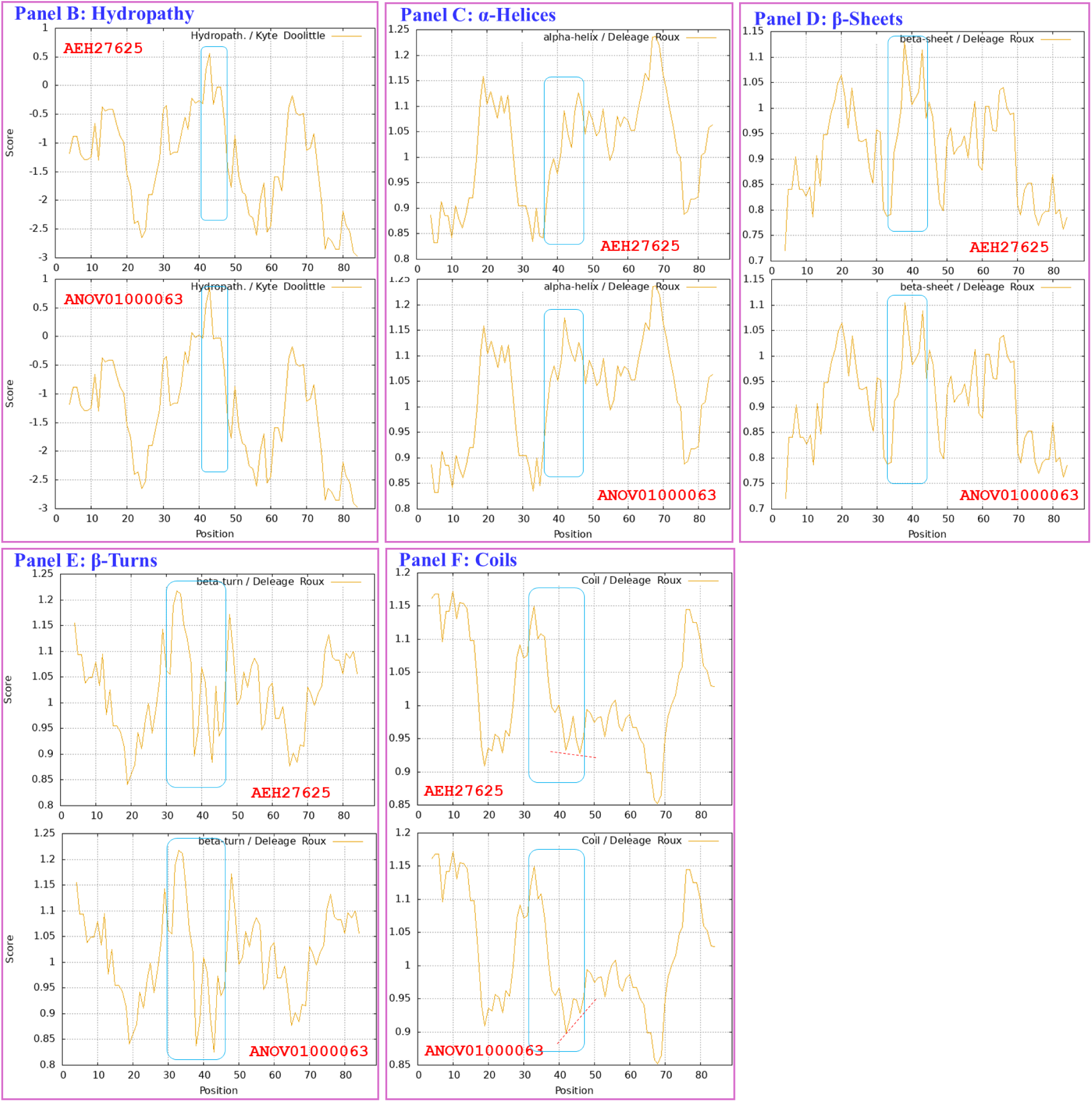
Correlation of the changes in the primary and secondary structures of the HMG-box_ROX1-like domains of the reference MAT1-2-1 protein AEH27625 (under the AlphaFold code D7F2E9) derived from the *H. sinensis* strain CS2 and the mutant MAT1-2-1 protein encoded by the genome assembly ANOV01000063 derived from the *H. sinensis* strain Co18. Panel A shows an alignment of the amino acid sequences of the HMG-box_ROX1-like domains of the MAT1-2-1 proteins; amino acid substitutions are shown in red, whereas the hyphens indicate identical amino acid residues. The ExPASy ProtScale plots show the changes in hydrophobicity (Panel B) and in the 2D structure (Panels C−F for α-helices, β-sheets, β-turns, and coils, respectively) of the protein; the open rectangles in blue outline the changes in topology and waveform of the plots.

These changes in the amino acid sequence and hydrophobicity resulted in altered secondary structure (α-helices, β-sheets, β-turns, and coils) within the HMG-box_ROX1-like domain of the mutant MAT1-2-1 protein encoded by the genome assembly ANOV01000063, as illustrated in the ExPASy ProtScale plots for α-helices, β-sheets, β-turns, and coils shown in Panels C−F, respectively, in Figure 14 [Baxevanis *et al*. 1995; Thapar 2015]. These changes in the sequences and hydrophobicity may have altered the tertiary structure of the HMG-box_ROX1-like domain of the genome-encoded protein. The altered HMG-box_ROX1-like domain clustered into Branch a2 in the Bayesian clustering tree (*cf*. Figure 2; Table S8).

Figure 15 compares the hydrophobicities and structures of the HMG-box_ROX1-like domains of the MAT1-2-1 proteins encoded by the genome assemblies LKHE01001605 (14,016←14,135 & 14,191←14,283), LWBQ01000021 (14,016←14,135 & 14,191←14,283), and NGJJ01000619 (23,186←23,305 & 23,361←23,453) of the *H. sinensis* strains 1229, ZJB12195, and CC1406-20395, respectively [Li *et al*. 2016a; Jin *et al*. 2020; Liu *et al*. 2020], with the hydrophobicity and structure of the reference protein AEH27625 (under the AlphaFold code D7F2E9) derived from the *H. sinensis* strain CS2 [Zhang *et al*. 2009]. Panel A shows Y-to-H (aromatic tyrosine to basic histidine; hydropathy indices changed from −1.3 to −3.2 [Kyte & Doolittle 1982]) and S-to-A substitutions (serine, which has polar neutral side chains, to aliphatic alanine; hydropathy indices changed from −0.8 to 1.8), which resulted in altered hydrophobicity, as illustrated in the hydropathy plot in Panel B. These changes altered the secondary structures surrounding the mutation sites in the HMG-box_ROX1-like domains of the mutant MAT1-2-1 proteins encoded by the genome assemblies LKHE01001605, LWBQ01000021, and NGJJ01000619, as illustrated in the ExPASy ProtScale plots for α-helices, β-sheets, β-turns, and coils shown in Panels C−F, respectively [Baxevanis *et al*. 1995; Thapar 2015], and subsequently altered the tertiary structures of the core stereostructures of the domains. The mutant HMG-box_ROX1-like domains of the genome-encoded proteins clustered into Branch b3 in the Bayesian clustering tree (*cf*. Figure 2, Table S8).

**Figure 15.**
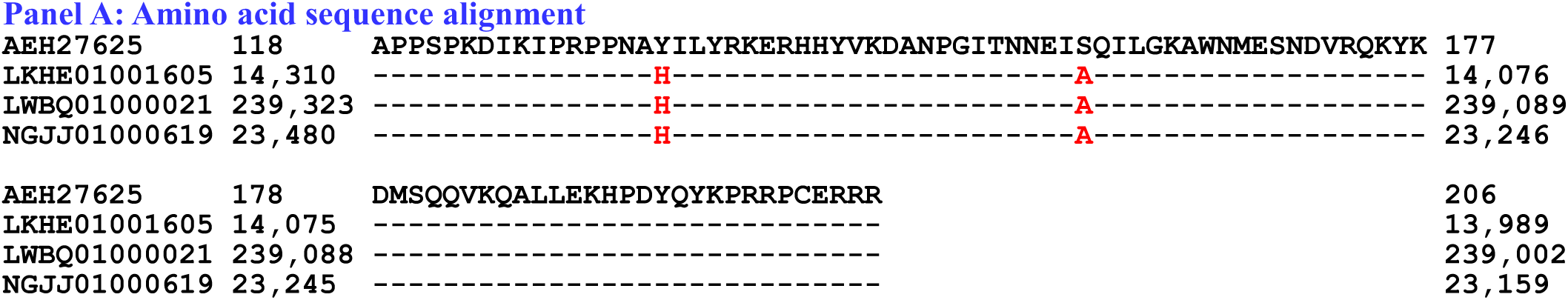
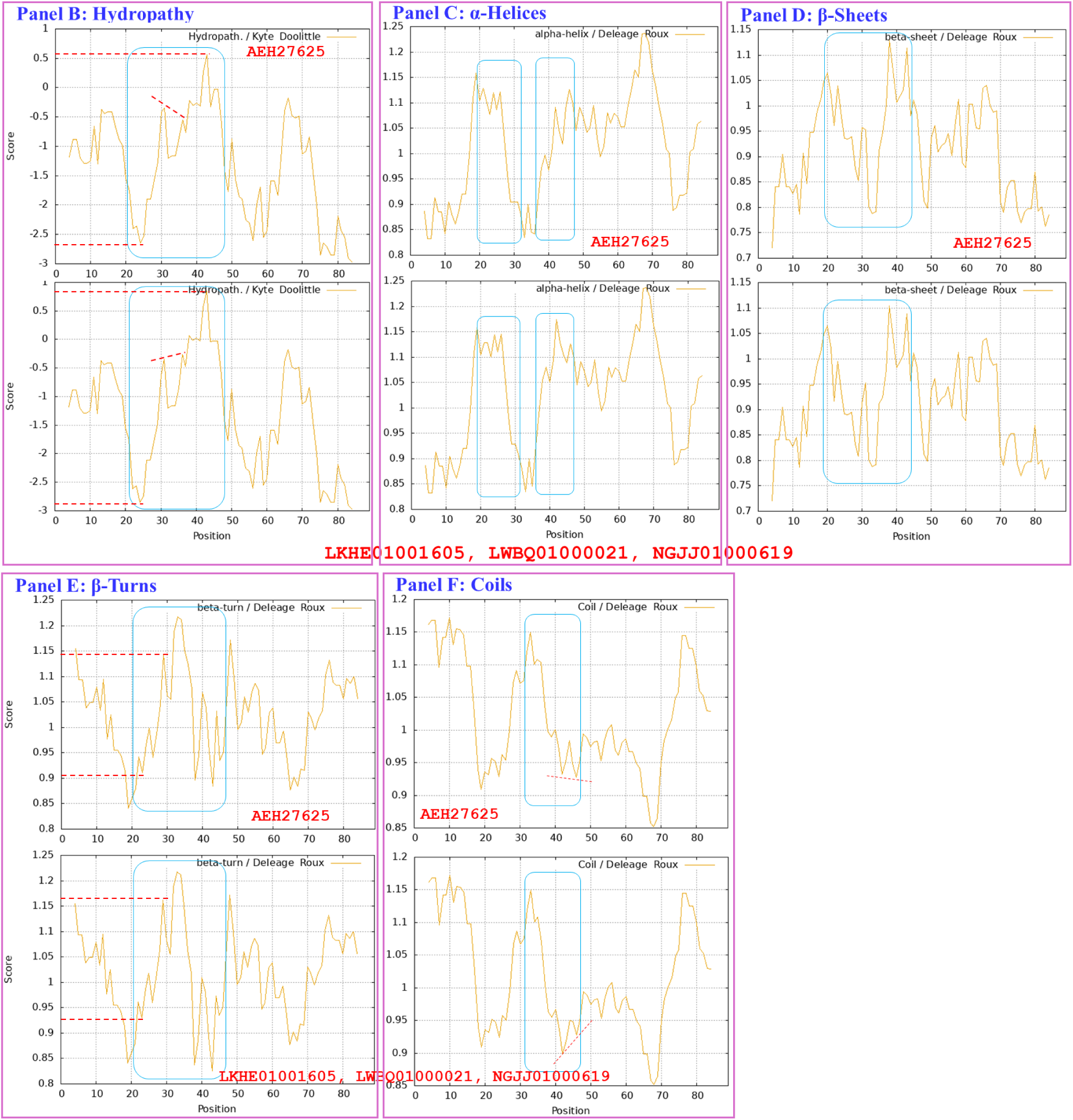
Correlation of the changes in the primary and secondary structure of the HMG-box_ROX1-like domain of the reference MAT1-2-1 protein AEH27625 (under the AlphaFold code D7F2E9) derived from the *H. sinensis* strain CS2 in and the mutant MAT1-2-1 proteins encoded by the genome assemblies LKHE01001605, LWBQ01000021, and NGJJ01000619 derived from *H. sinensis* strains 1229, ZJB12195, and CC1406-20395, respectively. Panel A shows an alignment of the amino acid sequences of the HMG-box_ROX1-like domains of the MAT1-2-1 proteins; amino acid substitutions are shown in red, whereas the hyphens indicate identical amino acid residues. The ExPASy ProtScale plots show the changes in hydrophobicity (Panel B) and in the 2D structures (Panels C−F for α-helices, β-sheets, β-turns, and coils, respectively) of the proteins; the open rectangles in blue outline the changes in topology and waveform seen in the plots.

The genome assembly JAAVMX000000000 of the *H. sinensis* strain IOZ07 does not contain a gene encoding the MAT1-2-1 protein [Shu *et al*. 2020].

Figure 16 compares the hydrophobicity and structure of the HMG-box_ROX1-like domain of the MAT1-2-1 proteins encoded by the transcriptome assembly GCQL01020543 (553←765) of the *H. sinensis* strain L0106 and by the metatranscriptome assembly OSIN7649 (379→591) of the mature *C. sinensis* insect‒fungal complex [Liu *et al*. 2015; Xia *et al*. 2017] with the hydrophobicity and structure of the reference protein AEH27625 (under the AlphaFold code D7F2E9) derived from the *H. sinensis* strain CS2 [Zhang *et al*. 2009]. Panel A shows Y-to-H substitutions (aromatic tyrosine to basic histidine; hydropathy indices changed from −1.3 to −3.2 [Kyte & Doolittle 1982]). The substitutions caused decreased hydrophobicity, as illustrated by the changes in topology and waveform seen in the hydropathy plot in Panel B. These changes resulted in alterations of the secondary structure surrounding the mutation sites in the HMG-box_ROX1-like domains of the MAT1-2-1 proteins encoded by the transcriptome assembly GCQL01020543 and the metatranscriptome assembly OSIN7649, as illustrated by the changes in topology and waveform shown in the ExPASy ProtScale plots for α-helices, β-sheets, β-turns, and coils in Panels C−F.

**Figure 16.**
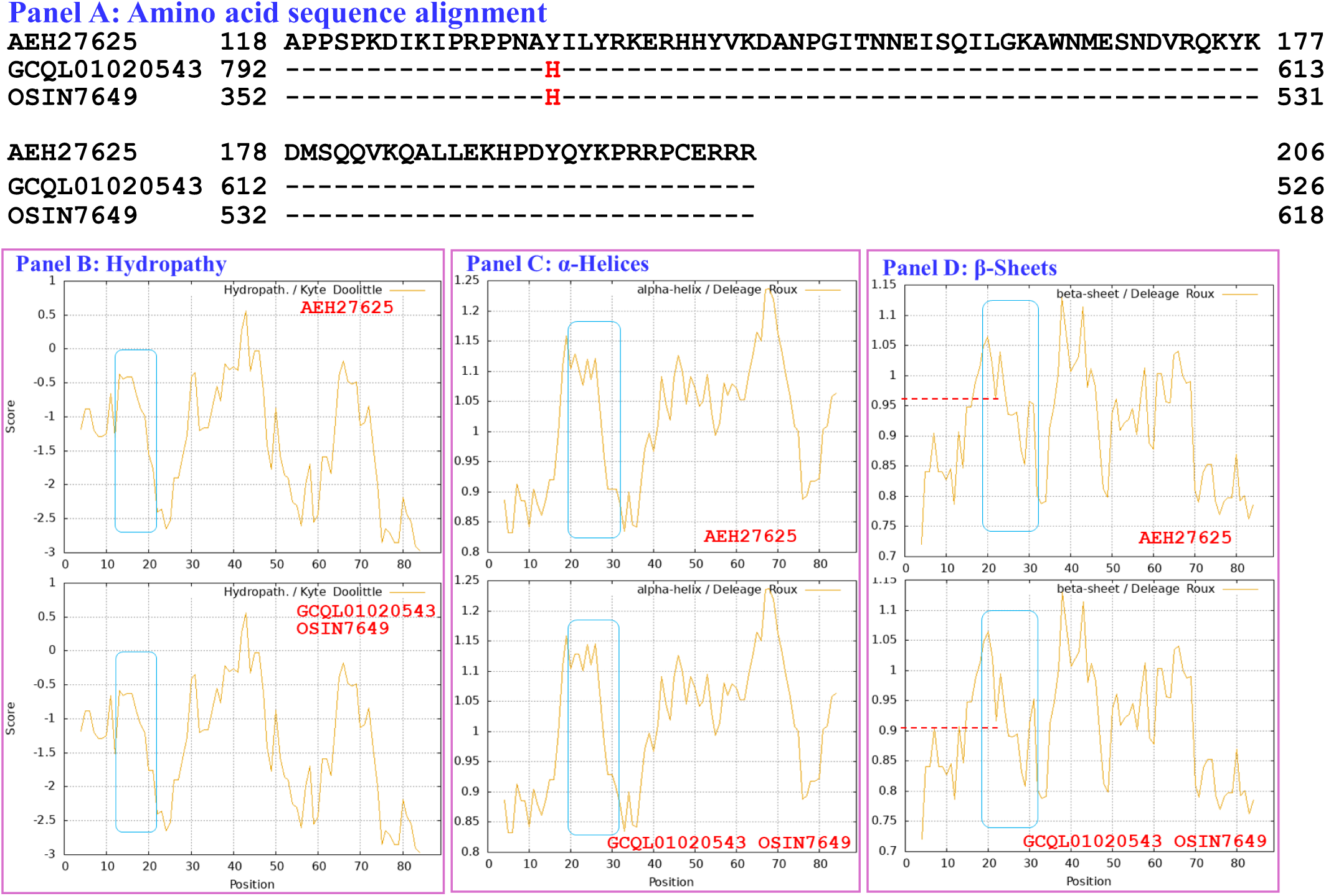
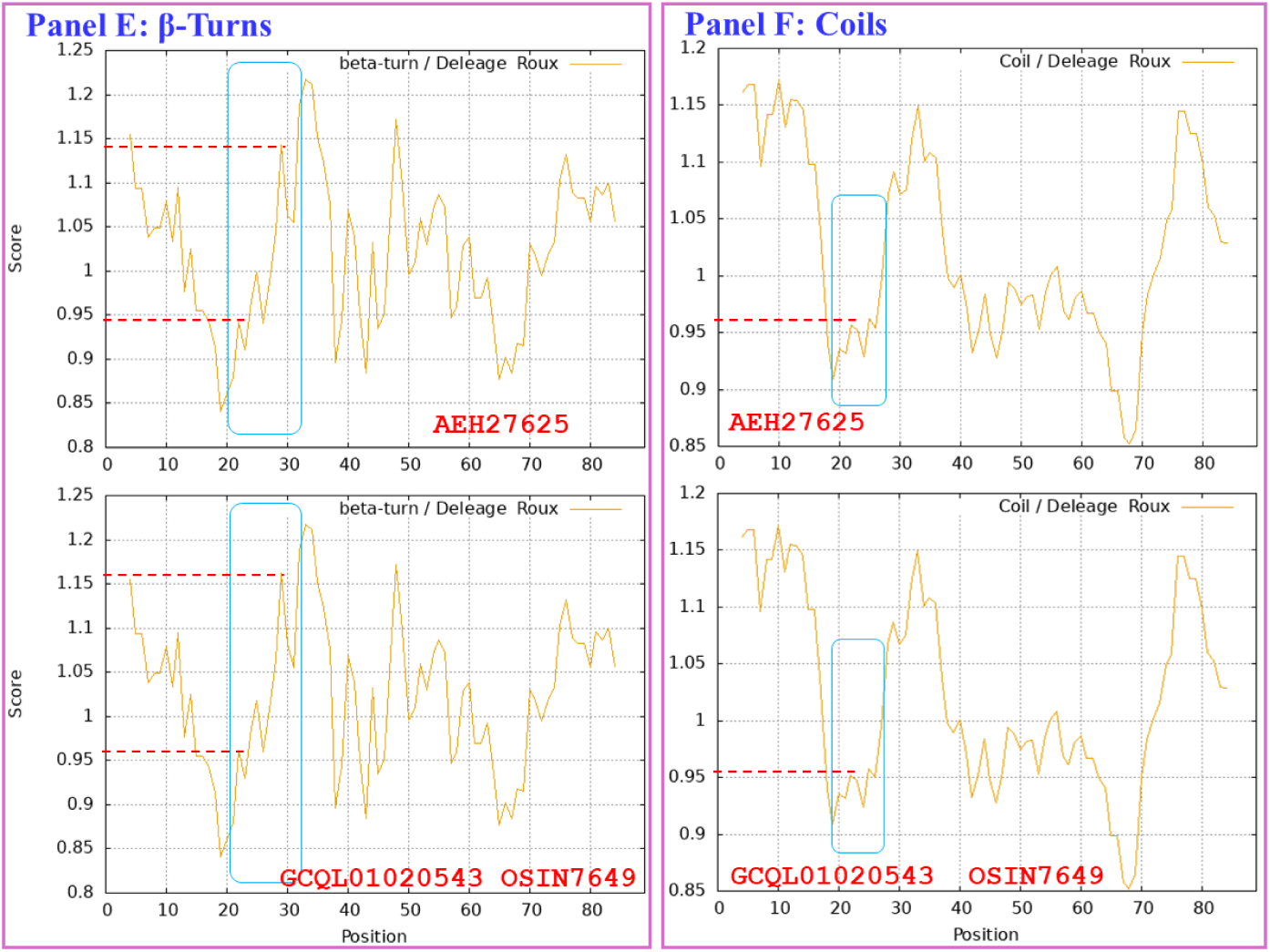
Correlation of the primary and secondary structure of the HMG-box_ROX1-like domain of the reference MAT1-2-1 protein AEH27625 (under the AlphaFold code D7F2E9) derived from the *H. sinensis* strain CS2 with the primary and secondary structures of the mutant MAT1-2-1 proteins encoded by the transcriptome assembly GCQL01020543 derived from the *H. sinensis* strain L0106 and the metatranscriptome assembly OSIN7649 (379→591) of the mature C. sinensis insect‒fungal complex. Panel A shows an alignment of the amino acid sequences of the HMG-box_ROX1-like domains of the MAT1-2-1 proteins; amino acid substitutions are shown in red, whereas the hyphens indicate identical amino acid residues. The ExPASy ProtScale plots show the changes in hydrophobicity (Panel B) and in the 2D structures (Panels C−F for α-helices, β-sheets, β-turns, and coils, respectively) of the proteins; the open rectangles in blue outline the changes in topology and waveform seen in the plots.

Similar to the Y-to-H substitutions in the HMG-box_ROX1-like domains of the mutant MAT1-2-1 proteins under various AlphaFold 3D structural models (such as V9LWC9, V9LVS8, D7F2E3, V9LVU8, D7F2G5, V9LW71, V9LWG5, and U3N6V5) shown in Figure 12, the changes in hydrophobicities and primary and secondary structures shown in Figure 16 may subsequently alter the tertiary structures of the HMG-box_ROX1-like domains of the transcriptome-encoded and metatranscriptome-encoded mutant proteins. The mutant HMG-box_ROX1-like domains of the transcriptome- and metatranscriptome-encoded proteins clustered into Branch b1α in the Bayesian clustering tree (*cf*. Figure 2, Table S8).

### III-9 Heterogenous fungal sources of the MAT1-1-1 and MAT1-2-1 proteins

Although wild-type *C. sinensis* isolates are often considered impure as *H. sinensis*, analysis of mutant full-length mating proteins revealed their heterogeneous fungal sources, as shown in Table S3. Of the wild-type *C. sinensis* isolates whose full-length MAT1-1-1 and MAT1-2-1 proteins were analyzed (*cf*. Figures 3−8 and 10−13), 34 had ITS genotyping information in the GenBank database and belonged to GC-biased Genotypes #1 and #3 of genome-independent *O. sinensis* fungi (Table S3) [Zhu & Li 2017; Li *et al*. 2022]. Although the ITS sequences of 29 of the 34 wild-type *C. sinensis* isolates appeared to show high homology (≥97%) to the ITS sequences of GC-biased Genotype #1 *H. sinensis*, the mutant MAT1-1-1 and MAT1-2-1 proteins might not necessarily be derived from GC-biased Genotype #1 *H. sinensis* within the impure fungal complex containing more than one fungal species [Li *et al*. 2016b], while the *H. sinensis* ITS sequences were most easily amplified during regular simple-step PCR under the experimental settings, unless the PCR experiments using multiple pairs of primers and combining with amplicon cloning techniques with selection and sequencing of sufficient (usually 30−50) white colonies. In other words, because of the impure nature of the wild-type *C. sinensis* isolates [Li *et al*. 2016b, 2022; Zhu & Li 2017], the mutant proteins may be derived from cooccurring unvalidated heterospecific or genotypic fungi.

## IV. Discussion

### IV-1 Correlation of the hydrophobicity and the primary, secondary, and tertiary structures of the DNA-binding domains of the MAT1-1-1 and MAT1-2-1 proteins

The MATα_HMGbox domain of the MAT1-1-1 protein and the HMG-box_ROX1-like domain of the MAT1-2-1 protein are distinct DNA-binding domains that are critical for fungal mating-type regulation but differ in their evolutionary lineage, structural architecture, and functional roles [Jackson *et al*. 2013; Zheng *et al*. 2013; Kim *et al*. 2015]. The HMGbox domains in mating proteins are multifunctional motifs central to the transcriptional regulation of mating-related genes, *O. sinensis* sexual reproduction, and host adaptation in the *C. sinensis* insect‒fungal complex. Our study demonstrates that various 1−3 amino acid substitutions within the MATα_HMGbox and HMG-box_ROX1-like domains of the MAT1-1-1 and MAT1-2-1 proteins, respectively, of the wild-type *C. sinensis* isolates and *C. sinensis* insect‒fungal complexes result in changes in the hydrophobic properties and in the secondary and tertiary structures of the DNA-binding domains of the proteins (*cf*. Figures 3−16). The variable sequences of the DNA-binding domains clustered into different clades in the Bayesian clustering trees (*cf*. Figures 1−2; Tables S5−S8).

The MATα_HMGbox domain of the MAT1-1-1 protein contains a core structure of 3 α-helices that ensures high-affinity, sequence-dependent binding to AT-rich DNA motifs [Ramšak *et al*. 2020, 2021; Ramšak & Kück 2022]. This binding facilitates the assembly of transcriptional machinery in a way that involves mating-type-specific genes and activates gene transcription by inducing DNA conformational changes through physical bending or twisting of the target DNA segments [Martin *et al*. 2010; Ait Benkhali *et al*. 2013; Ramšak & Kück 2022]. The domain participates directly in promoter recognition of specific genes related to the heterothallic reproductive process [Yamamoto *et al*. 1997].

Table S4 shows the hydropathy values for amino acids that are defined by their side chains according to Kyte & Doolittle [1982] (provided by https://web.expasy.org/protscale/). Larger hydropathy values indicate stronger hydrophobicity, while negative values indicate hydrophilicity [Kyte & Doolittle 1982; Baxevanis *et al*. 1995; Thapar 2015]. The core α-helical stereostructures of the MATα_HMGbox domains are stabilized through a hydrophobic core that is composed of nonpolar amino acids such as isoleucine (Ile, I), valine (Val, V), leucine (Leu, L), phenylalanine (Phe, F), cysteine (Cys, C), methionine (Met, M), and alanine (Ala, A); these amino acids have hydropathy indices of 4.5, 4.2, 3.8, 2.8, 2.5, 1.9, and 1.8, respectively (*cf*. Table S4) [Kyte & Doolittle 1982].

The HMG-box_ROX1-like domain of the MAT1-2-1 protein also consists of 3 asymmetrically arranged α-helices that are connected by flexible loops, forming a key hydrophobic core that is stabilized through the presence of the aforementioned nonpolar amino acids (*cf*. Table S4). The HMG-box_ROX1-like domain of the MAT1-2-1 protein regulates gene expression in a sequence-specific manner by binding to ATTAAT or ATTGTT motifs [Jackson *et al*. 2013; Zheng *et al*. 2013; Kim *et al*. 2015]. This domain has been implicated in chromatin remodeling and transcriptional repression through its interaction with other corepressors, similar to the repressor of oxygen-regulated genes 1 (ROX1) domain in yeast, through opposing pathways to the MATα_HMGbox domain of the MAT1-1-1 protein [Balasubramanian *et al*. 1993; Zitomer *et al*. 1997; Zheng *et al*. 2013]. ROX1 represses its own promoter, creating a feedback loop for hypoxic response regulation as a central hypoxia-responsive regulator that plays roles in chromatin remodeling following DNA binding and in the integration of metabolic signals [Kastaniotis & Zitomer 2000].

To date, no studies have directly addressed heterodimerization between the MAT1-1-1 and MAT1-2-1 proteins of *O. sinensis*, although formation of heterodimers between the MATα_HMGbox domain of the MAT1-1-1 protein and the HMG-box_ROX1-like domain of the MAT1-2-1 protein is believed to be a critical mechanism in the regulation of fungal mating, particularly in heterothallic ascomycetes [Kües & Casselton 1993; Asante-Owusu *et al*. 1996; Jacobsen *et al*. 2002]. However, the MATα_HMGbox and HMG-box_ROX1-like domains likely interact with each other, enabling them to cooperate synergistically in DNA binding through their complementary electrostatic surfaces [Hancock *et al*. 2019]. The heteromorphic stereostructures of the HMGbox domains of the mutant mating proteins derived from wild-type *C. sinensis* isolates presented in this study likely alter the complementary cooperation of the mating proteins of the self-sterile *O. sinensis* under heterothallic or hybrid outcrossing and affect the subsequent functionalities of activation or repression of mating-related gene transcription [Li *et al*. 2023c, 2024b, 2025].

### IV-2 Heterogenous fungal sources of the MAT1-1-1 and MAT1-2-1 proteins

The analysis of mutant full-length mating proteins derived from wild-type *C. sinensis* isolates (Figures 3−8 and 10−13) revealed their heterogeneous fungal sources (Table S3) [Zhang *et al*. 2009, 2011, 2012, 2013, 2014; Zhang & Zhang 2015]. A total of 34 wild-type *C. sinensis* isolates were examined for molecular taxonomy ITS sequences that are available in GenBank and were found to belong to genomically independent GC-biased Genotypes #1 and #3 of *O. sinensis* fungi. However, whether the impure wild-type *C. sinensis* isolates cooccur with other heterospecific fungi remains unclear, which remains to be determined through the use of culture-dependent techniques or culture-independent PCR protocols involving the use of amplicon cloning techniques with the selection and sequencing of sufficient amounts (usually ≥30) of white colonies.

Li et al. [2016b] conducted a similar *C. sinensis* isolate study and reported 2 other wild-type *C. sinensis* isolates, CH1 and CH2. With using multiple pairs of PCR and rigorous amplicon cloning techniques, these isolates were proven to coexist GC-biased Genotype #1 and AT-biased Genotypes #4−5 of *O. sinensis*, and heterospecific *Paecilomyces hepiali*. Li et al. [2022. 2023c, 2023d, 2024b] further proved that the ITS sequences of Genotypes #2−17 of *O. sinensis* with multiple transition and transversion point mutations are not the repetitive copies in the genome of GC-biased Genotype #1 *H. sinensis* but belong to independent *O. sinensis* fungi, regardless of whether they are GC- or AT-biased. Although displaying the key genetic features of fungal complexes, these wild-type isolates phenotypically exhibited *in vitro* growth and microscopic morphological characteristics of psychrophilic *H. sinensis* in phenotype. Functionally, in the inoculation experiments on larvae of *Hepialus armoricanus* (n=100 larvae each study group), the infection rate was significantly increased 15- to 39-fold to 55.2% for the wild-type isolates CH1 and CH2 from nearly non-infectious 1−3% for *H. sinensis* (*P*<0.001) and the larval death latency was largely shortened to 5−8 days for the CH1 and CH2 from 35−50 days for *H. sinensis*.

### IV-3 The MAT1-1-1 and MAT1-2-1 proteins are involved in the sexual reproduction of *O. sinensis*

The MAT1-1-1 and MAT1-2-1 proteins coregulate mating compatibility and sexual development during the sexual reproduction of *O. sinensis*, a process in which the MATα_HMGbox and HMG-box_ROX1-like domains play key roles. The hypothesis of self-fertilization via a homothallism or psuedohomothalism strategy has been proposed for *O. sinensis* [Bushley *et al*. 2013; Hu *et al*. 2013] on the basis of genetic research. However, Zhang and Zhang [2015] argued against this hypothesis and suggested that *O. sinensis* uses facultative hybridization in its sexual reproduction on the basis of findings of differential occurrence of the *MAT1-1-1* and *MAT1-2-1* genes in more than 170 wild-type *C. sinensis* isolates derived from natural *C. sinensis* insect‒fungal complex specimens collected from different production areas on the Qinghai‒Tibet Plateau. Furthermore, Li *et al*. [2023c, 2024b, 2025] reported differential occurrence, differential translation, and alternative splicing of the *MAT1-1-1* and *MAT1-2-1* genes and heteromorphic stereostructures of the MAT1-1-1 and MAT1-2-1 proteins of wild-type *C. sinensis* isolates. In addition, the analysis of DNA-binding domains presented in this study further revealed diverse stereostructures of the MATα_HMGbox domain of the MAT1-1-1 protein and the HMG-box_ROX1-like domain of the MAT1-2-1 protein. Apperantly, *O. sinensis* experiences self-sterility and uses a heterothallic or hybrid strategy, even parasexuality, to accomplish sexual reproduction during the lifecycle of the *C. sinensis* insect‒fungal complex [Kück 1993; Bennett & Johnson 2003; Sherwood & Bennett 2009; Seervai *et al*. 2013; Zhu & Li 2017; Nakamura *et al*. 2019; Samarasinghe *et al*. 2020; Steensels *et al*. 2021; Mishra *et al*. 2021; Kück *et al*. 2021; Li *et al*. 2023c, 2023e, 2024b, 2025].

Unlike homothallic reproduction, heterothallic or hybrid reproduction of *O. sinensis* requires mating partners. Among the 17 genotypes of *O. sinensis* [Kinjo & Zang 2001; Chen *et al*. 2004, 2011; Stensrud *et al*. 2005, 2007; Xiao *et al*. 2008; Zhu *et al*. 2010; Barseghyan *et al*. 2011; Li *et al*. 2013, 2022; Mao *et al*. 2013; Zhu & Li 2017; Du *et al*. 2020; Hėnault *et al*. 2020], GC-biased Genotype #1 *H. sinensis* has been proposed as the sole anamorph of *O. sinensis* [Wei *et al*. 2006]. Wei *et al*. [2016] reported successful industrial cultivation of *C. sinensis* insect‒fungal complexes; the cultivated complexes, however, exhibited a species contradiction between the anamorphic inoculants of 3 *H. sinensis* strains and the sole teleomorph of AT-biased Genotype #4 that was detected in the fruiting body of the cultivated *C. sinensis* insect‒fungal complexes. The sequences of AT-biased Genotypes #4−6 and #15−17, as well as the sequences of GC-biased Genotypes #2−3 and #7−14 of *O. sinensis*, do not reside in the genome assemblies ANOV00000000, JAAVMX000000000, LKHE00000000, LWBQ00000000, or NGJJ00000000 of the *H. sinensis* strains Co18, IOZ07, 1229, ZJB12195, and CC1406-20395, respectively [Stensrud *et al*. 2007; Xiao *et al*. 2008; Gao *et al*. 2012; Hu *et al*. 2013; Li *et al*. 2013, 2016a, 2020b, 2022, 2023d, 2023e; 2024a; Mao *et al*. 2013; Zhu & Li 2017; Jin *et al*. 2020; Liu *et al*. 2020; Shu *et al*. 2020]. Li *et al*. [2024a] further demonstrated that the sequences of Genotypes #2−17 of *O. sinensis* did not appear as repetitive copies in the genome of Genotype #1 *H. sinensis*, invalidating the hypothesis proposed by Li *et al*. [2013, 2017, 2020c] that “RIP mutation” induces or generates “ITS pseudogenes” and “rRNA pseudogenes” in the genome of *H. sinensis*. These data indicate that the 17 genotypes of *O. sinensis* are genome-independent and that they belong to different fungi. In addition, the data demonstrate that the sole anamorph hypothesis for *H. sinensis* proposed by Wei *et al*. [2006] fails to pass examination via Koch’s postulates.

Li *et al*. [2020a, 2023b] revised the inoculant information for the industrial cultivation project and reported that the cultivation project used cultures of *C. sinensis* ascospores rather than pure *H. sinensis* strains as inoculants, as previously reported by Wei *et al*. [2016]. However, Li *et al*. [2023d, 2023e] reported the detection of GC-biased Genotypes #1 and #14 and AT-biased Genotypes #5−6 and #16, as well as *P. hepiali,* in ascospores of natural *C. sinensis* insect‒fungal complexes through a culture‒independent approach using a PCR amplicon cloning technique with the selection and sequencing of ≥30 white colonies. In addition, Li *et al*. [2013] used a culture-dependent protocol and reported the detection of GC-biased Genotype #1 and AT-biased Genotype #5 in 8 cultures of mono-ascospores. Li *et al*. [2016b] demonstrated the nearly non-infectious feature of *H. sinensis* conidia and mycelia on 200 larvae of *H. armoricanus* in the inoculation experiments. Furthermore, Li *et al*. [2020b, 2022, 2023d, 2023e] and Gao *et al*. [2012] reported that AT-biased Genotype #4 of *O. sinensis* occurred in the stromata throughout the entire course of *C. sinensis* maturation and that the stromal fertile portion (SFP) contained numerous ascocarps of the *C. sinensis* insect‒fungal complex; the abundance of Genotype #4 was high in immature stromata, which are in asexual growth stages, and largely declined in mature stromata and SFP, which are in sexual production stages or in the transitional stages from asexual growth to sexual reproduction. Unfortunately, Genotype #4 of *O. sinensis* was absent from the *C. sinensis* ascospores. Thus, although Wei *et al*. [2016] reported the detection of the sole teleomorphic AT-biased Genotype #4 in the cultivated *C. sinensis* insect‒fungal complex and Li *et al*. [2020a, 2023b] reviewed the inoculant information, the fungal source of the sole teleomorphic, AT-biased Genotype #4 of *O. sinensis* remains scientifically uncertain in the artificial cultivation of the *C. sinensis* insect‒fungal complex.

In addition to the formal reports of unsuccessful attempts at artificial cultivation of the fruiting bodies of *C. sinensis* [Holliday *et al*. 2008; Stone 2010; Hu *et al*. 2013], Zhang *et al*. [2013] summarized the 40-year history of cultivation failures of insect‒fungal complexes using a “pure” mycological strategy in academic research-oriented settings and Qin *et al*. [2018] summarized the obstacles to the cultivation of *O. sinensis* fruiting bodies and ascospores. In contrast, Wei *et al*. [2016] reported success in such a cultivation effort in industrial product-oriented settings. This industrial success might be attributed to the application of a “mycologically impure” cultivation strategy on the basis of two facts: Cultures of mycologically impure ascospores of natural *C. sinensis* are used as inoculants [Li *et al*. 2020a, 2023b]; these are most likely combined with cocultures of *C. sinensis* stroma and/or a stromal fertile portion (SFP) containing numerous ascocarps that, most importantly, contain AT-biased Genotype #4 of *O. sinensis*, consistent with the discovery of the sole teleomorphic *O. sinensis* genotype [Wei *et al*. 2016]; Soil collected from the natural *C. sinensis* production areas on the Qinghai‒Tibet Plateau was added to the industrial cultivation system as described by Wei *et al*. [2016].

Unfortunately, purification and genomic sequencing of GC- and AT-biased Genotypes #2−17 of *O. sinensis* have not been reported to date. Thus, genomic and transcriptomic information regarding mutations in the *MAT1-1-1* and *MAT1-2-1* genes in the genome-independent *O. sinensis* fungi of Genotypes #2−17, especially the DNA-binding domains of the MAT1-1-1 and MAT1-2-1 proteins, is lacking. Table S3 shows that MAT1-1-1 proteins ALH25057, ALH25005, and ALH25006 and MAT1-2-1 proteins AIV43040, AFX66443, ACV60417, AFH35020, and ACV60418 were most likely produced by Genotype #3 of *O. sinensis*, if GC-biased Genotype #3 of *O. sinensis* does not coexist with other fungi in the wild-type *C. sinensis* isolates XZ12_16, XZ05_8, XZ-LZ07-H1, XZ06-124, and XZ-LZ07-H2.

Regardless of whether the MAT1-1-1 and MAT1-2-1 proteins form heterodimers or interact via complementary electrostatic surfaces and thereby enable cooperative DNA binding, the synergy of the mating proteins of *O. sinensis* constitutes the core mechanism that regulates mating type recognition, triggering downstream nuclear fusion signaling pathways and fruiting body development [Debuchy *et al*. 2006; Jones & Bennett 2011; Zheng & Wang 2013; Wilson *et al*. 2015; Sun *et al*. 2019]. The differential occurrence, differential transcription, and alternative splicing of mating-type and pheromone receptor genes in *H. sinensis* and *C. sinensis* isolates invalidates the self-fertilization hypothesis at the genomic and transcriptomic levels but suggests self-sterility of *O. sinensis* under heterothallic or hybrid reproduction [Hu *et al*. 2013; Bushley *et al*. 2013; Zhou *et al*. 2013; Wei *et al*. 2006; Li *et al*. 2019, 2023c, 2024b, 2025]. Furthermore, at the protein level, the discovery of many mutant MAT1-1-1 and MAT1-2-1 proteins reported by Li *et al*. [2025] that contain mutant MATα_HMGbox domains and HMG-box_ROX1-like domains, respectively, and have been detected in wild-type *C. sinensis* isolates, and the insect‒fungal complex demonstrated in this study, may indicate that mating proteins from heterogeneous fungal sources may accomplish coordinated heterothallic or hybrid reproduction of *O. sinensis* during the lifecycle of *C. sinensis* insect‒fungal complexes.

## Conclusions

Analysis of the DNA-binding domains of mating proteins revealed heteromorphic stereostructures of the MATα_HMGbox and HMG-box_ROX1-like domains of the MAT1-1-1 and MAT1-2-1 proteins, respectively. A few amino acid substitutions at various sites in the DNA-binding domains were demonstrated to be present in numerous mutant mating proteins derived from wild-type *C. sinensis* isolates. These substitutions correlated with changes in the hydrophobic properties and secondary and tertiary structures of the proteins under different AlphaFold 3D structural cords clustered into various Bayesian clustering clades. The diversity of heteromorphic MATα_HMGbox and HMG-box_ROX1-like domains of the mating proteins present in wild-type *C. sinensis* isolates may lead to stereostructure-related functional alterations in the mating process. The observed variation in the stereostructures of the DNA-binding domains of the mutant MAT1-1-1 and MAT1-2-1 proteins suggest that the mating proteins arise from diverse fungal sources and that they are produced by cooccurring *O. sinensis* genotypes or heterospecific fungi coexisting in wild-type *C. sinensis* isolates and natural *C. sinensis* insect-fungal complexes that have been identified in mycobiota, molecular, metagenomic, and metatranscriptomic studies, regardless of whether culture-dependent or culture-independent research strategies were used.

## Supporting information

C:\JSZ documents\Products\Cordyceps sinensis\Manuscripts\2025\2025 MAT domains isolates

## Acknowledgments

The authors are grateful to Prof. Mu Zang, Prof. Ru-Qin Dai, Prof. Ping Zhu, Prof. Wei Liu, Prof. Zong-Qi Liang and Prof. Yong-Jie Zhang for their consultation.

## Supplementary Materials

The following supporting information can be downloaded at www.xxxxxx.

**<u>Figure S1.</u>** Alignment of the MAT1-1-1 protein sequences with various animo acid residue substitutions derived from wild-type *C. sinensis* isolates and from the genome and metatranscriptome assemblies of *H. sinensis* strains or the *C. sinensis* insect−fungi complexes. The underlined segment in blue refer to the MATα_HMGbox domain (amino acids 51→225) of the reference MAT1-1-1 protein AGW27560, and the 9 external amino acid residues upstream and downstream of the domain are shown blue but not underlined. The amino acid substitution is shown in red, whereas the hyphens indicate identical amino acid residues and the spaces denote unmatched protein sequence gaps.

**<u>Figure S2.</u>** Alignment of the MAT1-2-1 protein sequences with various animo acid residue substitutions derived from wild-type *C. sinensis* isolates and from the genome and metatranscriptome assemblies of *H. sinensis* strains or the *C. sinensis* insect−fungi complexes. The underlined segment in blue refer to the HMG-box_ROX1-like domain (127→197 of the reference sequence AEH27625, and the 9 external amino acid residues upstream and downstream of the domain are shown blue but not underlined. The amino acid substitution is shown in red, whereas the hyphens indicate identical amino acid residues and the spaces denote unmatched protein sequence gaps.

**<u>Figure S3.</u>** Correlation of the changes in the primary and secondary structures of the MATalpha_HMGbox domains of MAT1-1-1 proteins: the reference protein AGW27560 derived from the *H. sinensis* strain CS68-2-1229 and the truncated protein encoded by the metatranscriptome assembly GAGW01008880 derived from the *C. sinensis* insect‒fungi complex. Panel A shows alignment of amino acid sequences of the MATalpha_HMGbox domain sequences of the MAT1-1-1 proteins; the hyphens indicate identical amino acid residues. The ExPASy ProtScale plots show below the changes in hydrophobicity (Panel B) and in the 2D structure (Panels C−F for α-helices, β-sheets, β-turns, and coils) with open rectangles in blue outlining the truncation region in the plots.

**<u>Table S1.</u>** GenBank accession numbers (in red in parentheses) for the full-length MAT1-1-1 proteins in the AlphaFold database under the corresponding AlphaFold UniProt codes **[Li *et al*. 2025]**.

**<u>Table S2.</u>** GenBank accession numbers (in red) for the full-length MAT1-2-1 proteins of 69 *H. sinensis* strains or *C. sinensis* isolates under the corresponding AlphaFold UniProt codes **[Li *et al*. 2025]**.

**<u>Table S3.</u>** Wild-type *C. sinensis* isolates, GenBank accession numbers for the ITS nucleic acid sequences and mating protein sequences, and percentage similarities *vs*. GC-biased *O. sinensis* Genotypes #1−3 and #7−9.

**<u>Table S4.</u>** Amino acid scales based on the general chemical characteristics of their side chains for ProtScale analysis (https://web.expasy.org/protscale/) to predict hydrophobicity and secondary structures (α-helices, β-sheets, β-turns, and coils) of proteins.

**<u>Table S5.</u>** Summary of the Bayesian clustering results in Figure S1 with the amino acid substitutions in the MATα_HMGbox domains of the 19 full-length MAT1-1-1 proteins of wild-type *C. sinensis* isolates under the AlphaFold UniProt codes and GenBank accession numbers.

**<u>Table S6.</u>** Summary of the Bayesian clustering results in Figure S1 with the amino acid substitutions in the MATα_HMGbox domains of the MAT1-1-1 proteins encoded by the genome assemblies of *H. sinensis* strains and the metatranscriptome assemblies of natural *C. sinensis* under the GenBank accession numbers.

**<u>Table S7</u>**. Summary of the Bayesian clustering results in Figure 4 with the amino acid substitutions in the HMG-box_ROX1-like domains of the 25 full-length MAT1-2-1 proteins of wild-type *C. sinensis* isolates under the AlphaFold UniProt codes and GenBank accession numbers.

**<u>Table S8.</u>** Summary of the Bayesian clustering results in Figure 4 with the amino acid substitutions and deletions in the HMG-box_ROX1-like domains of the MAT1-2-1 proteins encoded by the genome and transcriptome assemblies of *H. sinensis* strains and metatranscriptome assembly of natural *C. sinensis* insect‒fungi complexes under the GenBank accession numbers.

## Author Contributions

Conceptualization, XZL, YLL, and JSZ; methodology, JSZ; formal analysis, JSZ; investigation, XZL and JSZ; data curation, XZL and JSZ; writing–original draft preparation, JSZ; writing— review and editing, XZL, YLL, and JSZ; project administration, YLL; funding acquisition, YLL. All authors have read and agreed to the published version of the manuscript.

## Institutional Review Board Statement

Not applicable because this paper is an *in silico* reanalysis of public data.

## Informed Consent Statement

Not applicable because this paper is a public bioinformatic data reanalysis.

## Data Availability Statement

All sequence and 3D structure data are available in the GenBank and AlphaFold databases, except for one set of metatranscriptome sequences from natural *C. sinensis* that was uploaded to the repository database www.plantkingdomgdb.com/Ophiocordyceps_sinensis/data/cds/Ophiocordyceps_sinensis_CDS.fas (accessed from 18 May 2017 to 18 January 2018) by Xia *et al*. [2017], which is currently inaccessible, but a previously downloaded cDNA file (accessed on 18 January 2018) was used for the mating protein analysis.

