## Supplementary material for "Altered stereostructures of the DNA-binding domains in the mutant mating proteins of *Ophiocordyceps sinensis* and the *Cordyceps sinensis* insect‒fungal complex": C:\JSZ documents\Products\Cordyceps sinensis\Manuscripts\2025\2025 MAT domains isolates

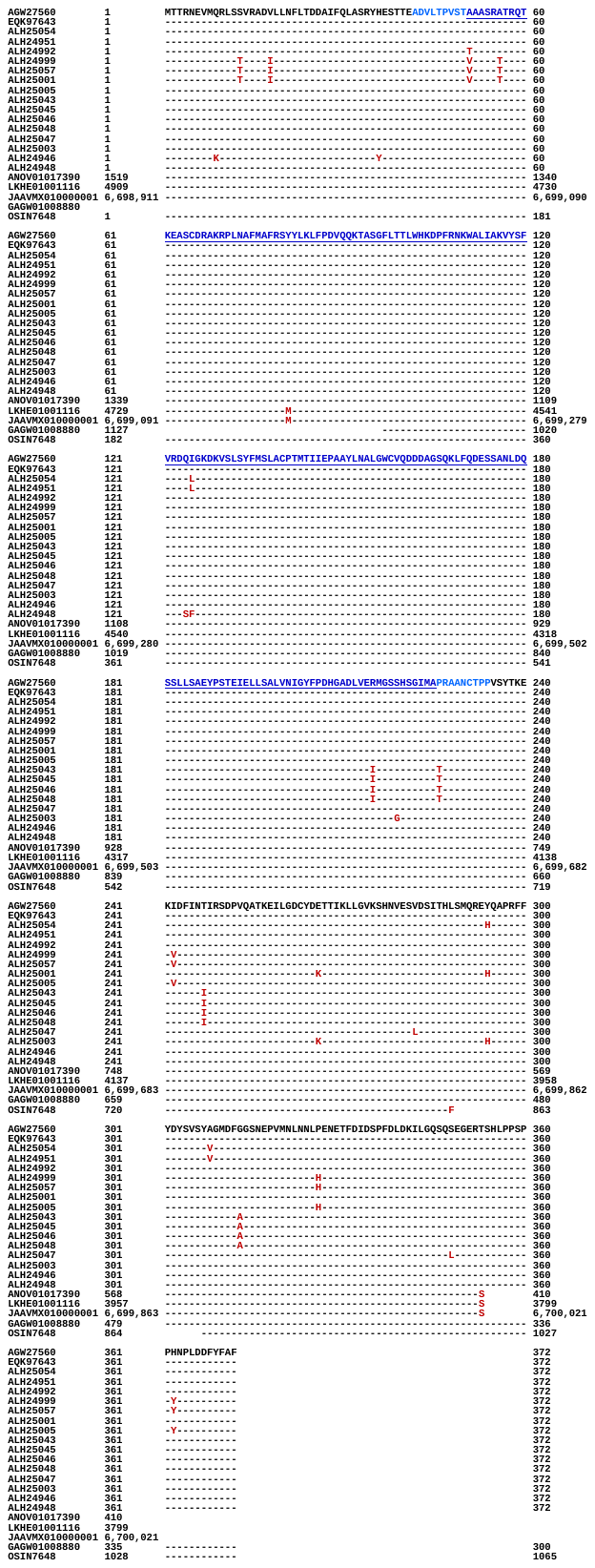

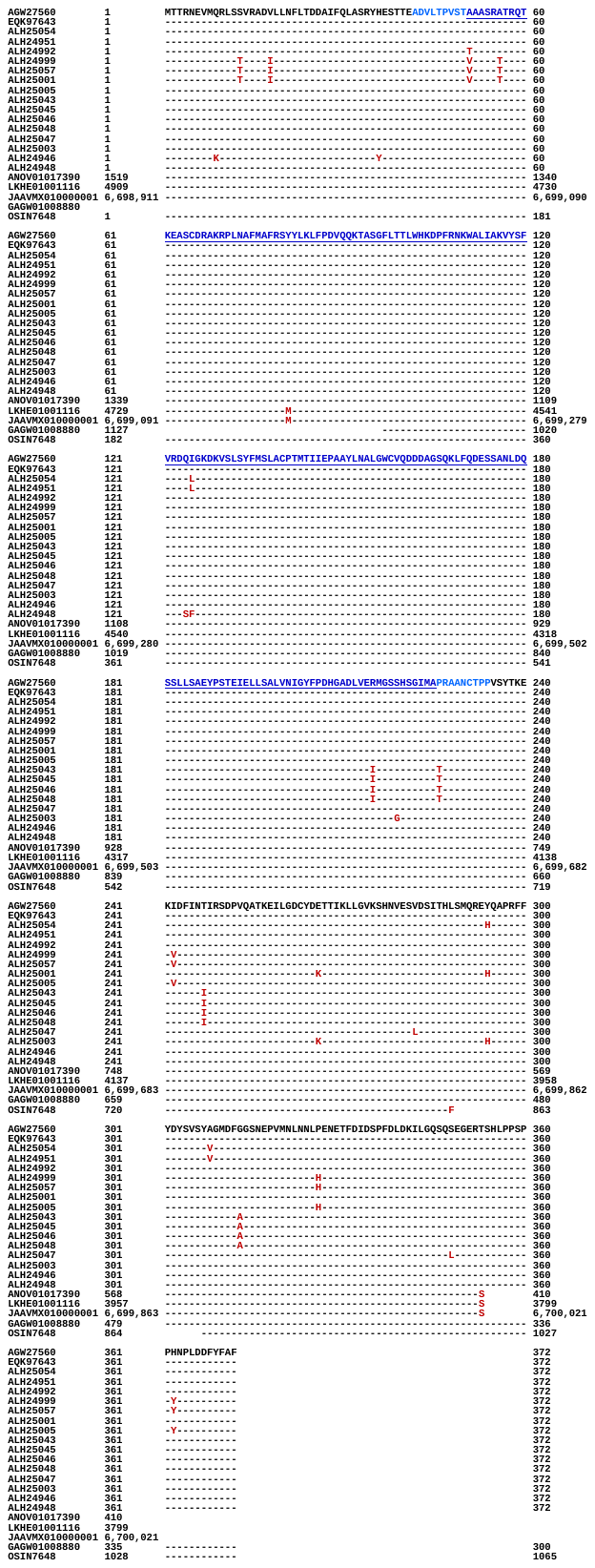

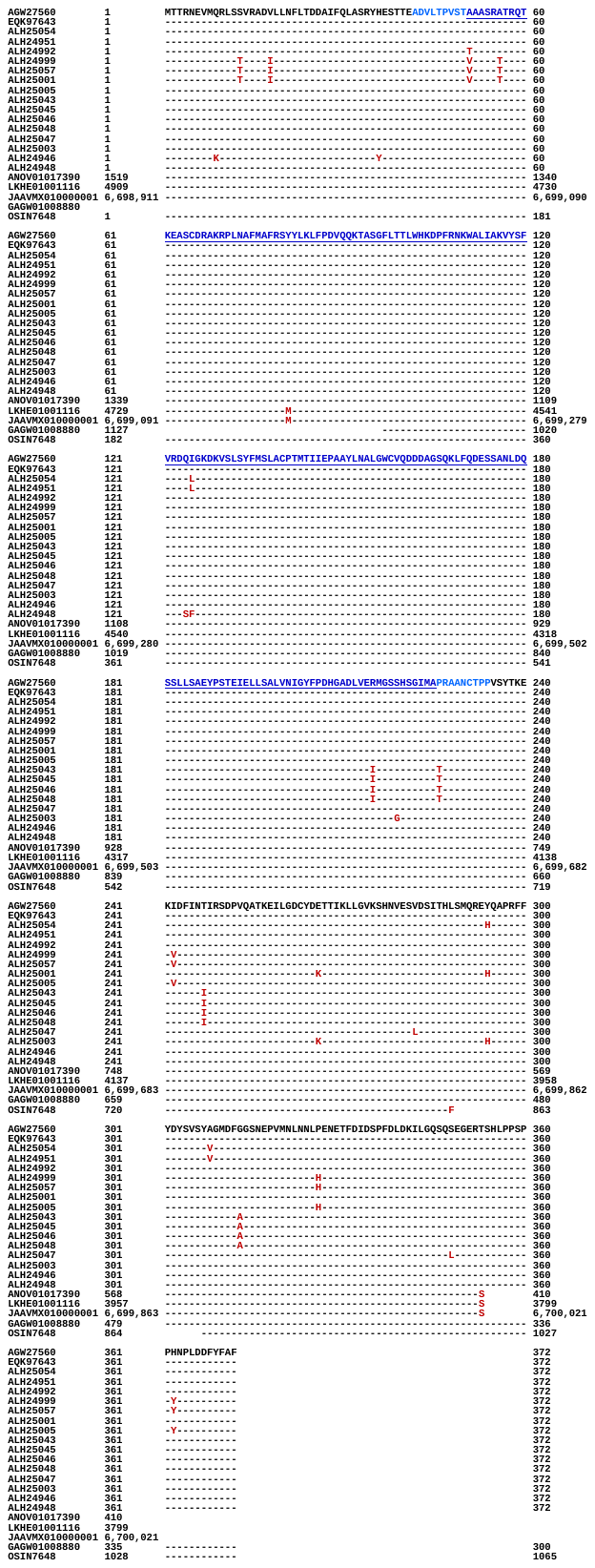

### Figure S1. Alignment of the reference MAT1-1-1 protein AGW27560 sequence derived from the *Hirsutella sinensis* strain CS68-2-1229 [Li *et al*. 2013] and the mutant protein sequences derived from the wild-type *Cordyceps sinensis* isolates with various animo acid residue substitutions derived from wild-type *C. sinensis* isolates [Zhang & Zhang 2015] and derived from the genome and metatranscriptome assemblies of *H. sinensis* strains or the *C. sinensis* insect−fungi complexes [Hu *et al*. 2013; Xiang *et al*. 2014; Li *et al*. 2016; Xia *et al*. 2017; Shu *et al*. 2020]. The underlined segment in blue refer to the MATα_HMGbox domain (amino acids 51→225) of the reference MAT1-1-1 protein AGW27560, and the 9 external amino acid residues upstream and downstream of the domain are shown blue but not underlined. The amino acid substitution is shown in red, whereas the hyphens indicate identical amino acid residues and the spaces denote unmatched protein sequence gaps.

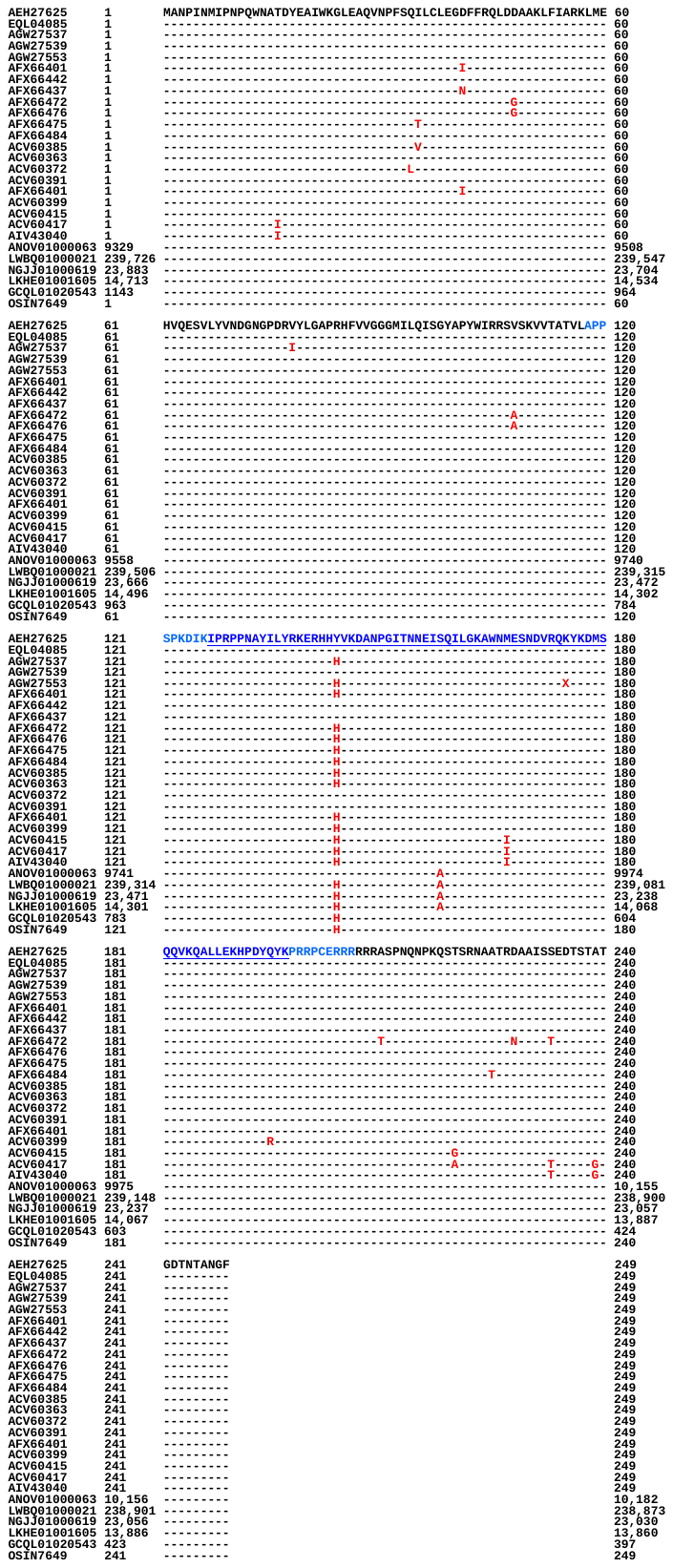

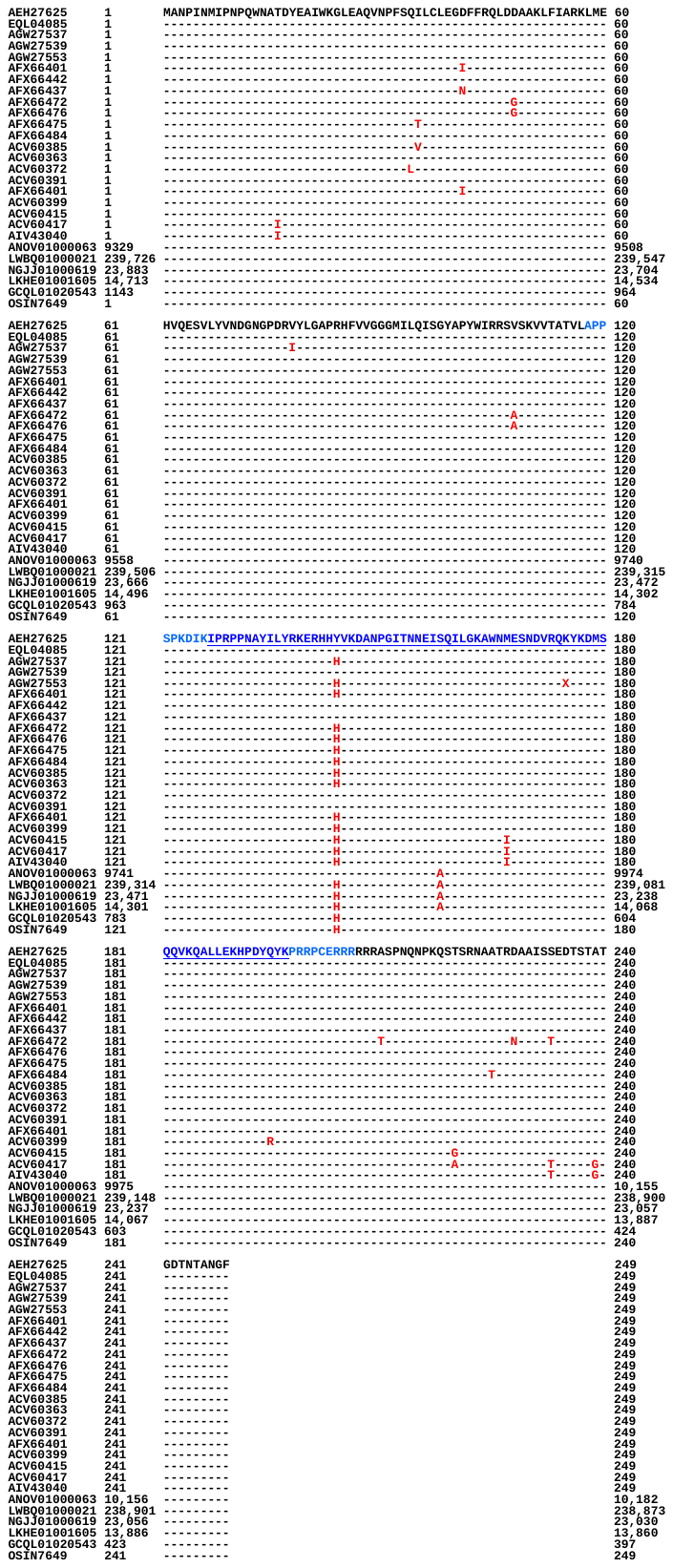

### Figure S2. Alignment of the reference MAT1-2-1 protein sequence AEH27625 derived from the *Hirsutella sinensis* strain CS2 [Zhang *et al*. 2009], and the mutant MAT1-2-1 proteins derived from the wild-type *Cordyceps sinensis* isolates with various animo acid residue substitutions [Zhang & Zhang 2015] and derived from the genome and metatranscriptome assemblies of *H. sinensis* strains or the *C. sinensis* insect−fungi complexes [Hu *et al*. 2013; Liu *et al*. 2015, 2020; Li *et al*. 2016; Xia *et al*. 2017; Jin *et al*. 2020]. The underlined segment in blue refer to the HMG-box_ROX1-like domain (127→197 of the reference sequence AEH27625, and the 9 external amino acid residues upstream and downstream of the domain are shown blue but not underlined. The amino acid substitution is shown in red, whereas the hyphens indicate identical amino acid residues and the spaces denote unmatched protein sequence gaps.

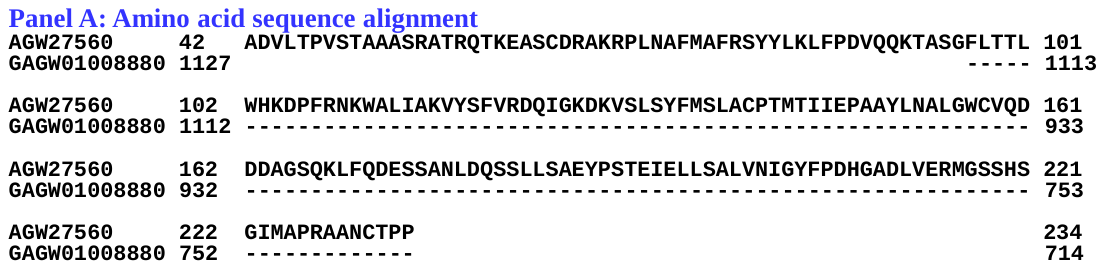

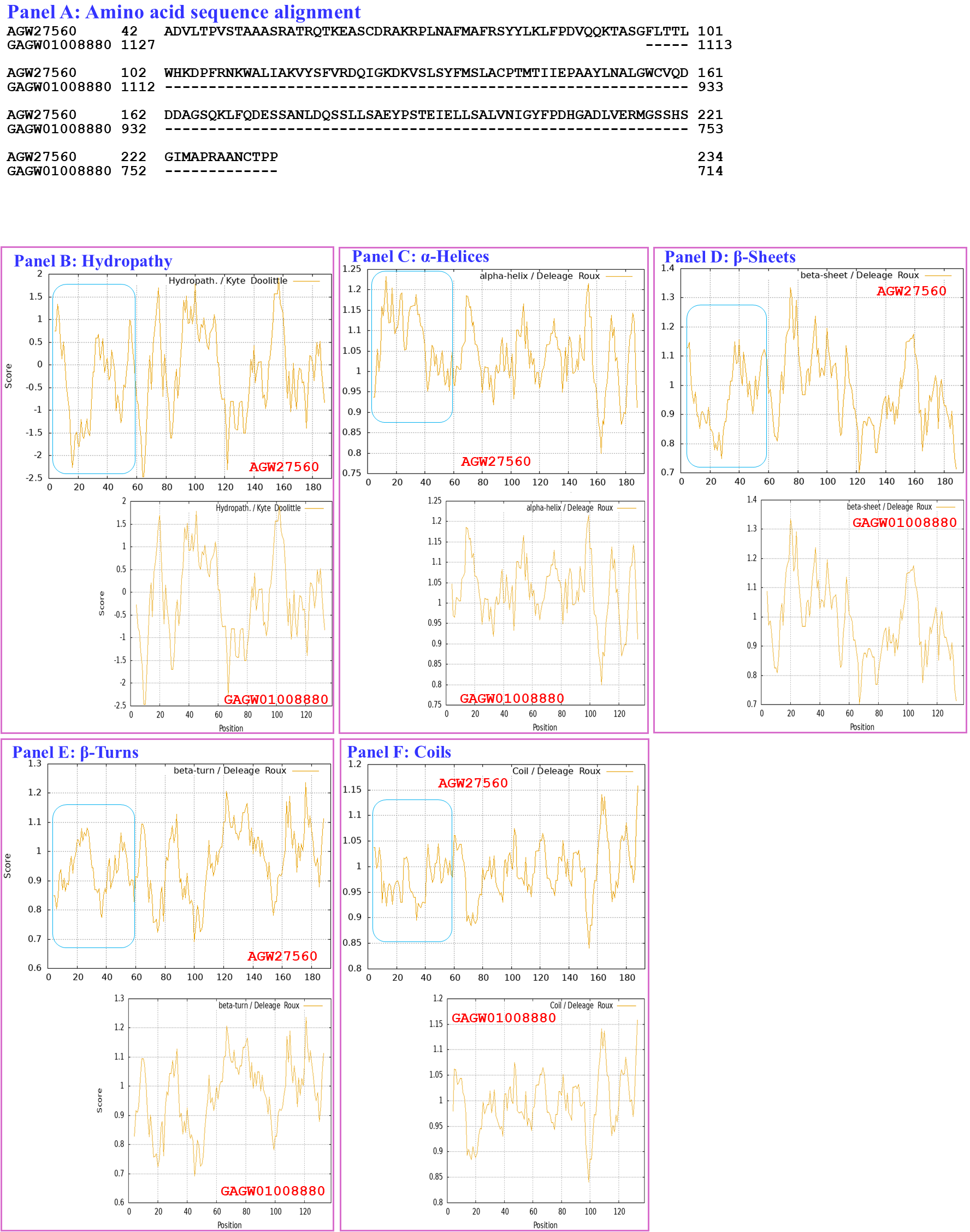

### Figure S3. Correlation of the changes in the primary and secondary structures of the MATalpha_HMGbox domains of MAT1-1-1 proteins: the reference protein AGW27560 is derived from the *H. sinensis* strain CS68-2-1229 [Li *et al*. 2013], and the truncated MAT1-1-1 protein is encoded by the metatranscriptome assembly GAGW01008880 derived from the *C. sinensis* insect-fungi complex [Xiang *et al*. 2014]. Panel A shows an alignment of the amino acid sequences in the MATα_HMGbox domains of the MAT1-1-1 proteins; amino acid substitutions are shown in red, whereas the hyphens indicate identical amino acid residues and the spaces denote unmatched protein sequence gaps. The ExPASy ProtScale plots show the changes in hydrophobicity (Panel B) and 2D structure (Panels C−F show the α-helices, β-sheets, β-turns, and coils, respectively); the open rectangles in blue outline the truncation region in the plots.

### Table S1. GenBank accession numbers (in red in parentheses) for the full-length MAT1-1-1 proteins in the AlphaFold database under the corresponding AlphaFold UniProt codes [Li *et al*. 2025].

| **AlphaFold UniProt code** | **Bayesian cluster-branch** | **Strain/isolate number (GenBank accession number for MAT1-2-1 protein)** |
| --- | --- | --- |
| U3N942 | **A1** | **GS09_111** (**ALH24945**), **CS68-2-1229** (**AGW27560**),  GS09_131 (**ALH24947**), ID10_1 (**ALH24954**),  IOZ07 (**KAF4512729**), NP10_1 (**ALH24955**),  NP10_2 (**ALH24956**), QH07_188 (**ALH24957**),  QH07_197 (**ALH24958**), QH09_122 (**ALH24959**),  QH09_131 (**ALH24960**), QH09_151 (**ALH24961**),  QH09_20L (**ALH24965**), QH09_33L (**ALH24967**),  QH09_37 (**ALH24968**), QH09_46 (**ALH24969**),  QH09_56 (**ALH24970**), QH09_66 (**ALH24971**),  QH09_78 (**ALH24972**), QH09_93 (**ALH24973**),  QH10_1 (**ALH24974**), QH10_4 (**ALH24975**),  QH10_7 (**ALH24976**), SC09_107 (**ALH24978**),  SC09_117 (**ALH24979**), SC09_128 (**ALH24980**),  SC09_147 (**ALH24981**), SC09_157 (**ALH24982**),  SC09_167 (**ALH24983**), SC09_180 (**ALH24984**),  SC09_190 (**ALH24985**), SC09_200 (**ALH24986**),  SC09_21 (**ALH24987**), SC09_36 (**ALH24988**),  SC09_37 (**ALH24989**), SC09_47 (**ALH24990**),  SC09_57 (**ALH24991**), SC09_77 (**ALH24993**),  SC10_18 (**ALH24996**), SC10_21 (**ALH24997**),  SC10_4 (**ALH24998**), XZ05_12 (**ALH25000**),  XZ05_3 (**ALH25002**), XZ05_7 (**ALH25004**),  XZ06_124 (**ALH25006**), XZ06_152 (**ALH25007**),  XZ07_108 (**ALH25009**), XZ07_133 (**ALH25010**),  XZ07_154 (**ALH25011**), XZ07_166 (**ALH25012**),  XZ07_176 (**ALH25013**), XZ07_180 (**ALH25014**),  XZ08_10 (**ALH25015**), XZ08_24 (**ALH25016**),  XZ08_26 (**ALH25017**), XZ08_4 (**ALH25018**),  XZ08_56 (**ALH25019**), XZ08_59 (**ALH25020**),  XZ08_A1 (**ALH25021**), XZ08_B1 (**ALH25022**),  XZ09_106 (**ALH25024**), XZ09_113 (**ALH25025**),  XZ09_118 (**ALH25026**), XZ09_15 (**ALH25027**),  XZ09_32 (**ALH25028**), XZ09_4 (**ALH25029**),  XZ09_46 (**ALH25030**), XZ09_48 (**ALH25031**),  XZ09_59 (**ALH25032**), XZ09_71 (**ALH25033**),  XZ09_80 (**ALH25055**), XZ10_15 (**ALH25035**),  XZ10_17 (**ALH25036**), XZ10_23 (**ALH25037**),  XZ10_7 (**ALH25038**), XZ12_1 (**ALH25056**),  XZ12_33 (**ALH25058**), XZ12_43 (**ALH25059**),  YN07_6 (**ALH25039**), YN07_8 (**ALH25040**),  YN09_101 (**ALH25041**), YN09_140 (**ALH25042**),  YN09_3 (**ALH25044**), YN09_72 (**ALH25049**),  YN09_81 (**ALH25050**), YN09_85 (**ALH25051**),  YN09_89 (**ALH25052**), YN09_96 (**ALH25053**) |
| A0A0N9QMM1 | **A1** | GS09_121 (**ALH24946**), GS09_201 (**ALH24949**),  GS09_225 (**ALH24950**), SC09_1 (**ALH24977**) |
| T5A511 | **A1** | **Co18** (**EQK97643**) (**KE657544** **410←1519**)  (**ANOV01017390 410←1519**) |
| A0A0N9R5B3 | **A2** | SC09_65 (**ALH24992**) |
| A0A0N7G849 | **A2** | SC09_97 (**ALH24995**) |
| A0A0N9QUF3 | **A3** | GS09_143 (**ALH24948**) |
| A0A0N9R4V2 | **A3** | YN09_61 (**ALH25047**) |
| A0A0N9QMS9 | **B** | YN09_22 (**ALH25043**), YN09_51 (**ALH25045**),  YN09_6 (**ALH25046**), YN09_64 (**ALH25048**) |
| A0A0N7G845 | **C** | GS09_229 (**ALH24951**), GS09_281 (**ALH24952**),  GS09_311 (**ALH25054**), GS10_1 (**ALH24953**),  QH09_164 (**ALH24962**), QH09_173 (**ALH24963**),  QH09_201 (**ALH24964**), QH09_210 (**ALH24966**),  SC09_87 (**ALH24994**) |
| A0A0N9QUK2 | **D1** | XZ05_8 (**ALH25005**) |
| A0A0N9QMT4 | **D2** | XZ07_H2 (**ALH24999**), XZ12_16 (**ALH25057**) |
| A0A0N9QMR3 | **E1** | XZ06_260 (**ALH25008**), XZ09_100 (**ALH25023**) |
| A0A0N9QMS4 | **E2** | XZ09_95 (**ALH25034**) |
| A0A0N7G850 | **E3** | XZ05_6 (**ALH25003**) |
| A0A0N9R4Q4 | **E4** | XZ05_2 (**ALH25001**) |

Note: *, Branch 1 in red, Branch 2 in pink, Branch 3 in purple, and Branch 4 in brown under the cluster codes (English letters) in the paratheses were determined *via* the Bayesian analysis shown in Figure 1 of [Li *et al*. 2025]. The “←” arrows indicate sequences in the antisense strands of the genome of the *H. sinensis* strain Co18.

### Table S2. GenBank accession numbers (in red) for the full-length MAT1-2-1 proteins of 69 *H. sinensis* strains or *C. sinensis* isolates under the corresponding AlphaFold UniProt codes [Li *et al*. 2025].

| **AlphaFold UniProt code** | **Bayesian cluster-branch** | **Strain/isolate number (GenBank accession number for MAT1-1-1 protein)** |
| --- | --- | --- |
| D7F2E9 | **I-1** | **CS2** (**AEH27625**) (**ACV60400**), SC-2 (**ACV60395**),  SC-4 (**ACV60396**), SC-5 (**ACV60398**),  SC-7 (**ACV60397**), XZ-LZ06-1 (**ACV60369**),  XZ-LZ06-108 (**ACV60373**), XZ-LZ06-21 (**ACV60371**),  XZ-LZ06-7 (**ACV60370**), XZ-LZ07-108 (**ACV60379**),  XZ-LZ07-30 (**ACV60377**), XZ-ML-191 (**ACV60376**),  YN-1 (**ACV60390**), YN-5 (**ACV60392**),  YN-6 (**ACV60393**), YN-8 (**ACV60394**),  SC09_47 (**AFX66423**), SC09_57 (**AFX66424**),  SC09_77 (**AFX66426**), SC09_97 (**AFX66428**),  XZ05_12 (**AFX66444**), XZ05_7 (**AFX66442**),  XZ06_152 (**AFX66445**), XZ07_11 (**AFX66447**),  XZ07_46 (**AFX66448**), XZ09_106 (**AFX66464**),  XZ09_113 (**AFX66465**), XZ09_15 (**AFX66455**),  YN09_101 (**AFX66482**), YN09_72 (**AFX66477**),  YN09_81 (**AFX66478**), YN09_85 (**AFX66479**),  YN09_89 (**AFX66480**), SC09-37 (**AFH35019**),  CS26-277 (**AGW27541**), CS36-1294 (**AGW27538**),  CS37-295 (**AGW27539**) |
| T5AF56 | **I-1** | **Co18** (**EQL04085**) (**ANOV01000063 9329→10182**) |
| V9LW10 | **I-2** | SC09_200 (**AFX66437**) |
| D7F2H1 | **I-2** | YN-4 (**ACV60391**) |
| D7F2F2 | **I-2** | XZ-LZ06-61 (**ACV60372**) |
| A0A0A0RCF5 | **II-1** | XZ12_16 (**AIV43040**) |
| D7F2J7 | **II-2** | XZ-LZ07-H1 (**ACV60417**), XZ-LZ07-H2 (**ACV60418**),  XZ06-124 (**AFH35020**), XZ05_8 (**AFX66443**) |
| D7F2F5 | **III** | XZ-LZ05-6 (**ACV60415**), XZ-SN-44 (**ACV60375**),  XZ05_2 (**AFX66441**), XZ06_260 (**AFX66446**),  XZ09_100 (**AFX66463**), XZ09_80 (**AFX66461**),  XZ09_95 (**AFX66462**) |
| V9LWC9 | **IV-1** | YN09_64 (**AFX66476**) |
| V9LVS8 | **IV-2** | YN09_6 (**AFX66472**), YN09_22 (**AFX66473**),  YN09_51 (**AFX66474**) |
| D7F2E3 | **V-1** | XZ-NQ-154 (**ACV60363**), XZ-NQ-155 (**ACV60364**),  GS09_111 (**AFX66388**), QH09-93 (**AFH35018**),  CS560-961 (**AGW27542**) |
| D7F2G5 | **V-2** | QH-YS-199 (**ACV60385**) |
| D7F2H9 | **V-2** | SC-3 (**ACV60399**) |
| V9LW71 | **V-2** | QH09_11 (**AFX66401**) |
| V9LVU8 | **V-2** | YN09_61 (**AFX66475**) |
| V9LWG5 | **V-2** | ID10_1 (**AFX66484**) |
| U3N6V5 | **V-2** | CS6-251 (**AGW27537**) |
| ‡ | **V-1** | NP10_1 (AFX66485), NP10_2 (AFX66486),  YN09_3 (AFX66471), YN09_96 (AFX66481),  YN09_140 (AFX66483) |

Note: **, Branch 1 in red and Branch 2 in pink under the cluster codes (Roman numerals) in the paratheses were determined *via* the Bayesian analysis shown in Figure 2 of [Li *et al*. 2025]. ‡, The 5 MAT1-2-1 protein sequences in green are included in the GenBank database but not in the AlphaFold database. The “→” arrow indicates the sequence in the sense strand of the genome of the *H. sinensis* strain Co18.The “→” arrow indicates the sequence in the sense strand of the genome of the *H. sinensis* strain Co18.

### Table S3. Wild-type *C. sinensis* isolates, GenBank accession numbers for the ITS nucleic acid sequences and mating protein sequences, and percentage similarities *vs*. GC-biased *O. sinensis* Genotypes #1−3 and #7−9.

| **Wild-type *C. sinensis* isolate** | **GenBank accession #** | | |  | **% similarity *vs*. GC-biased *O. sinensis* genotype** | | | | | | |
| --- | --- | --- | --- | --- | --- | --- | --- | --- | --- | --- | --- |
|  | **ITS1-5.8S-ITS2** | **Mating protein** | |  |  |  |  |  |  |  |  |
|  |  | **MAT1-1-1** | **MAT1-2-1** |  | **#1** | **#2** | **#3** | **#7** | **#8** | **#9** | **#10** |
| GS09_143 | JQ325056 | ALH24948 | AFX66391 |  | **100%** | 95.0% | 88.8% | 89.5% | 95.2% | 83.1% | 83.1% |
| SC09_65 | JQ325090 | ALH24992 | AFX66425 |  | **100%** | 95.0% | 88.8% | 89.5% | 95.2% | 83.1% | 83.1% |
| GS09_111 | JQ325053 | ALH24945 | AFX66388 |  | **100%** | **97.2%** | 95.0% | 95.0% | 89.5% | 95.2% | 83.1% |
| QH09_11 | JQ325066 |  | AFX66401 |  | **100%** | **97.2%** | 95.0% | 95.0% | 89.5% | 95.2% | 83.1% |
| QH09-93 | JQ286746 | ALH24973 | AFH35018 |  | **100%** | **97.2%** | 95.0% | 95.0% | 89.5% | 95.2% | 83.1% |
| QH-YS-199 | FJ654226 |  | ACV60385 |  | **99.8%** | 96.8% | 94.8% | 94.7% | 89.3% | 95.0% | 82.9% |
| XZ-NQ-154 | FJ654206 |  | ACV60363 |  | **99.8%** | 96.8% | 94.8% | 94.7% | 89.3% | 95.0% | 82.9% |
| XZ-NQ-155 | FJ654207 |  | ACV60364 |  | **99.8%** | 96.8% | 94.8% | 94.7% | 89.3% | 95.0% | 82.9% |
| GS09_229 | JQ325059 | ALH24951 | AFX66394 |  | **99.1%** | 94.4% | 88.6% | 88.9% | 94.6% | 82.8% | 82.8% |
| GS09_281 | JQ325061 | ALH24952 | AFX66396 |  | **99.1%** | 94.0% | 88.6% | 88.9% | 94.6% | 82.8% | 82.8% |
| GS10_1 | JQ325064 | ALH24953 | AFX66399 |  | **99.1%** | 94.0% | 88.6% | 88.9% | 94.6% | 82.8% | 82.8% |
| QH09_164 | JQ325077 | ALH24962 | AFX66412 |  | **99.1%** | 94.0% | 88.6% | 88.9% | 94.6% | 82.8% | 82.8% |
| QH09_173 | JQ325078 | ALH24963 | AFX66413 |  | **99.1%** | 94.0% | 88.6% | 88.9% | 94.6% | 82.8% | 82.8% |
| QH09_201 | JQ325080 | ALH24964 | AFX66415 |  | **99.1%** | 94.0% | 88.6% | 88.9% | 94.6% | 82.8% | 82.8% |
| QH09_210 | JQ325081 | ALH24966 | AFX66416 |  | **99.1%** | 94.0% | 88.6% | 88.9% | 94.6% | 82.8% | 82.8% |
| SC09_87 | JQ325092 | ALH24994 |  |  | **99.1%** | 94.0% | 88.6% | 88.9% | 94.6% | 82.8% | 82.8% |
| GS09_311 | JQ325062 | ALH25054 | AFX66397 |  | **98.9%** | 94.2% | 88.4% | 88.7% | 94.4% | 82.6% | 82.8% |
| XZ05_2 | JQ325106 | ALH25001 | AFX66441 |  | **98.9%** | 95.0% | 88.6% | 90.1% | 95.3% | 83.5% | 83.5% |
| XZ09_80 | JQ325126 | ALH25055 | AFX66461 |  | **98.9%** | 96.8% | 95.0% | 93.7% | 90.0% | 95.3% | 83.5% |
| XZ-LZ05-6 | FJ654259 |  | ACV60415 |  | **98.9%** | 96.8% | 95.0% | 93.7% | 90.0% | 95.3% | 83.5% |
| YN09_61 | JQ325140 | ALH25047 | AFX66475 |  | **98.9%** | **97.2%** | 95.0% | 94.1% | 90.2% | 96.1% | 84.0% |
| ID10_1 | JQ325149 | ALH24954 | AFX66484 |  | **98.9%** | **97.2%** | 95.0% | 94.1% | 90.2% | 96.1% | 84.0% |
| SC-3 | FJ654238 |  | ACV60399 |  | **98.7%** | 96.8% | 95.0% | 93.7% | 90.4% | 95.5% | 83.7% |
| XZ06_260 | JQ325111 | ALH25008 | AFX66446 |  | **98.7%** | 96.8% | 94.8% | 93.5% | 89.9% | 95.2% | 83.5% |
| XZ-SN-44 | FJ654218 |  | ACV60375 |  | **98.7%** | 96.8% | 94.8% | 93.5% | 89.9% | 95.2% | 83.5% |
| YN09_51 | JQ325139 | ALH25045 | AFX66474 |  | **97.6%** | 95.0% | 87.1% | 89.1% | 94.0% | 82.0% | 82.0% |
| YN09_6 | JQ325137 | ALH25046 | AFX66472 |  | **97.2%** | 95.1% | 86.4% | 88.4% | 93.5% | 81.2% | 81.2% |
| YN09_22 | JQ325138 | ALH25043 | AFX66473 |  | **97.0%** | 94.9% | 86.2% | 88.2% | 93.3% | 81.0% | 81.0% |
| YN09_64 | JQ325141 | ALH25048 | AFX66476 |  | **97.0%** | 94.5% | 86.6% | 88.6% | 93.5% | 81.6% | 81.6% |
| XZ12_16 | KM197540 | ALH25057 | AIV43040 |  | 95.7% | 96.1**%** | **99.6%** | 87.7% | 92.2% | 81.1% | 81.1% |
| XZ05_8 | JQ325108 | ALH25005 | AFX66443 |  | 96.1% | 95.8% | **98.6%** | 91.5% | 87.9% | 92.5% | 81.1% |
| XZ-LZ07-H1 | FJ654148 |  | ACV60417 |  | 96.1% | 95.8% | **98.6%** | 91.5% | 87.9% | 92.5% | 81.1% |
| XZ06-124 | JQ286748 | ALH25006 | AFH35020 |  | 96.3% | 95.8% | **98.4%** | 91.8% | 88.1% | 92.7% | 81.3% |
| XZ-LZ07-H2 | FJ654149 |  | ACV60418 |  | 96.3% | 95.8% | **98.4%** | 91.8% | 88.1% | 92.7% | 81.3% |

Note: The accession numbers in green refer to the MAT1-1-1 and MAT1-2-1 protein sequences recorded in the GenBank database but not in the AlphaFold database. Those in red recorded in both the GenBank and AlphaFold databases. Percent numbers in blue indicate high homology (≥97%) to the reference sequences of GC-biased Genotypes #1 (AB067721), #2 (MG770309), #3 (HM595984), #7 (AJ488254), #8 (GU246286), #9 (GU246288), and #10 (GU246287) [Zhu & Li 2017; Li *et al*. 2022]

.

### Table S4. Amino acid scales based on the general chemical characteristics of their side chains for ProtScale analysis (https://web.[expasy.org/protscale](https://web.expasy.org/protscale/)/) to predict hydrophobicity and secondary structures (α-helices, β-sheets, β-turns, and coils) of proteins.

|  |  | **Chemical-physical property** | **hydropathy index *** | **α-Helix** | | **β-Sheet** | | **β-Turn** | | **Coil** | |
| --- | --- | --- | --- | --- | --- | --- | --- | --- | --- | --- | --- |
| Phenylalanine | Phe, F | Aromatic | 2.800 | | 1.195 | | 1.393 | | 0.624 | | 0.797 |
| Tryptophan | Trp, W | Aromatic | -0.900 | | 1.090 | | 1.306 | | 0.546 | | 0.941 |
| Tyrosine | Tyr, Y | Aromatic | -1.300 | | 0.787 | | 1.266 | | 0.795 | | 1.109 |
| Isoleucine | Ile, I | Aliphatic | 4.500 | | 1.003 | | 1.799 | | 0.240 | | 0.886 |
| Valine | Val, V | Aliphatic | 4.200 | | 0.990 | | 1.965 | | 0.387 | | 0.772 |
| Leucine | Leu, L | Aliphatic | 3.800 | | 1.236 | | 1.261 | | 0.670 | | 0.810 |
| Alanine | Ala, A | Aliphatic | 1.800 | | 1.489 | | 0.709 | | 0.788 | | 0.824 |
| Cysteine | Cys, C | with polar neutral side chains | 2.500 | | 0.966 | | 1.191 | | 0.965 | | 0.953 |
| Methionine | Met, M | with polar neutral side chains | 1.900 | | 1.363 | | 1.210 | | 0.436 | | 0.810 |
| Serine | Ser, S | with polar neutral side chains | -0.800 | | 0.739 | | 0.928 | | 1.316 | | 1.130 |
| Threonine | Thr, T | with polar neutral side chains | -0.700 | | 0.785 | | 1.221 | | 0.739 | | 1.148 |
| Asparagine | Asn, N | with polar neutral side chains | -3.500 | | 0.772 | | 0.604 | | 1.572 | | 1.167 |
| Glutamine | Gln, Q | with polar neutral side chains | -3.500 | | 1.164 | | 0.840 | | 0.997 | | 0.947 |
| Histidine | His, H | Basic | -3.200 | | 1.003 | | 0.863 | | 0.970 | | 1.068 |
| Lysine | Lys, K | Basic | -3.900 | | 1.172 | | 0.721 | | 1.302 | | 0.897 |
| Arginine | Arg, R | Basic | -4.500 | | 1.224 | | 0.920 | | 0.912 | | 0.893 |
| Aspartic acid | Asp, D | Acidic | -3.500 | | 0.924 | | 0.541 | | 1.197 | | 1.197 |
| Glutamic acid | Glu, E | Acidic | -3.500 | | 1.504 | | 0.567 | | 1.149 | | 0.761 |
| Glycine | Gly, G | Unique amino acid | -0.400 | | 0.510 | | 0.657 | | 1.860 | | 1.251 |
| Proline | Pro, P | Unique amino acid | -1.600 | | 0.492 | | 0.354 | | 1.415 | | 1.540 |

Note: An **amino acid scale** is defined at https://web.[expasy.org/protscale](https://web.expasy.org/protscale/)/ by a numerical value assigned to each type of amino acid. The most frequently used scales are the hydrophobicity or hydrophilicity scales and the secondary structure conformational parameter scales, but many other scales exist, which are based on the different chemical and physical properties of the amino acids. The ExPASy ProtScale program provides 57 predefined scales based on the literature [Deleage, & Roux 1987]. *, Hydropathy index [Kyte & Doolittle 1982]; the larger the value, the stronger the hydrophobicity; negative values indicate hydrophilicity.

### Table S5. Summary of the Bayesian clustering results in Figure S1 with the amino acid substitutions in the MATα_HMGbox domains of the 19 full-length MAT1-1-1 proteins of wild-type *C. sinensis* isolates under the AlphaFold UniProt codes and GenBank accession numbers.

| ***C. sinensis* isolate** | **GenBank accession #** | **AlphaFold UniProt code** | **The Bayesian cluster based on** | | **Amino acid substitution in the MATα_HMGbox domain (*vs*.** **AGW27560)** |
| --- | --- | --- | --- | --- | --- |
|  |  |  | **full length sequence*** | **the sequence of MATα_HMGbox domain** |  |
| SC09_65 | ALH24992 | A0A0N9R5B3 | A2 | a2 | A-to-D |
| GS09_143 | ALH24948 | A0A0N9QUF3 | A3 | e2 | QI-to-SF |
| YN09_22 YN09_51 YN09_6 YN09_64 | ALH25043 ALH25045 ALH25046 ALH25048 | A0A0N9QMS9 | B | b | R-to-I |
| GS09_311 GS09_229 GS09_281 GS10_1 QH09_164 QH09_173 QH09_201 QH09_210 SC09_87 | ALH25054 ALH24951 ALH24952 ALH24953 ALH24962 ALH24963 ALH24964 ALH24966 ALH24994 | A0A0N7G845 | C | e1 | I-to-L |
| XZ07_H2 XZ12_16 | ALH24999 ALH25057 | A0A0N9QMT4 | D2 | d | A-to-V and A-to-T |
| XZ05_2 | ALH25001 | A0A0N9R4Q4 | E4 | d | A-to-V and A-to-T |
| XZ05_6 | ALH25003 | A0A0N7G850 | E3 | a2 | S-to-G |

Note: *, The Bayesian clustering results were reported previously by Li *et al*. [2025].

### Table S6. Summary of the Bayesian clustering results in Figure S1 with the amino acid substitutions in the MATα_HMGbox domains of the MAT1-1-1 proteins encoded by the genome assemblies of *H. sinensis* strains and the metatranscriptome assemblies of natural *C. sinensis* under the GenBank accession numbers.

| ***H. sinensis* strain** | **GenBank accession #** | **The MATα_HMGbox domain** | | |
| --- | --- | --- | --- | --- |
|  |  | **Nucleotide sequence arrange (deleted the intron sequence)** | **Bayesian cluster** | **Amino acid substitution or deletion (*vs*. AGW27560)** |
| IOZ07 | JAAVMX010000001 | 6,699,061→6,699,153 & 6,699,203→6,699.637 | c | Y-to-M |
| 1229 | LKHE01001116 | 4183←4620 & 4667←4759 | c | Y-to-M |
| Co18 | ANOV01017390 | 794←1129 & 1280←1396 | a1 | (100% identical) |
| † | OSIN7648 | 151→675 | a1 | (100% identical) |
| † | GAGW01008880 | 714←1127 | a1 | 46 aa deletions at the N-terminus of the domain |

Note: †, The MAT1-1-1 transcripts GAGW01008880 and OSIN7648 were derived from the *C. sinensis* insect-fungi complexes. The arrows “→” and “←” indicate sequences in the sense and antisense strands of the genomes, respectively; “&” refers to the removed intron portion.

### Table S7. Summary of the Bayesian clustering results in Figure 4 with the amino acid substitutions in the HMG-box_ROX1-like domains of the 25 full-length MAT1-2-1 proteins of wild-type *C. sinensis* isolates under the AlphaFold UniProt codes and GenBank accession numbers.

| ***C. sinensis* isolate** | **GenBank accession #** | **AlphaFold UniProt code** | **The Bayesian Cluster based on** | | **Amino acid substitution in the HMG-box_ROX1-like domain** **(*vs*. AEH27625)** |
| --- | --- | --- | --- | --- | --- |
|  |  |  | **full length sequence*** | **HMG-box_ROX1-like domain sequence** |  |
| XZ12_16 | AIV43040 | A0A0A0RCF5 | II-1 | b2β | V-to-H, M-to-I, and Q-to-R |
| XZ-LZ07-H1 XZ-LZ07-H2 XZ06-124 XZ05_8 | ACV60417 ACV60418 AFH35020 AFX66443 | D7F2J7 | II-2 | b2α | V-to-H and M-to-I |
| XZ-SN-44 XZ-LZ05-6 XZ05_2 XZ06_260 XZ09_80 | ACV60375 ACV60415 AFX66441 AFX66446 AFX66461 | D7F2F5 | III | b2α | V-to-H and M-to-I |
| XZ-NQ-154 XZ-NQ-155 GS09_111 QH09-93 CS560-961 | ACV60363 ACV60364 AFX66388 AFH35018 AGW27542 | D7F2E3 | V-1 | b1α | V-to-H |
| QH-YS-199 | ACV60385 | D7F2G5 | V-2 | b1α | V-to-H |
| QH09_11 | AFX66401 | V9LW71 | V-2 | b1α | V-to-H |
| YN09_6 YN09_22 YN09_51 | AFX66472 AFX66473 AFX66474 | V9LVS8 | IV-2 | b1α | V-to-H |
| YN09_61 | AFX66475 | V9LVU8 | V-2 | b1α | V-to-H |
| YN09_64 | AFX66476 | V9LWC9 | IV-1 | b1α | V-to-H |
| ID10_1 | AFX66484 | V9LWG5 | V-2 | b1α | V-to-H |
| CS6-251 | AGW27537 | U3N6V5 | V-2 | b1α | V-to-H |
| SC-3 | ACV60399 | D7F2H9 | V-2 | b1β | V-to-H and Q-to-R |

Note: *, The Bayesian clustering results were reported by Li *et al*. [2025].

### Table S8. Summary of the Bayesian clustering results in Figure 4 with the amino acid substitutions and deletions in the HMG-box_ROX1-like domains of the MAT1-2-1 proteins encoded by the genome and transcriptome assemblies of *H. sinensis* strains and metatranscriptome assembly of natural *C. sinensis* insect-fungi complexes under the GenBank accession numbers.

| ***H. sinensis* strain** | **GenBank accession #** | **The HMG-box_ROX1-like domain** | | |
| --- | --- | --- | --- | --- |
|  |  | **Nucleotide sequence arrange (deleted intron region)** | **Bayesian cluster** | **Amino acid substitution (*vs*. AEH27625)** |
| Co18 | ANOV01000063 | 9759→9851 & 9907→10,026 | a2 | S-to-A |
| 1229 | LKHE01001605 | 14,016←14,135 & 14,191←14,283 | b3 | Y-to-H and S-to-A |
| ZJB12195 | LWBQ01000021 | 239,029←239,148 & 239,204←239,269 | b3 | Y-to-H and S-to-A |
| CC1406-20395 | NGJJ01000619 | 23,186←23,305 & 23,361←23,453 | b3 | Y-to-H and S-to-A |
| L0106 | GCQL01020543 | 553←765 | b1α | Y-to-H |
| † | OSIN7649 | 379→591 | b1α | (100% identical) |

Note: †, The MAT1-2-1 transcript OSIN7649 was derived from the mature *C. sinensis* insect-fungi complex. The arrows “→” and “←” indicate sequences in the sense and antisense strands of the genomes, respectively; “&” refers to the removed intron portion.

### [REFERENCES]

1. Abramson J, Adler J, Dunger J, Evans R, Green T, Pritzel A, Ronneberger O, Willmore L, Ballard AJ, Bambrick J, Bodenstein SW, Evans DA, Hung CC, O'Neill M, Reiman D, Tunyasuvunakool K, Wu Z, Žemgulytė A, Arvaniti E, Beattie C, Bertolli O, Bridgland A, Cherepanov A, Congreve M, Cowen-Rivers AI, Cowie A, Figurnov M, Fuchs FB, Gladman H, Jain R, Khan YA, Low CMR, Perlin K, Potapenko A, Savy P, Singh S, Stecula A, Thillaisundaram A, Tong C, Yakneen S, Zhong ED, Zielinski M, Žídek A, Bapst V, Kohli P, Jaderberg M, Hassabis D, Jumper JM. Accurate structure prediction of biomolecular interactions with AlphaFold 3. Nature. 2024; 630(8016): 493−500. [doi: 10.1038/s41586-024-07487-w](doi:%2010.1038/s41586-024-07487-w)
2. Asante-Owusu RN, Banham AH, Böhnert HU, Mellor EJC, Casselton LA. Heterodimerization between two classes of homeodomain proteins in the mushroom *Coprinus cinereus* brings together potential DNA-binding and activation domains. [Gene](https://www.sciencedirect.com/journal/gene) 1996; 172(1): 25−31. <https://doi.org/10.1016/0378-1119(96)00177-1>
3. Balasubramanian B, Lowry CV, Zitomer RS. The Rox1 repressor of the *Saccharomyces cerevisiae* hypoxic genes is a specific DNA-binding protein with a high-mobility-group motif. Mol Cell Biol. 1993; 13(10): 6071−6078. [doi: 10.1128/mcb.13.10.6071-6078.1993](doi:%2010.1128/mcb.13.10.6071-6078.1993)
4. Barseghyan GS, Holliday JC, Price TC, Madison LM, Wasser SP. Growth and cultural-morphological characteristics of vegetative mycelia of medicinal caterpillar fungus *Ophiocordyceps sinensis* G.H. Sung *et al*. (Ascomycetes) Isolates from Tibetan Plateau (P. R. China). Intl. J. Med. Mushrooms. 2011; 13(6): 565−581. DOI: [10.1615/intjmedmushr.v13.i6.90](https://doi.org/10.1615/intjmedmushr.v13.i6.90)
5. [Baxevanis AD,](https://www.zhangqiaokeyan.com/search.html?doctypes=4_5_6_1-0_4-0_1_2_3_7_9&sertext=Baxevanis%20AD&option=202) [Bryant SH,](https://www.zhangqiaokeyan.com/search.html?doctypes=4_5_6_1-0_4-0_1_2_3_7_9&sertext=Bryant%20SH&option=202) [Landsman D](https://www.zhangqiaokeyan.com/search.html?doctypes=4_5_6_1-0_4-0_1_2_3_7_9&sertext=Landsman%20D&option=202). Homology model building of the HMG-1 box structural domain. Nucleic Acids Research 1995; 23(6): 1019−1029. DOI: [10.1093/nar/23.6.1019](https://doi.org/10.1093/nar/23.6.1019)
6. Ait Benkhali J, Coppin E, Brun S, Peraza-Reyes L, Martin T, Dixelius C, Lazar N, van Tilbeurgh H, Debuchy R. A Network of HMG-box Transcription Factors Regulates Sexual Cycle in the Fungus Podospora anserina. PLoS Genet 2013; 9(7): e1003642. <doi:10.1371/journal.pgen.1003642>
7. Bennett RJ, Johnson AD. Completion of a parasexual cycle in *Candida albicans* by induced chromosome loss in tetraploid strains. EMBO J. 2003; 22(10): 2505−2515. [DOI: 10.1093/emboj/cdg235](DOI:%2010.1093/emboj/cdg235)
8. Bushley KE, Li Y, Wang W-J, Wang X-L, Jiao L, Spatafora JW, Yao Y-J. Isolation of the MAT1-1 mating type idiomorph and evidence for selfing in the Chinese medicinal fungus *Ophiocordyceps sinensis*. Fungal Biol. 2013; 117(9): 599−610. [DOI: 10.1016/j.funbio.2013.06.001](DOI:%2010.1016/j.funbio.2013.06.001)
9. Chen C-S, Hseu R-S, Huang C-T. Quality control of *Cordyceps sinensis* teleomorph, anamorph, and Its products. Chapter 12, in (Shoyama, Y., Ed.) Quality Control of Herbal Medicines and Related Areas. InTech, Rijeka, Croatia (2011). pp. 223–238. www.intechopen.com
10. Chen Y-Q, Hu B, Xu F, Zhang W, Zhou H, Qu L-H. Genetic variation of *Cordyceps sinensis*, a fruit-body-producing entomopathogenic species from different geographical regions in China. FEMS Microbiol Lett. 2004; 230: 153–158. [Doi: 10.1016/S0378-1097(03)00889-9](Doi:%2010.1016/S0378-1097(03)00889-9) (accessed on 30 January 2025).
11. China Ministry of Agriculture and Rural Affairs. Announcement (No. 15 of 2021) of National Forestry and Grassland Administration: List of National Key Protected Wild Plants. September 7, 2021. Available online: [https://m.163.com/dy/article/HHCVOJPU055360T7.html](https://m.163.com/dy/article/HHCVOJPU055360T7.html%20) (accessed on 3 May 2025).
12. Dai R-Q, Lan J-L, Chen W-H, Li X-M, Chen Q-T, Shen C-Y. Discovery of a new fungus *Paecilomyces hepiali* Chen & Dai. Acta Agricult. Univ. Pekin. 1989; 15(2): 221−224.
13. David A, Islam S, Tankhilevich E, Sternberg MJE. The AlphaFold Database of Protein Structures: A Biologist’s Guide. J Mol Biol. 2022; 434(2): 167336. <https://doi.org/10.1016/j.jmb.2021.167336>
14. Debuchy R, Turgeo BG. Mating-Type Structure, Evolution, and Function in Euascomycetes. In (eds. Kües U. & Fischer, R.) Growth, Differentiation and Sexuality. Springer (2006). pp. 293–323.
15. Deleage G, Roux B. An algorithm for protein secondary structure prediction based on class prediction. Protein Engineering, Design and Selection 1987; 1: 289–294. <https://doi.org/10.1093/protein/1.4.289>
16. Du X-H, Wu D-M, Kang H, Wang H-C, Xu N, Li T-T, Chen K-L. Heterothallism and potential hybridization events inferred for twenty-two yellow morel species. IMA Fungus. 2020; 11: 4. <https://doi.org/10.1186/s43008-020-0027-1>
17. Gao L, Li X-H, Zhao J-Q, Lu J-H, Zhao J-G, Zhu J-S. Maturation of *Cordyceps sinensis* associates with alterations of fungal expressions of multiple *Ophiocordyceps sinensis* mutants with transition and transversion point mutations in stroma of *Cordyceps sinensis*. Beijing Da Xue Xue Bao. 2012; 44(3): 454–463. [doi: 10.3969/j.issn.1671-167X.2012.03.025](doi:%2010.3969/j.issn.1671-167X.2012.03.025)
18. Gasteiger E, Hoogland C, Gattiker A, Duvaud S, Wilkins MR, Appel RD, Bairoch A. Protein Identification and Analysis Tools on the ExPASy Server. Chapter 52 (In) John M. Walker (ed): The Proteomics Protocols Handbook, Humana Press (2005). pp. 571–607.
19. Guo M-Y, Liu Y, Gao Y-H, Jin T, Zhang H-B, Zhou X-W. Identification and bioactive potential of endogenetic fungi isolated from medicinal caterpillar fungus *Ophiocordyceps sinensis* from Tibetan Plateau. Int J Agric Biol. 2017; 19: 307‒313. [DOI: 10.17957/IJAB/15.0281](DOI:%2010.17957/IJAB/15.0281)
20. Hancock SP, Cascio D, Johnson RC. Cooperative DNA binding by proteins through DNA shape complementarity. Nucleic Acids Res. 2019; 47(16):8874‒8887. [doi: 10.1093/nar/gkz642](doi:%2010.1093/nar/gkz642)
21. Hawksworth DL, Crous PW, Redhead SA, Reynolds DR, Samson RA, Seifert KA, and 82 other authors. The Amsterdam declaration on fungal nomenclature. IMA Fungus. 2011; 2: 105−112. [DOI: 10.5598/imafungus.2011.02.01.14](DOI:%2010.5598/imafungus.2011.02.01.14)
22. Hėnault M, Marsit S, Charron G, Landry CR. The effect of hybridization on transposable element accumulation in an undomesticated fungal species. eLife. 2020; 9: e60474. [DOI: https://doi.org/10.7554/eLife.60474](DOI:%20https://doi.org/10.7554/eLife.60474)
23. Holliday J, Cleaver M. Medicinal value of the caterpillar fungi species of the genus Cordyceps (Fr.) Link (Ascomycetes). A review. Int. J. Med. Mushrooms. 2008; 10: 219–234. [DOI: 10.1615/IntJMedMushr.v10.i3.30](DOI:%2010.1615/IntJMedMushr.v10.i3.30)
24. Hu X, Zhang Y-J, Xiao G-H, Zheng P, Xia Y-L, Zhang X-Y, St Leger RJ, Liu X-Z, Wang C-S. Genome survey uncovers the secrets of sex and lifestyle in caterpillar fungus. Chin. Sci. Bull. 2013; 58: 2846–2854. [doi: 10.1007/s11434-013-5929-5](doi:%2010.1007/s11434-013-5929-5)
25. Huelsenbeck JP, Ronquist F. MRBAYES: Bayesian inference of phylogeny. Bioinformat. 2001; 17: 754−755. [DOI: 10.1093/bioinformatics/17.8.754](DOI:%2010.1093/bioinformatics/17.8.754)
26. Jackson D, Lawson T, Villafane R, Gary L. Modeling the structure of yeast MATα1: An HMG-Box motif with a C-terminal helical extension. Open J Biophysics, 2013, 3: 1−12. <http://dx.doi.org/10.4236/ojbiphy.2013.31001>
27. Jacobsen S, Wittig M, Pöggeler S. Interaction Between Mating-Type Proteins From the Homothallic Fungus Sordaria macrospora. Curr. Genet. 2002; 41: 150–158. [doi: 10.1007/s00294-002-0276-0](doi:%2010.1007/s00294-002-0276-0)
28. Jiang Y, Yao Y-J. A review for the debating studies on the anamorph of *Cordyceps sinensis*. Mycosistema. 2003; 22(1): 161–176.
29. Jin L‑Q, Xu Z‑W, Zhang B, Yi M, Weng C‑Y, Lin S, Wu H, Qin X-T, Xu F, Teng Y, Yuan S-J, Liu Z-Q, Zheng Y-G. Genome sequencing and analysis of fungus *Hirsutella sinensis* isolated from *Ophiocordyceps sinensis*. AMB Expr. 2020; 10: 105. <https://doi.org/10.1186/s13568-020-01039-x>
30. Jones SK, Bennett RJ. Fungal mating pheromones: choreographing the dating game. Fungal Genet. Biol. 2011; 48(7): 668–676. [doi: 10.1016/j.fgb.2011.04.001](doi:%2010.1016/j.fgb.2011.04.001)
31. Jumper J, Evans R, Pritzel A, Green T, Figurnov M, Ronneberger O, Tunyasuvunakool K, Bates R, Žídek A, Potapenko A, Bridgland A, Meyer C, Kohl SAA, Ballard AJ, Cowie A, Romera-Paredes B, Nikolov S, Jain R, Adler J, Back T, Petersen S, Reiman D, Clancy E, Zielinski M, Steinegger M, Pacholska M, Berghammer T, Bodenstein S, Silver D, Vinyals O, Senior AW, Kavukcuoglu K, Kohli P, Hassabis D. Highly accurate protein structure prediction with AlphaFold. Nature 2021; 596: 583–589. <https://doi.org/10.1038/s41586-021-03819-2>
32. Kang Q, Zhang J, Chen F, Dong C, Qin Q, Li X, Wang H, Zhang H and Meng Q. Unveiling mycoviral diversity in *Ophiocordyceps sinensis* through transcriptome analyses. Front. Microbiol. 2024; 15: 1493365. [doi: 10.3389/fmicb.2024.1493365](doi:%2010.3389/fmicb.2024.1493365)
33. Kastaniotis AJ, Zitomer RS. Rox1 mediated repression. Oxygen dependent repression in yeast. Adv Exp Med Biol. 2000; 475: 185–195. <https://pubmed.ncbi.nlm.nih.gov/10849660/>
34. Kim H-K, Jo S-M, Kim G-Y, Kim D-W, Kim Y-K, Yun S-H. A large-scale functional analysis of putative target genes of mating-type loci provides insight into the regulation of sexual development of the cereal pathogen *Fusarium graminearum*. PLoS Genet 2015; 11(9): e1005486. <doi:10.1371/journal.pgen.1005486>
35. Kinjo N, Zang M. Morphological and phylogenetic studies on *Cordyceps sinensis* distributed in southwestern China. Mycoscience. 2001; 42: 567–574. [DOI: 10.1007/BF02460956](DOI:%2010.1007/BF02460956)
36. Kück U, Bennett RJ, Wang L and Dyer PS. Editorial: Sexual and Parasexual Reproduction of Human Fungal Pathogens. Front. Cell. Infect. Microbiol. 2022; 12: 934267. [doi: 10.3389/fcimb.2022.934267](doi:%2010.3389/fcimb.2022.934267)

88. [Thapar](https://pubmed.ncbi.nlm.nih.gov/?term=Thapar+R&cauthor_id=25748361) R. Structure-specific nucleic acid recognition by L-motifs and their diverse roles in expression and regulation of the genome. Biochim Biophys Acta 2015; 1849(6): 677–687. [doi: 10.1016/j.bbagrm.2015.02.006](doi:%2010.1016/j.bbagrm.2015.02.006)
89. Tunyasuvunakool, K., Adler, J., Wu, Z, Green T, Zielinski M, Žídek A, Bridgland A, Cowie A, Meyer C, Laydon A, Velankar S, Kleywegt GJ, Bateman A, Evans R, Pritzel A, Figurnov M, Ronneberger O, Bates R, Kohl SAA, Potapenko A, Ballard AJ, Romera-Paredes B, Nikolov S, Jain R, Clancy E, Reiman D, Petersen S, Senior AW, Kavukcuoglu K, Birney E, Kohli P, Jumper J, Hassabis D. Highly accurate protein structure prediction for the human proteome. Nature 2021; 596: 590–596. <https://doi.org/10.1038/s41586-021-03828-1>
90. Turgeon BG, Yoder OC. Proposed nomenclature for mating type genes of filamentous ascomycetes. Fungal Genet. Biol. 2000; 31: 1‒5. DOI: [10.1006/fgbi.2000.1227](https://doi.org/10.1006/fgbi.2000.1227)
91. Varadi M, Bertoni D, Magana P, Paramval U, Pidruchna I, Radhakrishnan M, Tsenkov M, Nair S, Mirdita M, Yeo J, Kovalevskiy O, Tunyasuvunakool K, Laydon A, Žídek A, Tomlinson H, Hariharan D, Abrahamson J, Green T, Jumper J, Birney E, Steinegger M, Hassabis D, Velankar S. AlphaFold Protein Structure Database in 2024: providing structure coverage for over 214 million protein sequences. Nucleic Acids Res. 2024; 52(D1): D368–D375. [doi: 10.1093/nar/gkad1011](doi:%2010.1093/nar/gkad1011)
92. Wang Y, Stata M, Wang W, Stajich JE, White MM, Moncalvo JM. Comparative genomics reveals the core gene toolbox for the fungus-insect symbiosis. mBio. 2018; 9: e00636-18. [DOI: 10.1128/mBio.00636-18](DOI:%2010.1128/mBio.00636-18)
93. Wei J-C, Wei X-L, Zheng W-F, Guo W, Liu R-D. Species identification and component detection of *Ophiocordyceps sinensis* cultivated by modern industry. Mycosystema. 2016; 35(4): 404‒410.
94. Wei X-L, Yin X-C, Guo Y-L, Shen N-Y, Wei, J.-C. Analyses of molecular systematics on *Cordyceps sinensis* and its related taxa. Mycosystema. 2006; 25(2): 192–202.
95. Wilson AM, Wilken PM, van der Nest MA, Steenkamp ET, Wingfield MJ, Wingfield BD. Homothallism: an umbrella term for describing diverse sexual behaviours. IMA Fungus. 2015; 6(1): 207–214. DOI: [10.5598/imafungus.2015.06.01.13](https://doi.org/10.5598/imafungus.2015.06.01.13)
96. Wroblewski K, Kmiecik S. Integrating AlphaFold pLDDT Scores into CABS-flex for enhanced protein flexibility simulations. Comput Struct Biotechnol J. 2024; 30(23): 4350–4356. [doi: 10.1016/j.csbj.2024.11.047](doi:%2010.1016/j.csbj.2024.11.047)
97. Xia E-H, Yang D-R, Jiang J-J, Zhang Q-J, Liu Y, Liu Y-L, Zhang Y, Zhang H-B, Shi C, Tong Y, Kim C-H, Chen H, Peng Y-Q, Yu Y, Zhang W, Eichler EE, Gao L-Z. The caterpillar fungus, *Ophiocordyceps sinensis*, genome provides insights into highland adaptation of fungal pathogenicity. Sci. Rep. 2017; 7: 1806. <DOI:10.1038/s41598-017-01869-z>
98. Xia F, Liu Y, Shen G-L, Guo L-X, Zhou X-W. Investigation and analysis of microbiological communities in natural *Ophiocordyceps sinensis*. Can. J. Microbiol. 2015; 61: 104‒111. DOI: [10.1139/cjm-2014-0610](https://doi.org/10.1139/cjm-2014-0610)
99. Xiang L, Li Y, Zhu Y, Luo H, Li C, Xu X, Sun C, Song J-Y, Shi L-H, He L, Sun W, [Chen](https://www.sciencedirect.com/author/35483515200/shilin-chen) S-L. Transcriptome analysis of the *Ophiocordyceps sinensis* fruiting body reveals putative genes involved in fruiting body development and cordycepin biosynthesis. Genomics. 2014; 103: 154−159. <https://doi.org/10.1016/j.ygeno.2014.01.002>
100. Xiao W, Yang J-P, Zhu P, Cheng K-D, He H-X, Zhu H-X, Wang Q. Non-support of species complex hypothesis of *Cordyceps sinensis* by targeted rDNA-ITS sequence analysis. Mycosystema. 2009; 28(6): 724–730.
101. Xu T, Xu Q, Li J-Y. Toward the appropriate interpretation of Alphafold2. Front Artif Intell. 2023; 6: 1149748. [doi: 10.3389/frai.2023.1149748](doi:%2010.3389/frai.2023.1149748)
102. Yamamoto A, Ando Y, Yoshioka K, Saito K, Tanabe T, Shirakawa H, Yoshida M. Difference in affinity for DNA between HMG proteins 1 and 2 determined by surface plasmon resonance measurements. J Biochem. 1997;122(3): 586‒594. [doi: 10.1093/oxfordjournals.jbchem.a021793. PMID: 9348088](doi:%2010.1093/oxfordjournals.jbchem.a021793.%20PMID:%209348088)
103. Yang J-L, Xiao W, He H-X, Zhu H-X, Wang S-F, Cheng -KD, Zhu P. Molecular phylogenetic analysis of *Paecilomyces hepiali* and *Cordyceps sinensis*. Acta Pharmaceut. Sinica 2008; 43(4): 421‒426.
104. Yang J-Y, Tong X-X, He C-Y, Bai J, Wang F, Guo J-L. Comparison of endogenetic microbial community diversity between wild *Cordyceps sinensis*, artificial *C. sinensis* and habitat soil. Chin. J. Chin. Materia Medica. 2021; 46(12): 3106‒3115.
105. Yao Y-S, Zhu J-S. Indiscriminate use of the Latin name for natural *Cordyceps sinensis* and *Ophiocordyceps sinensis* fungi. Chin. J. Chin. Mater. Med. 2016; 41(7): 1316–1366.
106. Zhang S; Zhang Y-J. Molecular evolution of three protein-coding genes in the Chinese caterpillar fungus *Ophiocordyceps sinensis*. Microbiol. China. 2015; 42(8): 1549−1560.
107. Zhang S, Zhang Y-J, Liu X-Z, Wen H-A, Wang M, Liu D-S. Cloning and analysis of the MAT1-2-1 gene from the traditional Chinese medicinal fungus *Ophiocordyceps sinensis*. Fungal Biol. 2011; 115: 708−714.
108. Zhang S, Zhang Y-J, Shrestha B, Xu J-P, Wang C-S, Liu X-Z. *Ophiocordyceps sinensis* and *Cordyceps militaris*: research advances, issues and perspectives. Mycosystema. 2013; 32: 577−597.
109. Zhang S-W, Cen K, Liu Y, Zhou X-W, Wang C-S. Metatranscriptomics analysis of the fruiting caterpillar fungus collected from the Qinghai-Tibetan plateau. Sci. Sinica Vitae. 2018; 48(5): 562−570.
110. Zhang Y-J, Li E-W, Wang C-S, Li Y-L, Liu X-Z. *Ophiocordyceps sinensis*, the flagship fungus of China: terminology, life strategy and ecology. Mycol. 2012; 3(1): 2–10. <https://www.tandfonline.com/doi/full/10.1080/21501203.2011.654354>
111. Zhang Y-J, Sun B-D, Zhang S, Wàngmŭ, Liu X-Z, Gong W-F. Mycobiotal investigation of natural *Ophiocordyceps sinensis* based on culture-dependent investigation. Mycosistema. 2010; 29(4): 518–527.
112. Zhang Y-J, Xu L-L, Zhang S, Liu X-Z, An Z-Q, Wàngmŭ, Guo Y-L. Genetic diversity of *Ophiocordyceps sinensis*, a medicinal fungus endemic to the Tibetan Plateau: implications for its evolution and conservation. BMC Evol. Biol. 2009; 9: 290. <doi:10.1186/1471-2148-9-290>
113. Zhang Y-J, Zhang S, Li Y-L, Ma S-L, Wang C-S, Xiang M-C, Liu X, An Z-Q, Xu J-P, Liu X-Z. Phylogeography and evolution of a fungal–insect association on the Tibetan Plateau. Mol. Ecol. 2014; 23: 5337−5355. DOI: [10.1111/mec.12940](https://doi.org/10.1111/mec.12940)
114. Zheng, P.; Wang, C.-S. Sexuality Control and Sex Evolution in Fungi. Sci. Sin. Vitae 2013, 43, 1090–1097.
115. Zheng Q, Hou R, Zhang J-Y, Ma J, Ma J-W, Wu Z-S, Wang G-H, Wang C-F, Xu J-R. The MAT locus genes play different roles in sexual reproduction and pathogenesis in *Fusarium graminearum*. PLoS ONE 2013; 8(6): e66980. <doi:10.1371/journal.pone.0066980>
116. Zhong X, Gu L, Wang H-Z, Lian D-H, Zheng Y-M, Zhou S, Zhou W, Gu J, Zhang G, Liu X. Profile of *Ophiocordyceps sinensis* transcriptome and differentially expressed genes in three different mycelia, sclerotium and fruiting body developmental stages. Fungal Biol. 2018; 122: 943‒951. DOI: [10.1016/j.funbio.2018.05.011](https://doi.org/10.1016/j.funbio.2018.05.011)
117. Zhou XW, Li LJ, Tian EW. Advances in research of the artificial cultivation of *Ophiocordyceps sinensis* in China. *Critical* Rev Biotechnol 2013; *34*(3), 233–243. <https://doi.org/10.3109/07388551.2013.791245>
118. Zhu J-S, Gao L, Li X-H, Yao Y-S, Zhou Y-J, Zhao J-Q, Zhou Y-J. Maturational alterations of oppositely orientated rDNA and differential proliferations of CG:AT-biased genotypes of *Cordyceps sinensis* fungi and *Paecilomyces hepiali* in natural *C. sinensis*. Am. J. Biomed. Sci. 2010; 2(3): 217–238. [doi: 10.5099/aj100300217](doi:%2010.5099/aj100300217)
119. Zhu J-S, Guo Y-L, Yao Y-S, Zhou Y-J, Lu J-H, Qi Y, Chen W, Zheng T-Y, Zhang L, Wu Z-M, Zhang L-J, Liu X-J, Yin W-T. Maturation of *Cordyceps sinensis* associates with co-existence of *Hirsutella sinensis* and *Paecilomyces hepiali* DNA and dynamic changes in fungal competitive proliferation predominance and chemical profiles. J. Fungal Res. 2007; 5(4): 214–224.
120. Zhu J-S, Halpern GM, Jones K. The scientific rediscovery of a precious ancient Chinese herbal regimen: *Cordyceps sinensis*: Part I. J. Altern. Complem. Med. 1998a; 4(3): 289–303. DOI: [10.1089/acm.1998.4.3-289](https://doi.org/10.1089/acm.1998.4.3-289)
121. Zhu J-S, Halpern GM, Jones K. The scientific rediscovery of an ancient Chinese herbal medicine: *Cordyceps sinensis*: Part II. J. Altern. Complem. Med. 1998b; 4(4): 429–457. DOI: [10.1089/acm.1998.4.429](https://doi.org/10.1089/acm.1998.4.429)
122. Zhu J-S, Li C-L, Tan N-Z, Berger JL, Prolla TA. Combined use of whole-gene expression profiling technology and mouse lifespan test in anti-aging herbal product study. Proc. 2011 New TCM Products Innovation and Industrial Development Summit, Hangzhou, China (Nov 27, 2011). pp. 443–448. Available online: <https://xueshu.baidu.com/usercenter/paper/show?paperid=08341c17fa58c8f85584b92572b90f75&site=xueshu_se> (accessed on 30 Jannuary 2025)
123. Zhu J-S, Li Y-L. A Precious Transitional Chinese Medicine, *Cordyceps sinensis*: Multiple heterogeneous *Ophiocordyceps sinensis* in the insect-fungi complex. Lambert Academic Publishing, Saarbrüchen. Germany, 2017.
124. Zitomer RS, Limbach MP, Rodriguez-Torres AM, Balasubramanian B, Deckert J, Snow PM. Approaches to the study of Rox1 repression of the hypoxic genes in the yeast Saccharomyces cerevisiae. Methods. 1997; 11(3): 279–288. [doi: 10.1006/meth.1996.0422](doi:%2010.1006/meth.1996.0422)
